# Biodistribution-Driven Discovery Identifies a Glycosidase-Cleavable Linker to Reprogram Radiotheranostics

**DOI:** 10.1101/2025.09.11.675134

**Authors:** Woonghee Lee, Sai Reddy Doda, Kwamena E. Baidoo, Divya Nambiar, Joon-Yong Chung, Stephen Adler, Elijah Edmondson, Yuki Ueda, Satoshi Omiya, Xiaoyi Li, Hima Makala, Julia Sheehan-Klenk, Stanley Fayn, Orit Jacobson Weiss, Eric Lindberg, Jessica A. Beck, Amy K. LeBlanc, Frank I. Lin, Peter L. Choyke, Rolf Swenson, Martin J. Schnermann, Freddy E. Escorcia

## Abstract

While radiopharmaceutical therapy (RPT) has become part of the standard-of-care for patients with advanced prostate cancers and neuroendocrine tumors (NETs), cures are elusive and normal tissue toxicity remain a challenge. Chemical groups susceptible to cleavage by enzymes present in tumors, tumor microenvironment or in normal tissues, have the potential to improve the therapeutic index for RPT. Using DOTA-TATE as an example and drawing from strategies used to develop antibody-drug conjugates, we designed, and synthesized, a chemically diverse series of linkers between the chelator (DOTA) and the targeting vector (TATE). Of the 10 agents we tested, two with cleavable linker domains reduced kidney retention compared to DOTA-TATE: the previously reported DOTA-MVK(ε)-TATE, and a novel agent bearing cleavable beta-galactose (β-Gal) unit, DOTA-β-Gal-TATE. In murine models of NETs, positron emission tomography (PET) was used to image yttrium-86 (^86^Y)-labeled variants and show that, while the ^86^Y-DOTA-MVK(ε)-TATE exhibits similar tumor uptake to the parent non-cleavable ^86^Y-DOTA-TATE, ^86^Y-DOTA-β-Gal-TATE shows enhanced tumor uptake, resulting in up to 10-fold improvement in the tumor-to-kidney ratios compared to ^86^Y-DOTA-TATE. *In vitro* and *in vivo* studies confirm high efficiency, enzyme-specific cleavage of ^86^Y-DOTA-MVK(ε)-TATE and ^86^Y-DOTA-β-Gal-TATE, supporting a key role for cleavable linker chemistry in the observed outcomes. RPT studies using actinium-225 (^225^Ac)-labeled variants confirm that all agents are therapeutically effective and well tolerated. While both cleavable variants exhibit superior local control, overall survival, and more favorable toxicity profile when compared with ^225^Ac-DOTA-TATE, ^225^Ac-DOTA-β-Gal-TATE demonstrated lower nephrotoxicity. Our findings suggest a potentially generalizable strategy for improving the pharmacokinetics of radiopharmaceutical therapy agents.

**One Sentence Summary:** A β-galactose-cleavable linker reduces kidney toxicity, enhances tumor targeting and therapeutic efficacy in radiopharmaceutical therapy.

## INTRODUCTION

Radiopharmaceuticals, which can be used for therapeutic and diagnostic purposes, are often referred to as “radiotheranostics.” This approach has been in use since the advent of radioiodine in the 1940s, and, more recently, new paired targeted radiotheranostic agents have been clinically approved for several malignancies and many more are being tested in clinical trials*(1)*.

Among the most successful products are the positron emission tomography (PET) agents ^68^Ga-DOTA-TATE and ^64^Cu-DOTA-TATE, along with the therapeutic β particle-emitting ^177^Lu-DOTA-TATE, which are approved for imaging and treatment of somatostatin receptor (SSTR)-expressing neuroendocrine tumors (NETs)*(2)*, respectively. These peptide-based radiopharmaceuticals have revolutionized patient management for this disease. However, dose-limiting kidney toxicity remains a significant challenge. As part of standard care, patients receiving therapeutic ^177^Lu-DOTA-TATE must receive infusions of positively charged amino acids to reduce kidney accumulation of the radioconjugate and mitigate nephrotoxicity. Strategies that enhance tumor retention while minimizing off-target uptake, particularly in the kidneys, could substantially improve the therapeutic index of SSTR-targeted radiopharmaceutical therapy (RPT), and, perhaps, that of other renally cleared agents.

Targeted radiopharmaceuticals are typically comprised of a chelator (e.g. DOTA) that can form stable complexes with radiometals, the targeting vector (e.g. octreotate, or TATE) and a chemical linker that connects the two chemical moieties. A cleavable linker has the potential to tune the off-target, and, potentially, tumor-targeting properties of these agents. Prior efforts to develop cleavable linkers focused on peptide sequences that were designed to reduce kidney reuptake*(3, 4)*. Separately, there have been extensive studies to develop and optimize antibody-drug conjugates (ADCs) – agents that also comprise a targeting component, linker and cytotoxic payloads*(5)*. Many linkers have been examined with improved *in vivo* performance*(6, 7)*. These ADC-optimized linkers may have potential in radiopharmaceutical applications.

Here, we report a systematic exploration of a small library of chemically diverse DOTA-*linker*-TATE constructs, each bearing unique linkers that confer distinct properties (e.g. cleavable, albumin binding, etc). By applying a biodistribution-first approach and then PET imaging, we identify a β-galactose DOTA-TATE ligand, DOTA-β-Gal-TATE, that exhibits both superior tumor binding and significantly lower kidney retention in murine models of NETs when compared to parental DOTA-TATE and a previously reported DOTA-MVK(ε)-TATE renal brush border-cleavable variant. Finally, we also confirm that alpha particle therapy with ^225^Ac-DOTA-β-Gal-TATE of mice bearing NETs shows superior local control and overall survival, and improved toxicity profile compared with animals treated with parental ^225^Ac-DOTA-TATE. ^225^

Ac-DOTA-MVK(ε)-TATE achieved comparable local control and overall survival but exhibited greater nephrotoxicity, underscoring the advantage of ^225^Ac-DOTA-β-Gal-TATE. The work presented here has implications for other radiopharmaceuticals, peptide-based and beyond.

## RESULTS

### Design and synthesis of DOTA-TATE variants with unique linker properties

In the design of the panel of DOTA-*linker*-TATE constructs, we set out to compare strategies with the potential to alter *in vivo* biodistribution. We took note of two peptide-based linkers, methionine-valine-lysine (MVK) and glycyl-tyrosine (GY)*(8–11)*, which are cleavable by groups of proteases in the luminal space of the proximal tubule that facilitate cleavage and excretion prior to reabsorption*(1)*. We synthesized DOTA-MVK-, MVK(ε)-, GY*- and GY-TATE derivatives (**Fig. 1**) using established solid-phase peptide synthesis methods (**Fig S1-11**). Of these, MVK and GY* are designed to be non-cleavable isomeric controls, while GY and MVK(ε) should be cleavable variants. The important role of charge in reabsorption has been previously reported, with anionic charge being shown to reducing renal uptake and retention of other peptidic agents*(12)*. Accordingly, we prepared a tri-anionic a DOTA-GDDDG-TATE derivative. Additionally, numerous efforts have found significant impact of albumin-binding domains*(13–15)*, leading us to a sterol linker DOTA-Chol-TATE. Finally, building on studies from ADC linker efforts, we accessed a panel of glycans that are cleavable by naturally occurring glycosidases in renal proximal tubules. These include DOTA-linker-TATE variants of β-galactose (β-Gal), β-glucose (β-Glu), and β-glucuronide (β-GlcA). To our knowledge, such cleavable-glycan linkers have not been previously tested in the context of SSTR targeting or for broader radiopharmaceutical development purposes.

**Fig. 1.**
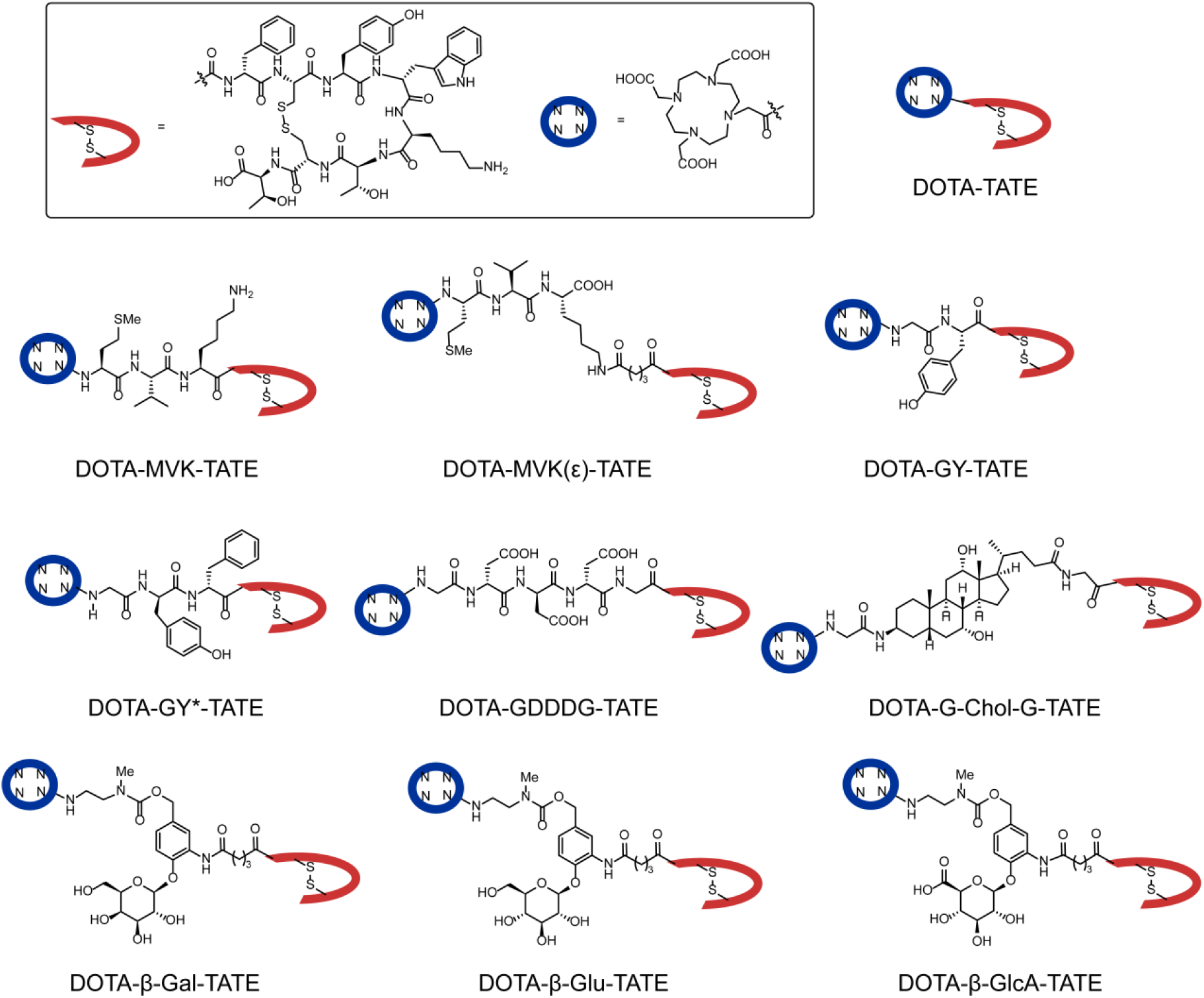
Structure of DOTA-*linker-*TATE-derivatives.

### DOTA-MVK(ε)-TATE and DOTA-β-Gal-TATE exhibit lower kidney retention compared with DOTA-TATE in non-tumor-bearing animals

The panel of 10 compounds were then tested in an initial biodistribution screen of 9 organs assessed in non-tumor-bearing mice at specific timepoints following administration of the radioconjugates. Each agent was initially labeled with ^111^In, a long-lived gamma-emitting radioisotope suitable for biodistribution studies, with high radiochemical yields and purity (**Fig. S12**). The compounds were then dosed (1.55 ± 0.12 MBq; 1 µg per mouse) in healthy athymic nude mice via intravenous (i.v.) injection in phosphate-buffered saline (PBS; vehicle). After 2 hours, the mice were sacrificed and *ex vivo* biodistribution analysis was performed (**Fig. 2**). This initial screen showed substantial variation in renal uptake among the compounds (**Fig. 2A**). The synthesized agents were compared to parental DOTA-TATE (**Fig. 2A-C**), which exhibits moderate kidney retention (6.1 %IA/g). Of the cleavable peptides, the GY (cleavable) and GY* (control) derivatives both exhibited elevated kidney signal (52 and 18 %IA/g, respectively). By contrast, the positioning of the MVK peptide linker was found to have a dramatic effect on renal uptake. The control conventional backbone linker DOTA-MVK-TATE showed the highest renal signal (76 %IA/g), but the DOTA-MVK(ε)-TATE exhibited among the lowest signal (4 %IA/g) of the tested variants, and, accordingly, was selected for further evaluation. The anionic linked compound DOTA-GDDDG-TATE also exhibited elevated kidney signal (13 %IA/g). Notably, the DOTA-G-Chol-G-TATE compound exhibited not only elevated kidney signal (9.6 %IA/g) but also significant retention in the blood (1.7 %IA/g), unlike the other agents which had almost completely cleared. This was an expected finding because this variant was designed to bind serum albumin. All three glycans linkers exhibited low levels of kidney retention (< 10%), but the DOTA-β-Gal-TATE was the lowest at 4.1 %IA/g and, therefore, selected for further characterization. Beyond the kidney, very low levels (<1 %IA/g) were found in the spleen, muscle, heart and femur, some accumulation (1 – 2 %IA/g) was observed for other organs (liver, intestine, lung) (**Fig. S13; Table S1**).

**Fig. 2.**
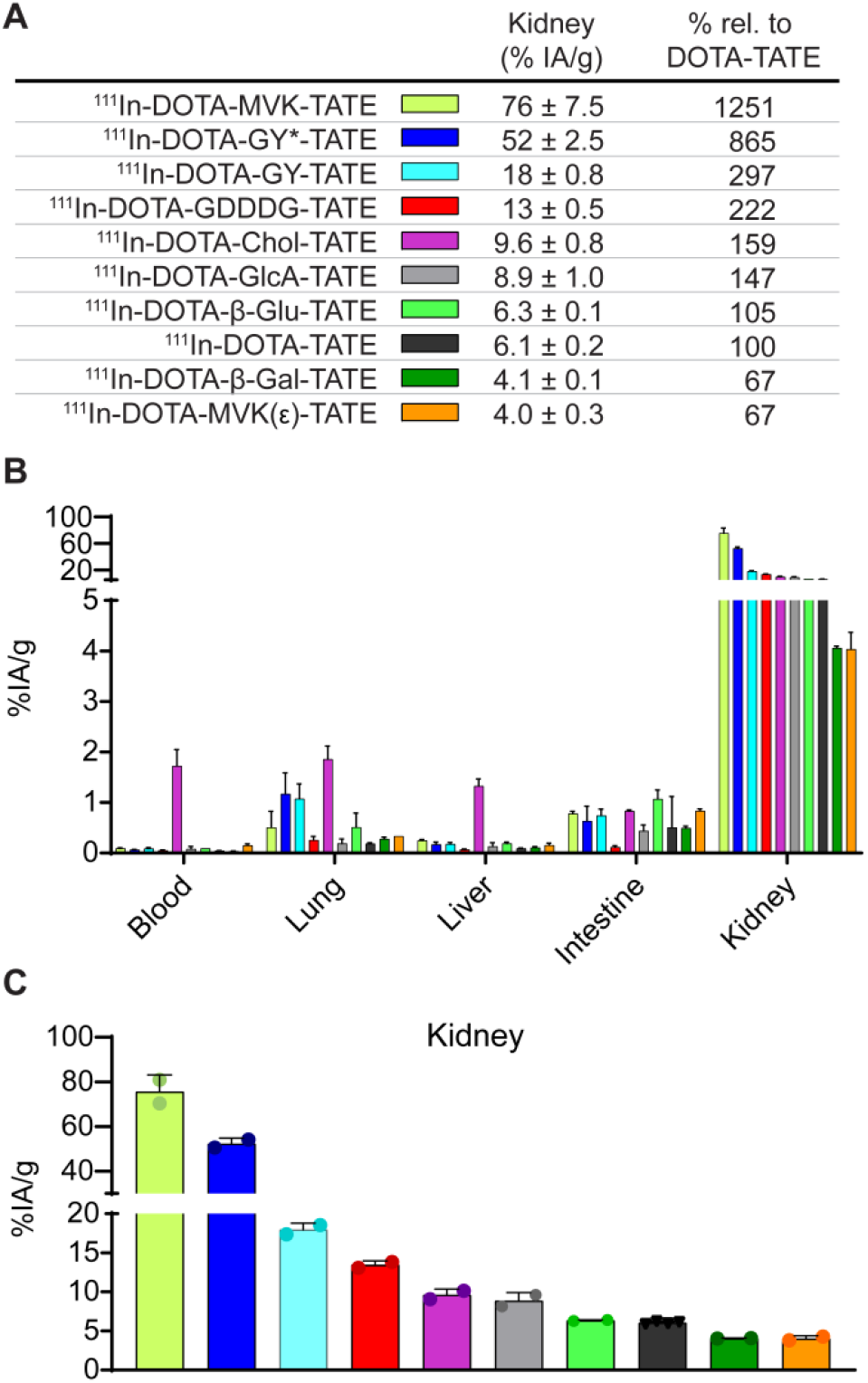
*Ex vivo* biodistribution of ^111^In-labeled DOTA-TATE derivatives in normal athymic nude mice at 2 h post injection to screen the effect in renal uptake of each linker. (**A**) Summary of kidney uptake values for each compound, expressed as % injected activity per gram of tissue (%IA/g), along with relative kidney retention compared to ^111^In-DOTA-TATE (set as 100%). (**B**) Biodistribution profiles of all ^111^In-DOTA-TATE derivatives across major organs. (**C**) Direct comparison of kidney uptake across the panel of compounds. Data are presented as mean ± standard deviation (n = 2 – 4 per cohort).

### DOTA-β-Gal-TATE demonstrates superior tumor uptake and low kidney retention in NET models by PET/CT imaging

Having identified the most promising agents from our biodistribution screen, DOTA-MVK(ε)-TATE and DOTA-β-Gal-TATE, we then compared the biodistribution of the ^111^In-labeled radioconjugates to ^111^In-DOTA-TATE in AR42J xenografts implanted in the right shoulder of athymic nude mice 1-hour post-injection (**Fig. 3A; Table S2**). ^111^In-DOTA-TATE demonstrated moderate tumor uptake (5.67 ± 0.92 %IA/g) and high renal accumulation (7.33 ± 0.04 %IA/g), yielding a tumor-to-kidney (T/K) ratio of 0.77. ^111^In-DOTA-MVK(ε)-TATE exhibited a comparable level of tumor uptake (5.44 ± 0.19 %IA/g; *p=* 0.693) but showed a notable reduction in kidney retention compared to ^111^In-DOTA-TATE (5.56 ± 0.29 %IA/g; *p* < 0.001), resulting in an improved T/K ratio of 0.98. In contrast, ^111^In-DOTA-β-Gal-TATE showed a significantly enhanced tumor uptake (10.77 ± 1.11 %IA/g, ∼90% increase compared to DOTA-TATE; *p* < 0.01), while simultaneously achieving the lowest kidney retention among the three compounds (5.19 ± 0.18 %IA/g, ∼29% reduction; *p* < 0.0001). This favorable biodistribution translated into a T/K ratio of 2.08, representing a 2.7-fold improvement over DOTA-TATE. These results highlight the potential of glycosyl cleavable linker strategies, particularly the β-galactose linker, for enhancing tumor selectivity and reducing off-target renal accumulation.

**Fig. 3.**
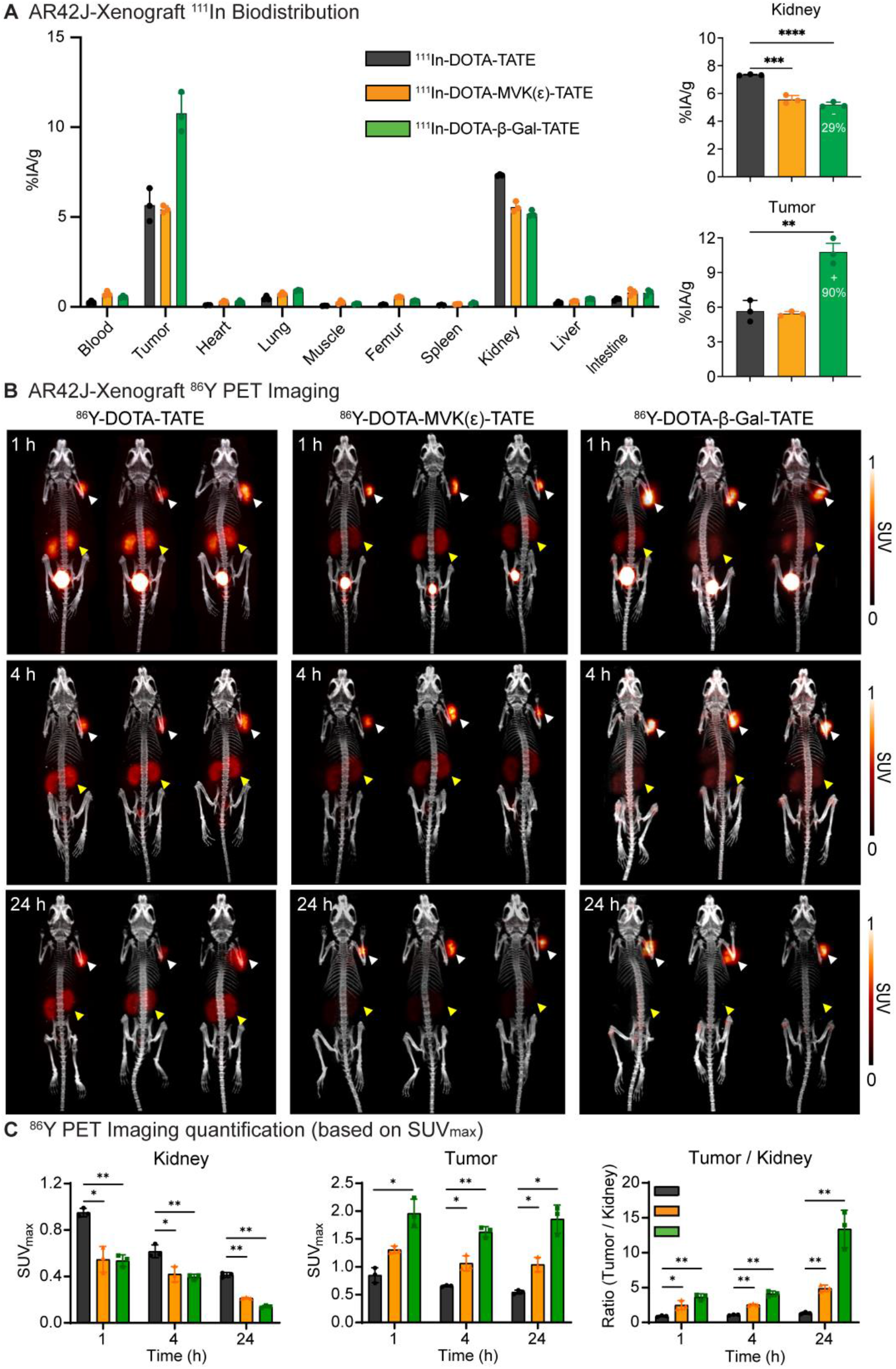
DOTA-TATE variants with cleavable linkers exhibit reduced renal uptake and improved tumor-to-kidney ratios compared to the parental DOTA-TATE in AR42J xenograft models. (**A**) *Ex vivo* biodistribution of ^111^In-labeled DOTA-TATE derivatives at 1 h post injection in AR42J tumor-bearing athymic nude mice (n = 3/cohort). Uptake in major organs including tumor and kidney is shown as %IA/g. The right panels highlight quantitative differences in kidney and tumor uptake across the three groups. (**B**) Representative maximum intensity projection (MIP) PET/CT images of ^86^Y-labeled compounds at 1, 4, and 24 h post injection. Tumors (white arrowheads) and kidneys (yellow arrowheads) are highlighted. (**C**) PET quantification based on SUV_max_ values for kidney, tumor, and tumor-to-kidney ratio at each timepoint. Data are presented as mean ± standard deviation (n = 3/cohort). Statistical significance was assessed using unpaired two-tailed t-tests: \**p* < 0.05, \*\**p* < 0.01, \*\*\**p* < 0.001, \*\*\*\**p* < 0.0001.

To assess *in vivo* biodistribution in a longitudinal fashion, PET/CT imaging and quantitative analysis was performed using ^86^Y-labeled versions of the compounds (^86^Y-DOTA-TATE, ^86^Y-DOTA-MVK(ε)-TATE, and ^86^Y-DOTA-β-Gal-TATE; **Fig. S14**) at 1, 4, 24 hours post-injection (**Fig. 3B, C**). The ^86^Y-DOTA-TATE showed prominent kidney retention (maximum standardized uptake values; SUV_max_: 0.95 ± 0.04 at 1 h, 0.62 ± 0.06 at 4 h, and 0.41 ± 0.02 at 24 h) and modest tumor signal (0.85 ± 0.14, 0.66 ± 0.02, and 0.55 ± 0.04, respectively). In contrast, ^86^Y-DOTA-MVK(ε)-TATE demonstrated a significantly reduced kidney signal (0.55 ± 0.11, 0.42 ± 0.07, and 0.21 ± 0.01, respectively; *p* < 0.05 vs ^86^Y-DOTA-TATE) and a slightly increased tumor uptake (1.31 ± 0.06, 1.06 ± 0.14, and 1.04 ± 0.13, respectively; *p* < 0.05 vs ^86^Y-DOTA-TATE). ^86^Y-DOTA-β-Gal-TATE exhibited a comparably low kidney signal as the MVK variant (0.54 ± 0.05, 0.40 ± 0.02, and 0.14 ± 0.01, respectively; *p* < 0.01 vs ^86^Y-DOTA-TATE), but achieved substantially higher tumor retention than both other agents (1.96 ± 0.26, 1.63 ± 0.09, and 1.86 ± 0.25, respectively; *p* < 0.05 vs ^86^Y-DOTA-TATE). These results explain why the tumor-to-background contrast was globally superior for ^86^Y-DOTA-β-Gal-TATE compared with the other variants. Its superior imaging was driven primarily by its significantly higher tumor uptake, resulting in a T/K ratio at 24 h was over 10-fold higher than that of ^86^Y-DOTA-TATE (*p* < 0.01) and approximately 2.7-fold higher than that of ^86^Y-DOTA-MVK(ε)-TATE (*p* < 0.05), with a similar trend observed in the mean standardized uptake value (SUV_mean_) analysis (**Fig. S15**).

To independently confirm these observations, *ex vivo* biodistribution studies were conducted using the same ^86^Y-labeled compounds at matched timepoints (**Fig. S16; Table S3**). The results closely paralleled the PET/CT imaging data, demonstrating that cleavable linker versions, particularly the β-Gal linker, exhibited significantly higher tumor uptake and reduced renal accumulation compared to the unmodified control DOTA-TATE.

Taken together, these findings unequivocally demonstrate that the β-galactose-cleavable linker dramatically enhances tumor selectivity while significantly reducing off-target renal uptake, thereby improving the T/K ratio for diagnostic imaging, and strongly supporting its potential for future therapeutic applications.

### Enzyme Cleavable Linkers Enable Renal Excretion of DOTA-TATE Derivatives

To elucidate the role of linker cleavage in reducing renal retention, we performed metabolite analysis of ^86^Y-labeled DOTA-TATE derivatives in normal, non-tumor-bearing athymic nu/nu mice. Mice were intravenously administered ^86^Y-DOTA-MVK(ε)-TATE or ^86^Y-DOTA-β-Gal-TATE (13–15 MBq in 200 μL PBS, 4 μg), and urine samples were collected at 1 hour post-injection. These samples were analyzed by radio-HPLC alongside synthetic standards corresponding to the expected cleavage products: ^86^Y-DOTA-G-M-OH and ^86^Y-DOTA-NHMe (**Fig. 4A**).

**Fig. 4.**
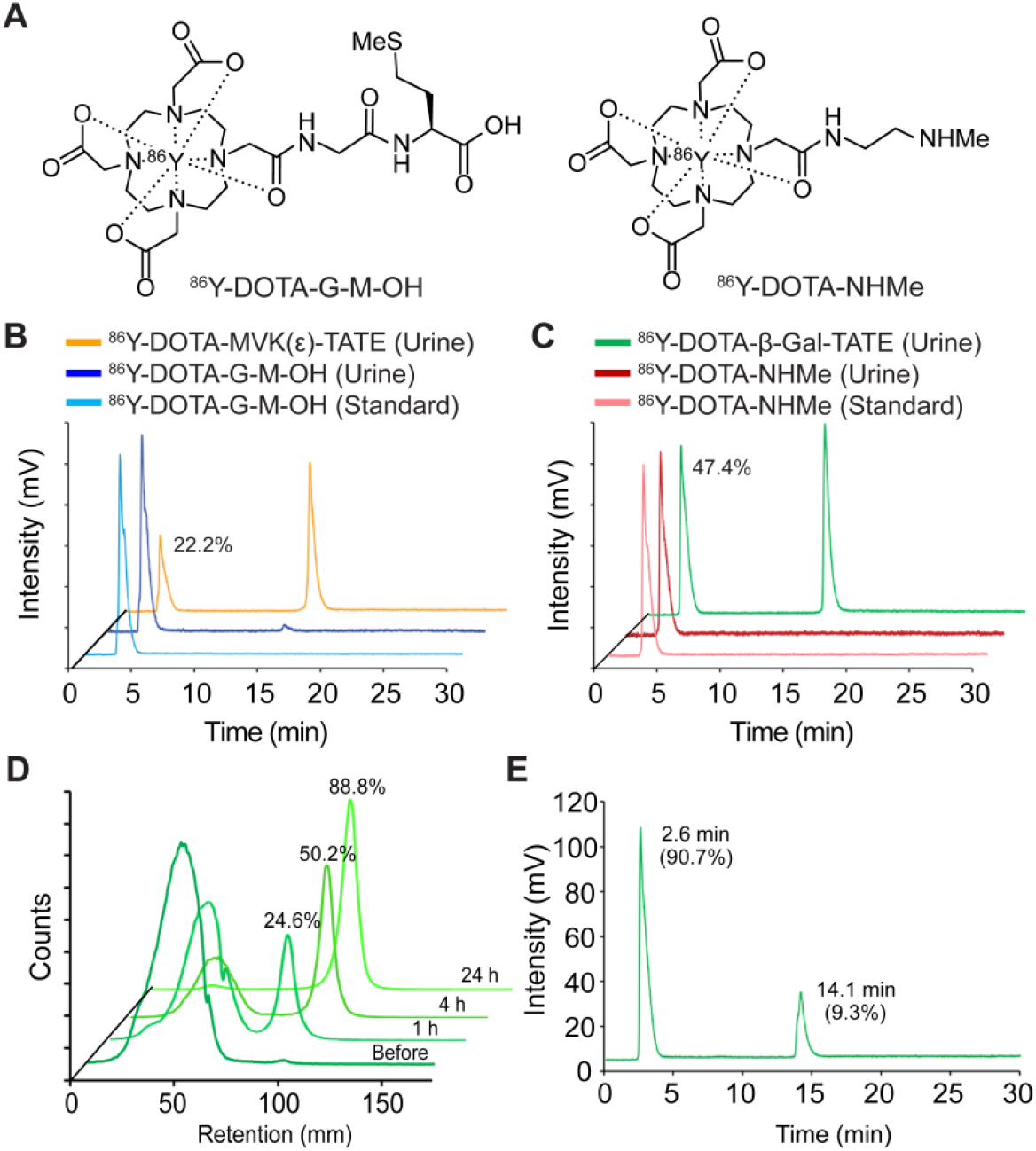
Enzymatic cleavage and renal excretion of DOTA-linker-TATE derivatives confirmed by radio-HPLC and radio-TLC analyses. (**A**) Chemical structures of the synthetic metabolite standards: ^86^Y-DOTA-G-M-OH, derived from ^86^Y-DOTA-MVK(ε)-TATE, and ^86^Y-DOTA-NHMe, derived from ^86^Y-DOTA-β-Gal-TATE. (**B**) Radio-HPLC profiles of urine samples collected 1 h post-injection of ^86^Y-DOTA-MVK(ε)-TATE and ^86^Y-DOTA-G-M-OH in normal athymic mice, along with the standard ^86^Y-DOTA-G-M-OH. (**C**) Radio-HPLC profiles of urine samples collected 1 h post-injection of ^86^Y-DOTA-β-Gal-TATE and ^86^Y-DOTA-NHMe in normal athymic mice, along with the standard ^86^Y-DOTA-NHMe. (**D**) Radio-TLC analysis of ^86^Y-DOTA-β-Gal-TATE following incubation with β-galactosidase at multiple time points. (**E**) Radio-HPLC chromatogram of the 24-hour β-galactosidase-treated sample of ^86^Y-DOTA-β-Gal-TATE, analyzed to confirm metabolite identity.

Radio-HPLC chromatograms of urine confirmed that both agents underwent enzymatic cleavage *in vivo*, generating metabolites that co-eluted precisely with their corresponding standards. Specifically, ^86^Y-DOTA-MVK(ε)-TATE was metabolized to ^86^Y-DOTA-G-M-OH, accounting for 22.2% of total urinary radioactivity (**Fig. 4B**), while ^86^Y-DOTA-β-Gal-TATE showed more extensive cleavage, with 47.4% of radioactivity corresponding to ^86^Y-DOTA-NHMe (**Fig. 4C**). These findings demonstrate *in vivo* enzymatic cleavage of both MVK(ε) and β-Gal linkers, with rapid urinary excretion of the resulting metabolites.

To further confirm β-galactosidase-specific cleavage, ^86^Y-DOTA-β-Gal-TATE was incubated *in vitro* with β-galactosidase and analyzed by radio-TLC over time. A time-dependent increase in the cleaved product ^86^Y-DOTA-NHMe was observed (**Fig. 4D**), accounting for 24.6%, 50.2%, and 88.8% of the total activity at 1, 4, and 24 hours, respectively. Follow-up radio-HPLC analysis of the 24 h sample (**Fig. 4E**) confirmed that 90.7% of the compound had been converted to ^86^Y-DOTA-NHMe, corroborating the radio-TLC findings.

Together, these data provide strong biochemical evidence that the rapid enzymatic cleavage of the β-galactose linker generates readily cleared, low-molecular-weight metabolites, thereby improving the observed *in vivo* T/K ratios and highlighting its translational potential for optimizing radiopharmaceutical design.

### Alpha-particle therapy with ^225^Ac-DOTA-TATE variants improves tumor control and survival in NET models, with ^225^Ac-DOTA-β-Gal-TATE showing the most favorable toxicity profile

We next evaluated whether the improved pharmacokinetic profiles conferred by cleavable linker constructs would translate into enhanced therapeutic efficacy and safety. Prior to conducting targeted alpha-particle therapy studies using three ^225^Ac-labeled variants (**Fig. S17**) in AR42J xenograft-bearing mice, we first assessed their *in vivo* pharmacokinetics through *ex vivo* biodistribution studies at 24 h post-injection (**Fig. 5A; Tables S4**).

**Fig. 5.**
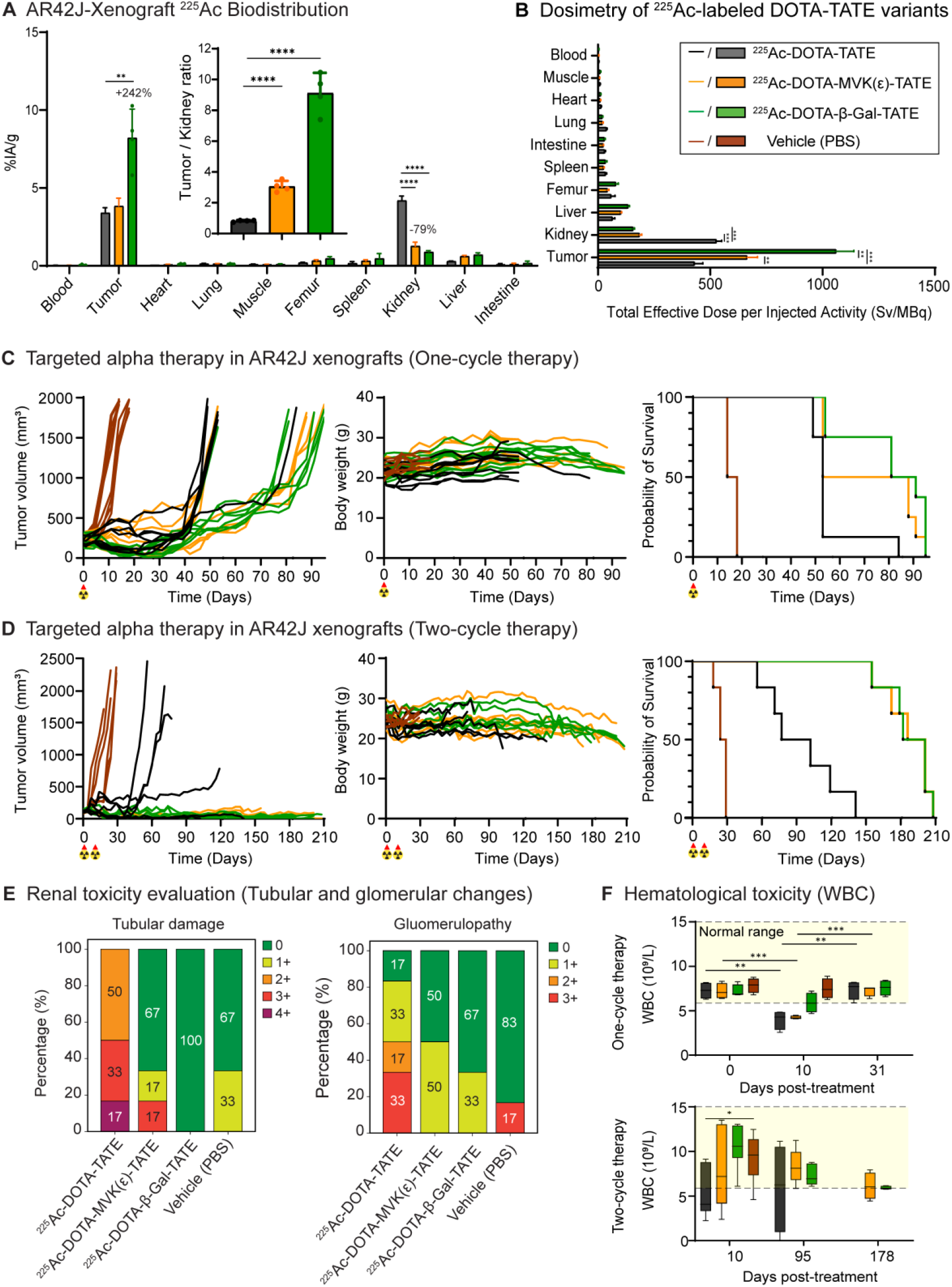
Targeted alpha therapy with ^225^Ac-DOTA-MVK(ε)-TATE and ^225^Ac-DOTA-β-Gal-TATE reduces renal uptake and enhances therapeutic efficacy and safety in murine models of neuroendocrine tumors. (**A**) *Ex vivo* biodistribution of ^225^Ac-labeled DOTA-TATE variants in AR42J xenograft-bearing mice at 24 hours post-injection (n = 4/cohort). Tumor-to-kidney uptake ratios are shown as insets. (**B**) Dosimetry estimates of ^225^Ac-DOTA-TATE, ^225^Ac-DOTA-MVK(ε)-TATE, and ^225^Ac-DOTA-β-Gal-TATE across major organs, calculated as total effective dose per injected activity (mSv/MBq). (**C**) Tumor volume, body weight, and Kaplan–Meier survival curves of mice treated with a one-cycle regimen(148 kBq) of ^225^Ac-labeled variants or vehicle control (n = 8/cohort). (**D**) Therapeutic responses in mice receiving a two-cycle regimen (2 × 148 kBq, 8-day interval) of each agent or vehicle control, showing tumor volume, body weight, and Kaplan–Meier survival outcomes (n = 6/cohort). (**E**) Quantitative renal toxicity scoring based on histological evaluation of tubular damage and glomerulopathy severity (0–4+ scale). (**F**) Peripheral white blood cell (WBC) counts for AR42J xenograft–bearing mice treated with vehicle or ^225^Ac-labeled DOTA-TATE variants under a one-cycle regimen (top) or a two-cycle regimen (bottom). Shaded areas indicate reference ranges. Results are presented as mean ± standard deviation. Statistical significance was determined using unpaired t-tests. \**p* < 0.05, \*\**p* < 0.01, \*\*\**p* < 0.001, \*\*\*\**p* < 0.0001.

To ensure the stability of the constructs prior to *in vivo* studies, we first assessed the radiochemical stability of the ^225^Ac-labeled variants. All three variants were incubated in PBS and human serum at 37°C and analyzed over a 10-day period using radio-TLC. As shown in **Fig. S18**, all constructs, including those incorporating cleavable linkers, maintained high stability with radiochemical purity above 95 percent for up to 7 days. After day 7, a gradual decline in stability was observed, with radiochemical purity dropping below 90 percent by day 10. These results indicate that while cleavable linkers do not compromise the short-term stability of the ^225^Ac-DOTA complex, degradation may occur over extended periods. This high stability within the initial 7-day therapeutic window confirms that the constructs are suitable for the subsequent *in vivo* evaluations of their pharmacokinetic and therapeutic profiles.

Compared to ^225^Ac-DOTA-TATE (4.19 ± 0.28 %IA/g), renal uptake was significantly reduced for ^225^Ac-DOTA-MVK(ε)-TATE (1.27 ± 0.25 %IA/g; *p* < 0.0001) and ^225^Ac-DOTA-β-Gal-TATE (0.89 ± 0.08 %IA/g; *p* < 0.0001). Notably, ^225^Ac-DOTA-β-Gal-TATE also showed markedly higher tumor uptake (8.22 ± 1.84 %IA/g) relative to ^225^Ac-DOTA-TATE (3.40 ± 0.34 %IA/g) and ^225^Ac-DOTA-MVK(ε)-TATE (3.83 ± 0.51 %IA/g), resulting in significantly improved tumor-to-kidney (T/K) ratios: 9.12 ± 1.31 for ^225^Ac-DOTA-β-Gal-TATE and 3.05 ± 0.37 for ^225^Ac-DOTA-MVK(ε)-TATE, compared to 0.81 ± 0.06 for ^225^Ac-DOTA-TATE (*p* < 0.0001). Although the liver uptake of ^225^Ac-DOTA-β-Gal-TATE (0.89 ± 0.49 %IA/g) was higher than that of ^225^Ac-DOTA-TATE (0.29 ± 0.02 %IA/g) and ^225^Ac-DOTA-MVK(ε)-TATE (0.61 ± 0.06 %IA/g), greater inter-mouse variability was observed. Collectively, the enhanced tumor targeting and substantially reduced renal retention observed with the β-Gal cleavable linker-containing variant support its potential to improve the therapeutic index as targeted alpha therapy compared to the others tested.

To evaluate the organ-level radiation absorbed dose resulting from different DOTA-TATE derivatives, we calculated the total effective dose per unit of injected activity (Eff Dose/IA, mSv/MBq) for ^225^Ac-labeled variants based on biodistribution data in AR42J xenograft-bearing mice (**Tables S5**). Dosimetric calculations showed that cleavable linkers markedly reduced renal radiation dose following administration of ^225^Ac-labeled variants (**Fig. 5B**). The estimated kidney dose for ^225^Ac-DOTA-TATE was 5.27 × 10^5^ mSv/MBq. In contrast, ^225^Ac-DOTA-MVK(ε)-TATE and ^225^Ac-DOTA-β-Gal-TATE showed substantially lower doses of 1.85 × 10^5^ and 1.56 × 10^5^ mSv/MBq (*p* < 0.001), respectively, corresponding to reductions of approximately 65 to 70 percent.

Tumor dose was significantly enhanced with the β-Gal variant. The absorbed dose to tumor tissue for ^225^Ac-DOTA-β-Gal-TATE reached 1.06 × 10^6^ mSv/MBq, which is approximately 2.5 times higher than that of ^225^Ac-DOTA-TATE (4.30 × 10^5^ mSv/MBq; *p* < 0.001) and 1.6 times higher than that of ^225^Ac-DOTA-MVK(ε)-TATE (6.64 × 10^5^ mSv/MBq; *p* < 0.01). In other normal tissues such as the liver, spleen, and lungs, ^225^Ac-DOTA-β-Gal-TATE exhibited slightly elevated absorbed doses compared to the parental compound. However, the liver also showed relatively higher variability in the β-Gal group, with a standard deviation of 5.92 × 10^3^ mSv/MBq, suggesting inter-animal differences in hepatic clearance or metabolism. Despite these modest increases in non-target tissues, the overall therapeutic index, as reflected by tumor-to-kidney and tumor-to-normal tissue dose ratios, remained most favorable for ^225^Ac-DOTA-β-Gal-TATE. These results support its potential to improve the balance between efficacy and safety in targeted alpha therapy.

Building on the favorable dosimetry profiles observed for the cleavable linker-modified constructs, we next evaluated their therapeutic efficacy *in vivo* following one cycle of ^225^Ac-labeled variants (**Fig. 5C; Fig. S19A**). AR42J xenograft-bearing mice were randomized into four groups and treated with a one-cycle regimen of either PBS (vehicle), ^225^Ac-DOTA-TATE, ^225^Ac-DOTA-MVK(ε)-TATE, or ^225^Ac-DOTA-β-Gal-TATE at an administered activity of 148 kBq per mouse. Tumor growth, overall survival, and body weight were monitored over 95 days. All treatment groups exhibited suppression of tumor growth compared to the vehicle group, which showed rapid and continuous tumor progression. Among the other cohorts, ^225^Ac-DOTA-β-Gal-TATE demonstrated the most sustained inhibition of tumor growth, with minimal progression observed up to day 35 post-treatment. Notably, the ^225^Ac-DOTA-β-Gal-TATE group maintained lower tumor volumes than the ^225^Ac-DOTA-MVK(ε)-TATE group from day 11 post-treatment onward, with statistically significant differences at multiple subsequent timepoints, supporting greater local tumor growth control.

In contrast, mice treated with ^225^Ac-DOTA-TATE showed only a temporary delay in tumor growth, with rapid regrowth observed after day 39. The ^225^Ac-DOTA-MVK(ε)-TATE group showed a moderate, but sustained, therapeutic effect. Although tumor uptake at 24 hours post-injection was similar between the ^225^Ac-DOTA-TATE and ^225^Ac-DOTA-MVK(ε)-TATE-treated groups, this did not account for the difference in therapeutic outcomes. To investigate further, a biodistribution study at 4 days showed that ^225^Ac-DOTA-MVK(ε)-TATE had significantly higher tumor uptake (2.41 ± 0.22 %IA/g) than ^225^Ac-DOTA-TATE (1.14 ± 0.37 %IA/g; *p* < 0.001), along with a higher tumor-to-kidney ratio (5.72 ± 0.91 versus 1.15 ± 0.44) (**Fig. S20**). These results suggest that the MVK(ε) linker improves tumor retention over time, contributing to its enhanced therapeutic efficacy despite similar early pharmacokinetics.

Consistent with these growth-control trends, absorbed-dose estimates at 148 kBq per mouse showed that ^225^Ac-DOTA-β-Gal-TATE delivered the greatest tumor dose (32.5 Gy) with a comparatively low kidney dose (4.8 Gy) among the three agents. By comparison, ^225^Ac-DOTA-MVK(ε)-TATE and ^225^Ac-DOTA-TATE delivered 20.4 and 13.2 Gy to tumor with kidney doses of 5.7 and 16.2 Gy. Thus, ^225^Ac-DOTA-β-Gal-TATE provided 2.46- and 1.59-fold higher tumor dose than ^225^Ac-DOTA-TATE and ^225^Ac-DOTA-MVK(ε)-TATE, respectively, while reducing kidney dose by about 70 percent versus ^225^Ac-DOTA-TATE and 15 percent versus ^225^Ac-DOTA-MVK(ε)-TATE. The differential dosing provides the biological basis for longer survival, as reflected in the overall survival results. Median survival times were 86 days (95% CI: 66.1–95.4) for ^225^Ac-DOTA-β-Gal-TATE, 70.5 days (95% CI: 54.9–88.6) for ^225^Ac-DOTA-MVK(ε)-TATE, and 53 days (95% CI: 46.3–65.5) for ^225^Ac-DOTA-TATE, while the vehicle group had a median survival of only 16 days (95% CI: 14.2–17.8). Each group included eight animals, and no subjects were censored. Kaplan–Meier survival analysis showed significant differences among the four treatment groups (log-rank test, χ^2^ = 44.07, df = 3, *p* < 0.0001; Gehan–Breslow–Wilcoxon test, χ^2^ = 40.65, df = 3, *p* < 0.0001), with the cleavable linker groups, particularly ^225^Ac-DOTA-β-Gal-TATE, demonstrating the greatest survival benefit. These findings support the potential of cleavable linker strategies to enhance therapeutic efficacy by improving tumor targeting and reducing off-target toxicity.

Body weight monitoring indicated that all treated animals experienced mild weight loss during the first week post-injection, consistent with the expected acute effects of alpha therapy. Mice treated with ^225^Ac-DOTA-β-Gal-TATE or ^225^Ac-DOTA-MVK(ε)-TATE gradually regained weight by day 14, maintaining stable body weights for the remainder of the study (**Fig. S19B**). In contrast, the ^225^Ac-DOTA-TATE group consistently showed lower body weights compared to the cleavable linker-treated groups, suggesting less effective tumor control and greater systemic impact. Mice in the vehicle group rapidly lost weight and were euthanized by day 18 due to tumor burden, preventing further evaluation.

To further assess therapeutic efficacy, AR42J xenograft-bearing mice received two cycles of ^225^Ac-labeled constructs (2 × 148 kBq, administered 8 days apart) (**Fig. 5D**). All treated groups exhibited enhanced antitumor activity compared with the vehicle control. Mice treated with ^225^Ac-DOTA-β-Gal-TATE or ^225^Ac-DOTA-MVK(ε)-TATE achieved prolonged tumor control with sustained growth inhibition, whereas the ^225^Ac-DOTA-TATE group showed a more transient response (**Fig. S19C**). Complete tumor regression occurred in two of six animals in each cleavable linker group. By the final monitoring point, tumor volumes remained below 100 mm^3^ in five of six mice in the ^225^Ac-DOTA-MVK(ε)-TATE group and in all six mice in the ^225^Ac-DOTA-β-Gal-TATE group, with both cleavable linker groups maintaining significantly smaller tumors throughout the study. These tumor control benefits were reflected in survival outcomes, with median survival times of 193.5 days (95% CI: 165.7–208.6) for ^225^Ac-DOTA-MVK(ε)-TATE, 191.5 days (95% CI: 167.0–208.4) for ^225^Ac-DOTA-β-Gal-TATE, and 89.5 days (95% CI: 60.6–128.0) for ^225^Ac-DOTA-TATE, compared with only 26.5 days (95% CI: 20.9– 30.1) for the vehicle group. Kaplan–Meier survival analysis confirmed significant differences among the four treatment groups (log-rank test, χ^2^ = 39.48, df = 3, *p* < 0.0001; Gehan–Breslow– Wilcoxon test, χ^2^ = 34.31, df = 3, *p* < 0.0001), with both cleavable linker groups showing markedly prolonged survival relative to the parental ^225^Ac-DOTA-TATE. However, pairwise comparisons between the two cleavable-linker groups did not detect a statistically significant difference in overall survival.

Body weight monitoring showed good tolerability for all radioligand-treated groups. Transient weight loss was observed following treatment administration, but mice gradually recovered and maintained stable body weights (**Fig. S19D**). The cleavable linker groups, in particular, demonstrated long-term weight stability, consistent with their superior tumor control and minimal systemic toxicity. These results support the therapeutic advantage of cleavable linker designs in targeted alpha-particle therapy under a two-cycle regimen.

To further assess tissue-level toxicity at the conclusion of the two-cycle therapy study, kidneys, livers, spleens, and tumors from representative mice were collected and evaluated histologically using hematoxylin and eosin (H&E) staining. Microscopic pathology was assessed and recorded in the MHL Pathology Database (Zaphod) according to a standardized five-point grading scale (normal, minimal, mild, moderate, marked). Terminology and criteria for pathological findings were consistent with the INHAND (International Harmonization of Nomenclature and Diagnostic Criteria) guidelines developed by the Society of Toxicologic Pathology.

The most prominent treatment-related effects were observed in the kidneys. Mice treated with ^225^Ac-DOTA-TATE showed multifocal tubular injury and glomerular changes, including tubular degeneration, epithelial necrosis, and karyomegaly in the cortical tubules, along with increased mesangial matrix in the glomeruli (**Fig. S21, S22**). Ordinal scoring corroborated these observations (**Fig. 5E**). For tubular damage, ^225^Ac-DOTA-TATE exhibited the most injury (50% grade 2+, 33% grade 3+, 17% grade 4+), whereas ^225^Ac-DOTA-MVK(ε)-TATE showed 17% grade 3+, 17% grade 1+, and 67% grade 0, and ^225^Ac-DOTA-β-Gal-TATE was 100% grade 0. For glomerulopathy, ^225^Ac-DOTA-TATE again scored worst, while MVK(ε) and β-Gal were largely grade 0 (50% and 67%, respectively; vehicle 83% grade 0). Thus, although both cleavable-linker constructs reduced renal injury relative to ^225^Ac-DOTA-TATE, the MVK(ε) linker-containing variant exhibited more tubular damage than β-Gal linker variant, underscoring the comparatively favorable renal safety of the β-Gal linker. These histological findings align with the lower renal absorbed doses for the cleavable-linker groups in the dosimetry analysis.

Liver sections were also examined (**Fig. S23**). Observed changes were minimal to mild and generally reactive in nature, including extramedullary hematopoiesis (EMH), which was interpreted as a response to tumor burden rather than treatment-related toxicity. Similarly, minimal treatment-related changes were observed in spleens across all treatment groups. Tumor sections, when present, were notably smaller in the cleavable linker groups, and were often associated with increased macrophage infiltration in the surrounding stroma (**Fig. S24**). These findings are consistent with the superior tumor control observed in the two-cycle therapy study.

Blood samples were collected at multiple timepoints during therapy studies and analyzed to assess toxicity (**Fig. S25**). Biochemical parameters of liver (e.g. alanine aminotransferase, bilirubin) and renal (blood urea nitrogen, and creatinine) toxicity remained mostly within normal limits across all groups. Hematologic markers such as hemoglobin levels and platelet counts were stable, while white blood cell counts showed a transient decrease following one cycle of ^225^Ac treatment that recovered at later time points in the β-Gal and MVK(ε) groups (**Fig. 5F)**. These findings further support the favorable tolerability profiles of the cleavable linker constructs. Importantly, the fact that blood markers for nephrotoxicity (e.g. blood urea nitrogen, creatinine) were unremarkable throughout the course of the therapy studies, suggests that these canonical markers do not reflect the histopathological changes occurring contemporaneously.

Taken together, these results support the use of enzyme-cleavable linkers as a promising strategy to enhance the therapeutic index of targeted alpha therapies. In particular, the β-galactose-cleavable variant exhibited superior efficacy and a favorable safety profile, warranting further translational investigation.

## DISCUSSION

We report on the systematic evaluation of a small library of DOTA-TATE derivatives containing unique linkers with distinct properties and identify a novel construct, DOTA-β-Gal-TATE, a glycosidase-cleavable linker that exhibits superior tumor uptake and lower kidney retention by *ex vivo* biodistribution and PET imaging studies when compared with the parental DOTA-TATE and the renal brush border enzyme-cleavable derivative DOTA-MVK(ε)-TATE. All three of variants exhibited therapeutic benefit when radiolabeled with the alpha particle-emitting radionuclide ^225^Ac. However, renal histopathology was assessed in the two-cycle treatment cohorts and demonstrated the greatest tubular and glomerular injury with ^225^Ac-DOTA-TATE, and lower injury with the cleavable-linker constructs. More tubulopathy was observed with ^225^Ac-DOTA-MVK(ε)-TATE than with ^225^Ac-DOTA-β-Gal-TATE. Consistent with these findings, ^225^Ac-DOTA-β-Gal-TATE not only achieved the most favorable renal safety profile but also demonstrated local control and overall survival comparable to ^225^Ac-DOTA-MVK(ε)-TATE in murine models of NETs, while both cleavable-linker groups outperformed ^225^Ac-DOTA-TATE.

Although dosimetry estimates predicted a higher tumor-absorbed dose for ^225^Ac-DOTA-β-Gal-TATE, this did not translate into a statistically significant local control or overall survival benefit compared to ^225^Ac-DOTA-MVK(ε)-TATE. This discrepancy suggests that the absorbed dose–effect relationship for tumors of these alpha-emitters is not yet fully characterized in this model. A plausible explanation is that the administered activities resulted in absorbed doses that fall on the initial, shallow portion of a sigmoidal dose–response curve, where incremental dose increases yield only marginal gains in therapeutic outcome. In contrast, a clear dose-effect relationship was evident in the kidneys, where the lower renal absorbed doses in the cleavable-linker groups directly corresponded with reduced histopathological toxicity. Importantly, we were only able to assess whole organ dosimetry and not microdosimetry in subcompartments of kidneys, which could influence the dose-effect as it relates to end organ toxicity.

For this work, we focused on DOTA-TATE because it is an approved clinical agent for imaging (^68^Ga, ^64^Cu) and treatment (^177^Lu) for patients with SSTR-positive NETs. Of note, ^225^Ac-DOTA-TATE is being investigated in patients with NETs who have progressed or recurred after ^177^Lu-DOTA-TATE treatment in a late-phase clinical trial (NCT05477576). The significant renal retention of these peptide-based agents is a point of concern, and is typically mitigated by positively-charged amino acid infusions over several hours to competitively inhibit proximal tubule reabsorption*(16–18)*. Although effective in reducing renal absorbed dose by 9–53%*(19, 20)*, these infusions are logistically burdensome. Additional strategies such as gelofusine infusion*(21)*, use of radioprotectants (e.g., amifostine, α1-microglobulin)*(22, 23)*, or pharmacologic inhibitors of endocytosis*(24, 25)* have been proposed or evaluated in preclinical studies. However, their clinical adoption has been limited due to risks of allergic reactions, toxicity, or lack of efficacy. Personalized dosimetry and dose fractionation have also been explored to optimize therapeutic indices*(26)*, yet these require specialized resources and are not yet universally implemented. There are potentially significant benefits to the creation of optimized agents with reduced intrinsic toxicity due to kidney retention, and potentially improved tumor uptake.

One strategy to improve targeting is the installation of albumin-binding domains (e.g., Evans blue, ibuprofen and related derivatives)*(27–30)*. We chose to test the sterol-linked derivative DOTA-G-Chol-G-TATE. In normal mice, biodistribution screening performed at 2 h post-injection exhibited markedly prolonged blood circulation time and elevated kidney retention (9.6 %IA/g) compared with DOTA-TATE (6 %IA/g). While albumin-binding strategies, exemplified by the promising clinical trial results of agents like DOTA-EB-TATE in humans*(31)*, have been shown to reduce renal accumulation and improve tumor targeting*(32)*, prolonged circulation has also been associated with increased kidney uptake and potential bone marrow toxicity*(33)*. Given the elevated and widespread organ uptake observed with DOTA-G-Chol-G-TATE in our study, biodistribution studies at later time points in tumor-bearing models will be essential to fully characterize its pharmacokinetics and guide further structural optimization. However, these studies suggest that when rapid tumor targeting is desirable, for example with early time point imaging or in therapeutic contexts, our cholesterol-based albumin binding strategy is not optimal.

We also examined the influence of molecular charge on renal retention. Several studies have suggested that anionic charges can reduce renal uptake of related TATE derivatives*(12, 34)*, prompting us to investigate a tri-anionic DOTA-GDDDG-TATE derivative. Unexpectedly, this construct exhibited higher kidney retention (13 %IA/g) compared with the parental DOTA-TATE (6 %IA/g). Although negative charges have commonly been employed to mitigate renal uptake, recent reports have paradoxically shown lower renal retention for certain positively charged ligands*(35)*. Taken together, these findings suggest that the effect of net molecular charge on renal reabsorption is context-dependent and may be secondary to other physicochemical or biological factors.

The incorporation of cleavable linkers is a promising strategy to reduce off-target effects like renal retention without requiring additional supportive care*(36)*. Several prior studies have investigated cleavable linkers in radiopharmaceuticals guided by the general hypothesis of reducing renal reuptake through the action of abundant proteases in the renal brush border*(37– 40)*. These prior efforts included two peptidic linkers we chose to investigate including GY, which is a putative substrate for carboxypeptidase M (CPM), and MVK(ε)-, which is a substrate for the renal brush border enzyme neutral endopeptidase (NEP)*(8–11)*. Critically, neither the GY or GY* linkers improved tumor targeting compared to DOTA-TATE although, the natural isomer L-tyrosine GY compound exhibited moderately lower uptake (18 %IA/g) than the unnatural GY*, suggesting that some cleavage may occur. By contrast, the MVK(ε) linker showed dramatic impact of the positioning of the MVK moiety on renal retention. The conventional backbone linker DOTA-MVK-TATE exhibited the highest kidney signal (76 %IA/g), whereas DOTA-MVK(ε)-TATE, in which the MVK was conjugated via the ε-amino group of lysine, showed one of the lowest signals (4 %IA/g). This pronounced difference is likely due to steric hindrance in the conventional DOTA-MVK-TATE structure, which limits NEP accessibility, particularly to the M–V bond, thereby reducing enzymatic recognition and cleavage efficiency*(41, 42)*. Notably, while preclinical efforts have clearly demonstrated that MVK(ε)-based linkers can dramatically lower kidney retention of radioconjugates in murine models, whether these improvements translate clinically can only be evaluated in human studies*(43)*.

Finally, noting studies from ADC linker efforts*(44, 45)*, we identified a panel of glycans that are cleavable by naturally occurring glycosidases. These include DOTA-linker-TATE variants of β-galactose (β-Gal), β-glucose (β-Glu), and β-glucuronide (β-GlcA). The presence of these enzymes in the kidney has been directly demonstrated in classic biochemical work by Robinson et al.*(46)* which identified and characterized β-galactosidase, β-glucosidase, and β-glucuronidase in rat kidney tissue. Critically, this finding extends to humans, as data from the Human Protein Atlas confirms significant protein expression of β-galactosidase (*GLB1*), β-glucosidase (*GBA*), and β-glucuronidase (*GUSB*) in the human renal tubules*(47)*. Although the study did not quantify their relative abundance, the high metabolic and reabsorptive activity of the proximal tubule suggests that these enzymes are present at functionally significant levels, supporting the feasibility of glycan linker cleavage in this setting. To our knowledge, such cleavable-glycan linkers have not been tested previously in the context of SSTR targeting or for radiopharmaceutical purposes.

Our initial hypothesis was that glycosidase activity might reduce kidney retention and subsequent absorbed dose to the organs. Indeed, dosimetric analysis showed that cleavable linker incorporation reduced the renal absorbed dose by approximately 65–70% compared with ^225^Ac-DOTA-TATE. Histopathology confirmed this protective effect, showing the greatest tubular and glomerular injury with ^225^Ac-DOTA-TATE, appreciable but reduced tubular damage with ^225^Ac-DOTA-MVK(ε)-TATE, but minimal changes with ^225^Ac-DOTA-β-Gal-TATE, consistent with the lower renal absorbed doses observed in the cleavable-linker groups. While these results indicate that cleavable linker strategies can mitigate alpha-particle–induced nephrotoxicity without compromising therapeutic efficacy, additional safety considerations remain. Decay of ^225^Ac produces alpha particles along with recoiled daughter isotopes such as ^221^Fr and ^213^Bi, raising the possibility of redistribution and off-target radiation dose. Although prior work, including the study by Tafreshi et al.*(48)*, has shown relatively low kidney signal from ^225^Ac and its daughters, a systematic assessment of ^213^Bi redistribution and renal accumulation will be important to fully characterize long-term safety.

An unanticipated, but striking, finding of these studies was the finding that the β-Gal linker increased tumor uptake. This finding was most pronounced on PET imaging, with ^86^Y-DOTA-β-Gal-TATE demonstrating substantially higher tumor retention compared to ^86^Y-DOTA-TATE, and is likely also responsible for the excellent efficacy of the β-Gal derivative. The origin of this effect is difficult to assess experimentally. However, we hypothesize that at least two complementary features may be contributing. Various studies have shown that glycans improve certain key physical properties of proteins, peptides and small molecules, and this enhances the circulating half-life leading to improved target-dependent binding*(49–53)*. A second feature that may contribute is glycosidase-mediated bond cleavage, perhaps in the tumor microenvironment, and retention of the cleaved chelate product (^86^Y-DOTA-NHMe, **Fig. 4A**). Glycosidase activity is elevated in many solid tumors, part of the motivation for glycan linkers in ADC and other prodrug settings*(54)*. Furthermore, several imaging-oriented studies have suggested that basic amine containing imaging probes exhibit lysosomotropic properties that lead to improved tumor retention*(55, 56)*. While further efforts are needed to fully distinguish these potentially synergistic mechanisms, our findings suggest that simultaneously increasing tumor uptake and decreasing kidney retention is possible with these optimized linker domains.

In sum, we demonstrate that the use of a glycosidase-cleavable linker for radiotheranostics applications confers a biodistribution, local control, overall survival, and toxicity advantage in murine models of NETs. Future studies will explore the generalizability of this approach across other biomolecule types (e.g. full-length antibodies and antibody fragments, peptidomimetics). If successful, this technology could transform radioconjugates limited to diagnostic purposes due to unfavorable pharmacokinetics and biodistributions to ones with therapeutic potential. Such approaches are critical to the further development of peptides, miniproteins, and antibody fragments as targeting vectors for radiopharmaceutical therapy.

## MATERIALS AND METHODS

### Synthesis of DOTA-TATE variants

DOTA-TATE derivatives were synthesized using standard solid-phase peptide synthesis (SPPS) methods and characterized by LC/HRMS. Detailed synthetic procedures and characterization data are provided in the Supplementary Information (**Schemes S1–S5**; **Figs. S1–S11**).

### Radiolabeling of DOTA-TATE variants

Radiolabeling of DOTA-TATE and its derivatives with ^111^In, ^86^Y, and ^225^Ac was performed using standard conditions optimized for each isotope, achieving radiochemical yields greater than 95% and high radiochemical purity. Analytical methods included radio-TLC and radio-HPLC to confirm product identity and purity. Detailed radiolabeling protocols, including reagent compositions, incubation conditions, and analytical parameters, are provided in the Supplementary Information (**Figs. S12, S14, S17**).

### Cell lines

AR42J cell lines were purchased from ATCC (Manassas, VA). AR42J cells were cultured in Kaighn’s Modification of Ham’s F-12 (ATCC) supplemented with 20% heat-inactivated fetal bovine serum (HI-FBS, Gibco, Grand Island, NY, USA) at 37 °C under 5% CO_2_, as recommended by ATCC. Cell lines were used for experiments within 15 passages to limit genetic drift.

### PET imaging and biodistribution

All animal procedures were performed in accordance with institutional guidelines and were approved by the Institutional Animal Care and Use Committee (IACUC) of the National Institutes of Health. Athymic nu/nu mice (female, 5 weeks, 16−18 g) were obtained from Charles River Laboratories (Wilmington, MA, USA) and allowed to acclimate for 2 weeks prior to the commencement of experiments. Biodistribution and PET imaging studies were performed in normal and AR42J tumor-bearing athymic nude mice to evaluate the pharmacokinetics and tumor-targeting properties of the DOTA-TATE variants. Experimental details, including animal preparation, radiotracer administration, imaging parameters, and quantitative analysis methods, are provided in the Supplementary Information.

### Urine metabolite and enzyme-mediated cleavage analysis

To analyze radioactive urine metabolites, ^86^Y-DOTA-MVK(ε)-TATE or ^86^Y-DOTA-β-Gal-TATE (13 – 15 MBq in 200 μL PBS, 4 μg) was intravenously injected into normal, non-tumor-bearing athymic nu/nu mice (n = 3). As reference controls, standard metabolites of ^86^Y-DOTA-MVK(ε)-TATE (^86^Y-DOTA-G-M-OH) and ^86^Y-DOTA-β-Gal-TATE (^86^Y-DOTA-NHMe) were also prepared and administered to normal athymic, non-tumor-bearing nu/nu mice. Urine samples were collected 1 hour post-injection, centrifuged at 10,000 rpm, and subsequently analyzed by radio-HPLC for each ^86^Y-labeled DOTA-TATE variant. The radio-HPLC peaks were compared with those of the standard metabolites (^86^Y-DOTA-G-M-OH or ^86^Y-DOTA-NHMe). Radio-HPLC analysis was conducted under the same conditions as those used for analyzing the radiolabeled the DOTA-TATE variants, using an Agilent 1260 Infinity II system with a TSK-gel Octadecyl-4PW column and a gradient elution method. For *in vitro* enzyme-mediated cleavage, β-galactosidase was prepared in PBS (10 mM, pH 7.2) at 10 KU/mL. ^86^Y-DOTA-β-Gal-TATE (13– 15 MBq in 200 μL PBS) was mixed 1:1 (v/v) with β-galactosidase solution and incubated at 37 °C. Aliquots were taken at 1, 4, and 24 h, centrifuged at 10,000 rpm, and the supernatants were analyzed by radio-TLC. The 24 h sample was further examined by radio-HPLC to confirm metabolite identity.

### Targeted Alpha Particle Therapy, Response Monitoring, and Toxicity Assessment

For alpha particle therapy, athymic nu/nu mice bearing AR42J tumors (100 – 200 mm^3^) were prepared and randomly assigned to one of four treatment groups: vehicle (PBS), ^225^Ac-DOTA-TATE (148 kBq, 1 μg), ^225^Ac-DOTA-MVK(ε)-TATE (148 kBq, 1 μg), and ^225^Ac-DOTA-β-Gal-TATE (148 kBq, 1 μg). Mice received either a one-cycle regimen (n=8 per cohort) or a two-cycle regimen (n=6 per cohort) at an 8-day interval. Tumor volume and body weight were measured twice weekly after treatment using a digital caliper. Tumor volume was calculated using the formula V (mm^3^) = (l × w^2^)/2, where **l** (length) was the larger of the two perpendicular tumor axes, and **w** (width) was the smaller axis. Survival was monitored, with the endpoint defined as a tumor volume exceeding 2000 mm^3^, a weight loss greater than 20% from baseline, or signs of distress in the mice.

Estimated absorbed doses for individual organs for ^225^Ac- and ^86^Y-labeled DOTA-TATE variants were derived from residence times calculated using %IA/g data from biodistribution studies. The time–activity curves incorporated data from both radionuclides to provide complete coverage of early and late time points. Details of the dosimetry methodology, including time– activity curve construction, integration approach, decay assumptions, and conversion to effective dose, are provided in the Supplementary Information.

Additionally, a basic metabolic panel (BMP) and complete blood count (CBC) analysis were performed to monitor hematological parameters and potential treatment-related toxicity. For the one-cycle therapy cohort, blood was collected at 0-, 10-, and 31-days post-treatment, whereas for the two-cycle therapy cohort, blood was collected at 10 days, 3 months, and 6 months post-treatment. Samples were obtained retro-orbitally into potassium EDTA-coated tubes and analyzed using the Vetscan VS2 (Abaxis, Union City, CA, USA) and VetScan HM5 hematology analyzer (Abaxis). Although these instruments provide species-specific reference intervals, manufacturer-supplied ranges may not fully reflect our study cohort. Therefore, to ensure appropriate comparisons over the extended monitoring period, baseline and longitudinal reference values were established from age-matched normal mice (n = 5) using the same instrumentation and protocols. The following parameters were assessed and compared: alanine transaminase (ALT, U/L), total bilirubin (TBIL, µmol/L), creatinine (CRE, µmol/L), blood urea nitrogen (BUN, mmol/L), total white blood cell count (WBC, 10^9^/L), hemoglobin (HGB, g/dL), and platelets (PLT, 10^9^/L).

At the conclusion of the two-cycle therapy study, kidneys, livers, spleens, and tumors from representative mice were collected and processed for histological evaluation using hematoxylin and eosin (H&E) staining. Microscopic pathology was assessed and recorded in the MHL Pathology Database (Zaphod) using a standardized five-point ordinal grading system (normal, minimal, mild, moderate, marked). Terminology and diagnostic criteria were consistent with the International Harmonization of Nomenclature and Diagnostic Criteria (INHAND) guidelines established by the Society of Toxicologic Pathology (https://www.toxpath.org/inhand.asp)*(57)*.

### Statistical Analyses

Biodistribution, PET quantification, and hematological data were analyzed using the Student’s *t*-test for unpaired data to assess differences between the mean values of two groups. Statistical significance was defined as \**p* < 0.05, *\*\*p < 0*.*01, ***p < 0*.*001*, and \*\*\*\**p* < 0.0001. Survival curves were generated using the Kaplan–Meier method and compared with the log-rank (Mantel–Cox) test as the primary statistical approach. The Gehan–Breslow–Wilcoxon test was applied as a secondary analysis to account for early differences in survival. All statistical analyses were performed using GraphPad Prism, version 10.3.1 (GraphPad Software, San Diego, CA, USA).

## Acknowledgments

This research was fully supported by the Intramural Research Program of the National Cancer Institute, National Institutes of Health (Molecular Imaging Branch, Center for Cancer Research). The contributions of the NIH authors were made as part of their official duties as NIH federal employees, are in compliance with agency policy requirements, and are considered Works of the United States Government. However, the findings and conclusions presented in this paper are those of the authors and do not necessarily reflect the views of the NIH or the U.S. Department of Health and Human Services. We also acknowledge the NIH Cyclotron Facility for the production and supply of the radioisotopes used in this study.

## Funding

This work was supported by the Intramural Research Program of the National Institutes of Health, including the National Cancer Institute, Center for Cancer Research, and the Molecular Imaging Branch. The projects funded under this support are ZIA BC011800, ZIA BC 010891, ZIA BC011506, and ZIA BC011564. This project has been funded in whole or in part with Federal funds from the National Cancer Institute, National Institute of Health, under Contract No. HHSN26120150003I. The contributions of the NIH authors are considered Works of the United States Government. The findings and conclusions presented in this paper are those of the authors and do not necessarily reflect the views or policies of the NIH or the U.S. Department of Health and Human Services, and mention of trade names, commercial products, or organizations does not imply endorsement by the U.S. Government.

## Author contributions

Conceptualization: M.J.S., F.E.E.

Methodology: W.L., S.R.D., K.E.B., E.L., R.S., M.J.S., F.E.E.

Investigation: W.L., K.E.B., D.N., Y.U., S.O., J.Y.C., X.L., H.M., J.S.K., S.F.

Visualization: W.L., J.Y.C., S.A., E.E., M.J.S.

Funding acquisition: M.J.S., F.E.E.

Project administration: W.L., K.E.B., S.R.D., E.L., J.A.B., A.K.L.

Supervision: M.J.S., F.E.E., R.S., O.J.W., F.I.L., P.L.C.

Writing – original draft: W.L., M.J.S., F.E.E.

Writing – review & editing: W.L., S.R.D., K.E.B., D.N., J.Y.C, S.A., E.E., Y.U., S.O., X.L.,

H.M., J.S.K., S.F., O.J.W., E.L., J.A.B., A.K.L., F.I.L., P.L.C., R.S., M.J.S., F.E.E.

## Competing interests

Authors declare that they have no competing interests.

## Data and materials availability

All data are available in the main text or the supplementary materials.

## Notes

### Competing Interest Statement

The authors have declared no competing interest.

