## Supporting Information for "Biodistribution-Driven Discovery Identifies a Glycosidase-Cleavable Linker to Reprogram Radiotheranostics"

### Synthesis and Characterization of DOTA-TATE Variants

**General Methods:** Starting materials were used as received unless otherwise noted. All moisture sensitive reactions were performed in an inert atmosphere of argon with oven dried glassware. Reagent grade solvents were used for extractions and flash chromatography. Reaction progress was monitored by LC/MS analysis performed on an Agilent UPLC/MS instrument equipped with a RP-C18 column (Poroshell 120 SB-C18, 4.6 3 50 mm, 2.7 mm or Zorbax 300SB-C18, 4.6 3 50 mm, 3.5 mm), dual atmospheric pressure chemical ionization (APCI)/electrospray (ESI) mass spectrometry detector, and photodiode array detector. Flash chromatography was performed by using a RediSepRf NP-silica (40–63 mm 60 A˚) or a Teledyne RediSepRf Gold RP-C18 column (20–40 mm 100 A˚) in a Teledyne ISCO CombiFlash Rf 200 purification system unless otherwise specified. Fmoc-amino acids were procured from AAPTEC and Chem-Impex Intl. Bio-grade solvents were purchased from Oakwood chemicals. Reagents used for couplings and deprotection were obtained from Chem-Impex Intl. Fmoc-serine preloaded Rink amide resin was purchased from AAPTEC. Automated synthesis of peptides was carried out on an AAPTEC Focus-6V synthesizer employing standard Fmoc chemistry. LC/MS were recorded on an Agilent Technologies 1290 Infinity LC/XT MSD instrument.

DMF – Dimethylformamide

DCM – Dichloromethane

TIPS – Triisopropylsilane

HBTU – 2-(1*H*-benzotriazol-1-yl)-1,1,3,3-tetramethyluronium hexafluorophosphate

NMM – 4-Methylmorpholine

TFA – Trifluroacetic acid

**General Procedure for Solid Phase Peptide synthesis:**

**Deprotection**: The Fmoc group from amino acids were removed by shaking the resin with 20% piperidine in DMF for 15 min (twice; 15.0 mL/g of the resin). The resin was drained, washed with DMF (three times; 15.0 mL/g of the resin) and was taken to the next step.

**Washing:** The resin was drained from the deprotection/coupling mixture, and washed with DMF three times (15.0 mL/g) and after the final Fmoc-deprotection, the resin was washed with DCM three times

**Coupling:** 8.0 Equiv. of the amino acid to be coupled, was activated with 8.0 equiv. of HBTU, and 16.0 equiv. of NMM in a mixing vessel, and thoroughly mixed for 2.0 min by bubbling nitrogen through the solution. This activated amino acid was then transferred to the resin in the reaction vessel under a positive pressure of nitrogen, and the vessel was shaken for 1 h while nitrogen gas was bubbled through the solution every 5 min. After 1 h of coupling time, the resin was drained, washed with DMF (thrice) and, used for the next step

**DOTA coupling:** 8.0 Equiv. of the DOTA-Tris(tBu) to be coupled, was activated with 8.0 equiv. of HBTU, and 16.0 equiv. of NMM in a mixing vessel, and thoroughly mixed for 2.0 min by bubbling nitrogen through the solution. This activated DOTA-Tris(tBu) acid was then transferred to the resin in the reaction vessel under a positive pressure of nitrogen, and the vessel was shaken for 1h while nitrogen gas was bubbled through the solution every 5 min. After 1h of coupling time, the resin was drained, washed with DMF (3x) and used for the next step deprotection.

**Peptide cleavage from the resin**: A cleavage cocktail of 98:1:1 - TFA:TIPS:water was added to the peptide vessel with the resin (15.0 mL/g) and shaken for 1 h. The resin was filtered and washed with 2 x 10.0 mL of cleavage cocktail and the filtrates were combined and evaporated under reduced pressure at room temperature (RT). The residue was triturated with anhydrous ether (20.0 mL) and decanted. This process was repeated two more times and the fully deprotected peptide residues were purified by prep. HPLC. The syntheses started with 0.1 mmol of O-*t*-Butyl-L-threonine-2-chlorotrityl resin. After the cleavage, the crude peptides were taken to the next S-S cyclization step without further purification.

**Intramolecular disulfide formation:** The crude peptide was dissolved in 10% DMSO of 0.001 M $NH_4HCO_3$ (100 mL per 100 mg of crude peptide) and stirred at RT under open atmosphere for 2 days. After completion of the reaction, the solution was freeze-dried and purified and used for the next step without further purification.

**Preparative HPLC conditions**: Column: X-Terra® (Waters corp.) C18 RP; 150 x 30 mm; 5.0 microns; solvent A: water with 0.1% TFA (v/v) and solvent B: acetonitrile with 0.1% TFA (v/v); Elution rate: 40.0 mL/min; Gradient: 10% B – 65% B over 40 min; Detection @ 220 nm. Fractions with the required mass and purity of >95% were pooled and freeze-dried to yield the peptides as colorless fluffy solids.

Analytical HPLC conditions **A**: Column: Zorbax C18; 3.5 microns; 50 x 4.6 mm; Solvent A: Water with 0.1% TFA and B: acetonitrile with 0.1% TFA; Elution rate: 1.0 mL/min; 5% B – 95% B over 7 min; Detection @ 220 nm.

Analytical HPLC conditions **B**: Column: Zorbax C18; 2.7 microns; 50 x 4.6 mm; Solvent A: Water with 0.1% TFA and B: acetonitrile with 0.1% TFA; Elution rate: 1.0 mL/min; 5% B – 95% B over 5 min; Detection @ 220 nm.

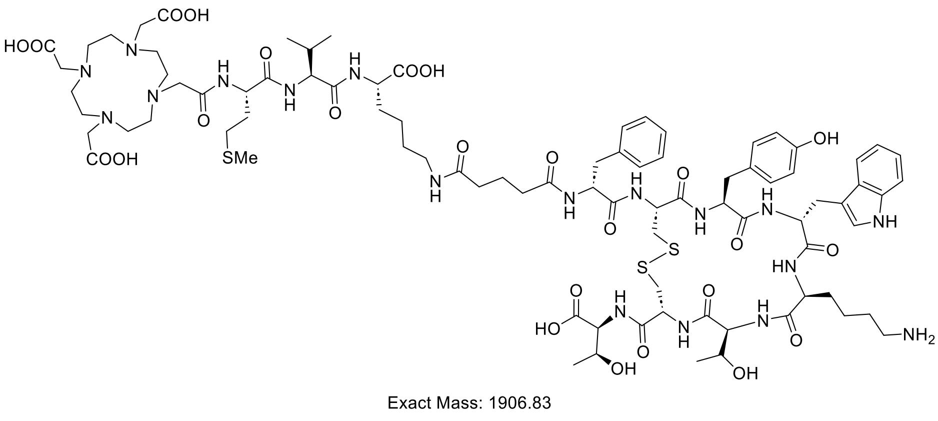


**DOTA-MVK(Ɛ)-TATE:** Synthesized following the general procedure.

LC-MS (ESI): (m/z) = 1907 [M+H]

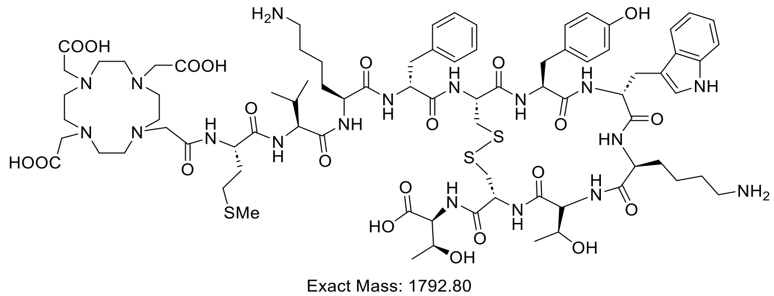


**DOTA-MVK-TATE:** Synthesized following the general procedure.

LC-MS (ESI): (m/z) = 1793 [M+H]

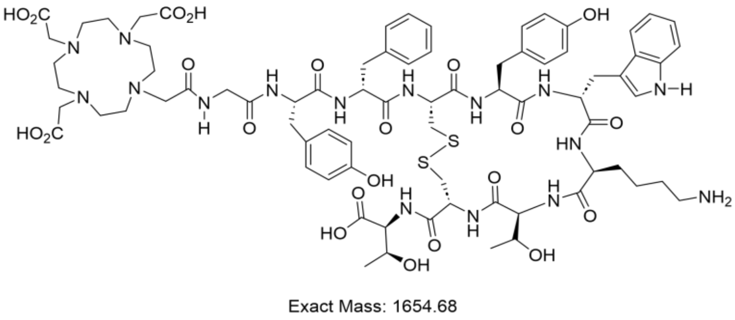


**DOTA-GY-TATE:** Synthesized following the general procedure.

LC-MS (ESI): (m/z) = 1655 [M+H]

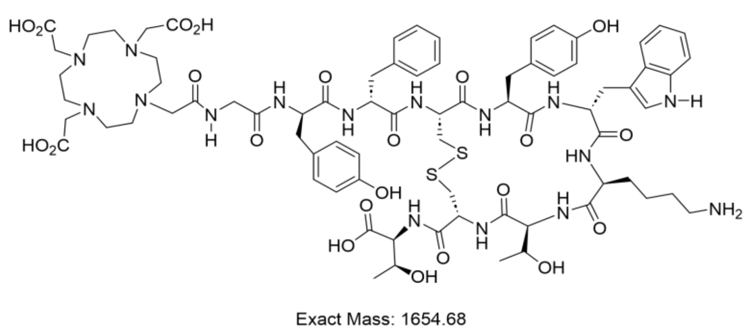


**DOTA-GY*-TATE:** Synthesized following the general procedure.

LC-MS (ESI): (m/z) = 1655 [M+H]

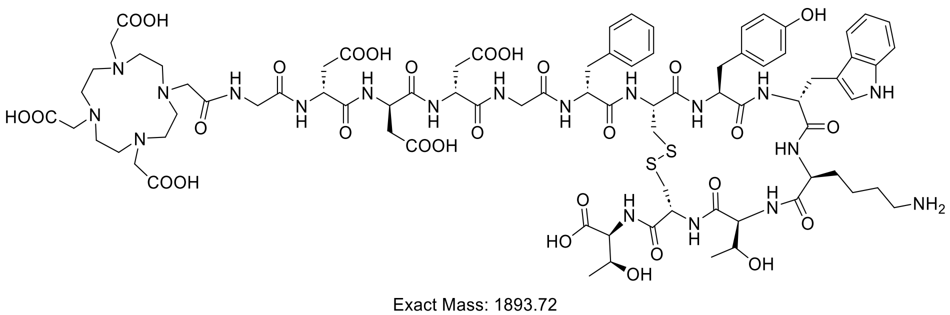


**DOTA-GDDDG-TATE:** Synthesized following the general procedure.

LC-MS (ESI): (m/z) = 1895 [M+H]

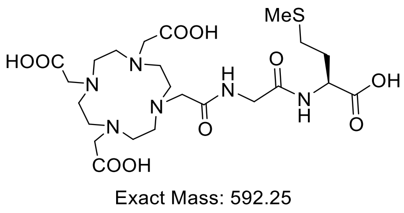


**DOTA-G-M-OH:** Synthesized following the general procedure.

LC-MS (ESI): (m/z) = 593 [M+H]

**DOTA-β-Gal-TATE Synthesis:**

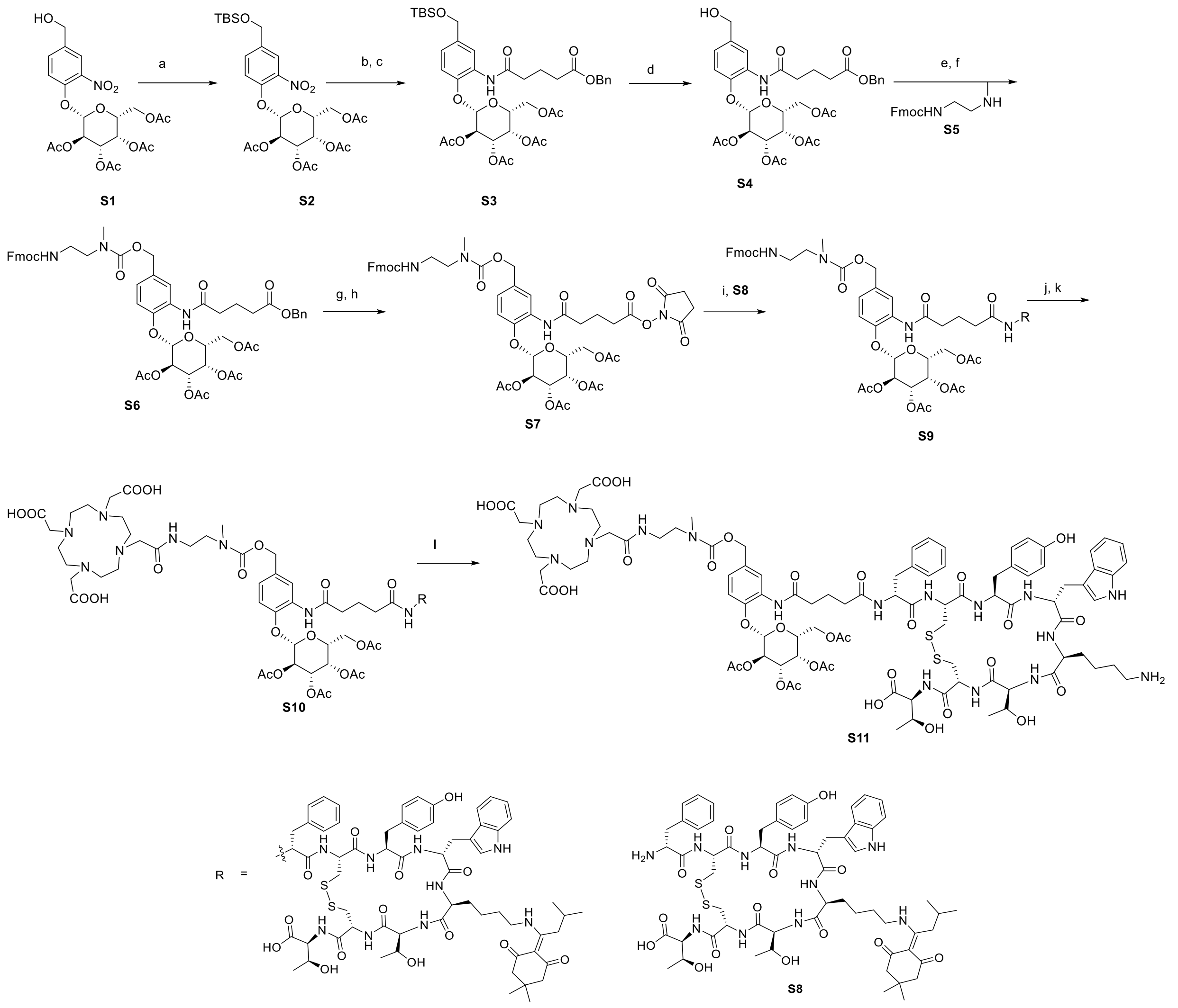

**Scheme S1.** *Reagents and conditions*: a) Imidazole, TBDMSCl, DCM, RT, overnight, 90%; b) Zinc, $NH_4Cl$, MeOH, 0 °C, 2 h, quantitative; c) 5-(benzyloxy)-5-oxopentanoic acid, HATU, DIPEA, DMF, RT, overnight, 85%; d) TsOH, MeOH, RT, 30 min, 95%; e) 4-nitrophenyl chloroformate, pyridine, DCM, 0 °C - RT, 2 h, quantitative; f) **5**, DIPEA, DMF, RT, 30 min, 71%; g) Pd/C (10 wt. %), $H_2$, MeOH, RT, 4 h, quantitative; h) TSTU, DIPEA, DMF, RT, 1 h, 85%; i) **S8**, 1:1 ratio of acetonitrile and 0.1M phosphate buffer (pH = 7.3), RT, 16 h, 65%; j) piperidine, DMF, RT, 30 min, quantitative; k) DOTA-NHS-ester**,** DIPEA, DMF, RT, 3 h, 60%; l) hydrazine hydrate, DMF, RT, 30 min, 45%.

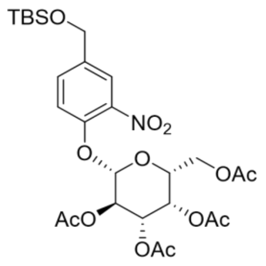

Exact Mass: 613.22

**(2R,3S,4S,5R,6S)-2-(acetoxymethyl)-6-(4-(hydroxymethyl)-2-nitrophenoxy)tetrahydro-2H-pyran-3,4,5-triyl triacetate (S2):** The (2R,3S,4S,5R,6S)-2-(acetoxymethyl)-6-(4-(hydroxy methyl)-2-nitrophenoxy)tetrahydro-2H-pyran-3,4,5-triyl triacetate **S1***(58)* (1.0 g, 2.002 mmol) was dissolved in DCM (30 mL), was added sequentially TBS-Cl (452.7 mg, 3.003 mmol) and imidazole (204.5 mg, 3.003 mmol) at 0 °C. The resulting reaction stirred overnight at RT. After completion, added water to the reaction mixture and extracted with DCM (2 x 100 mL). The combined organic layers were washed with brine solution (20 mL) and concentrated to give crude product. The crude product was purified by flash chromatography on silica gel eluting with ethyl acetate in hexane (0 to 50% gradient) to give the title compound as a white solid (1.1 g, 1.8 mmol, 90%).

$^{1}$H NMR (500 MHz, $CDCl_3$) δ 8.25 (d, *J* = 1.9 Hz, 1H), 7.79 (s, 1H), 7.30 – 7.19 (m, 8H), 6.96 (dd, *J* = 8.3, 2.0 Hz, 1H), 6.84 (d, *J* = 8.4 Hz, 1H), 5.42 – 5.34 (m, 2H), 5.09 (dd, *J* = 10.6, 3.4 Hz, 1H), 5.03 (d, *J* = 6.4 Hz, 3H), 4.93 (d, *J* = 7.8 Hz, 1H), 4.59 (s, 2H), 4.16 (dd, *J* = 11.2, 7.0 Hz, 1H), 4.09 – 3.99 (m, 2H), 2.42 (q, *J* = 6.8 Hz, 4H), 2.34 (dt, *J* = 16.0, 7.3 Hz, 3H), 2.09 (s, 3H), 2.04 – 1.92 (m, 11H), 1.88 (p, *J* = 7.4 Hz, 2H), 0.84 (s, 9H), -0.00 (s, 6H) ppm.

$^{13}$C NMR (126 MHz, $CDCl_3$) δ 177.4, 172.9, 172.7, 170.9, 170.8, 170.4, 170.2, 170.0, 143.9, 136.9, 136.0, 135.9, 128.6, 128.6, 128.3, 128.2, 128.2, 121.1, 118.0, 113.1, 99.9, 77.4, 77.3, 77.1, 76.8, 71.1, 70.1, 69.2, 66.7, 66.3, 66.2, 64.7, 61.3, 36.4, 33.5, 33.2, 32.8, 26.0, 21.0, 20.8, 20.6, 20.6, 20.6, 19.9, 18.4, -5.2 ppm.

LC-MS (ESI): (m/z) = 614 [M+H]

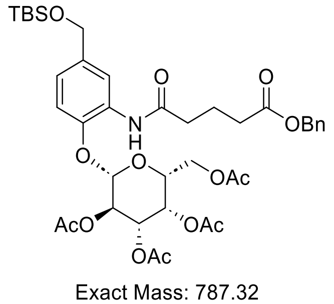

Exact Mass: 787.32

**(2R,3S,4S,5R,6S)-2-(acetoxymethyl)-6-(2-(5-(benzyloxy)-5-oxopentanamido)-4-(((tert-butyldimethylsilyl)oxy)methyl)phenoxy)tetrahydro-2H-pyran-3,4,5-triyl triacetate (S3):**

**S2** (1.1 g, 1.8 mmol) was taken up in 40 mL methanol and cooled to 0 °C in ice bath zinc (1.2 g, 18 mmol) and ammonium chloride (0.96 g, 18 mmol) were added sequentially. The reaction was stirred on ice for 15 min, then the ice bath was removed, and stirring was continued at RT for 2 h. The reaction mixture was filtered through celite with methanol, and the filtrate was concentrated in vacuo. The crude residue was re-suspended in DCM (100 mL) and saturated $NaHCO_3$ solution (50 mL). The layers were separated and the aqueous layers were extracted with DCM (2 x 100 mL). The combined organic layers dried over sodium sulfate and concentrated in vacuo to give crude compound, the crude product was used for next step without further purification.

To a stirred solution of the above crude product (1.8 mmol) in DMF (10 mL), was added sequentially 5-(benzyloxy)-5-oxopentanoic acid (480 mg, 2.16 mmol), HATU (1.026 g, 2.7 mmol), and DIPEA (0.64 mL 3.6 mmol) at 0 °C. The resulting mixture was stirred at RT for overnight. After completion of the reaction, added water to the reaction mixture at 0 °C and extracted with ethyl acetate (3 x 150 mL). The combined organics were washed with water (50 mL), brine solution (50 mL). The combined organic layers dried over sodium sulfate and concentrated in vacuo to give crude compound. The crude product was purified by flash chromatography on silica gel eluting with ethyl acetate in hexane (0 to 100% gradient) to give the title compound as a white solid (1.2 g, 1.53 mmol, 85%).

$^1$H NMR (500 MHz, $CDCl_3$) δ 8.26 (s, 1H), 7.79 (s, 1H), 7.27 – 7.17 (m, 5H), 6.96 (d, $J$ = 8.4 Hz, 1H), 6.85 (d, $J$ = 8.4 Hz, 1H), 5.42 – 5.35 (m, 2H), 5.09 (dd, $J$ = 10.6, 3.4 Hz, 1H), 5.03 (s, 2H), 4.94 (d, $J$ = 7.8 Hz, 1H), 4.59 (s, 2H), 4.16 (dd, $J$ = 11.2, 6.9 Hz, 1H), 4.11 – 4.00 (m, 2H), 2.42 (td, $J$ = 7.4, 4.4 Hz, 4H), 2.08 (d, $J$ = 1.2 Hz, 3H), 2.03 – 1.90 (m, 12H), 0.84 (t, $J$ = 1.3 Hz, 10H), 0.00 (d, $J$ = 1.4 Hz, 6H) ppm.

$^{13}$C NMR (126 MHz, $CDCl_3$) δ 172.9, 170.8, 170.8, 170.3, 170.1, 169.9, 143.9, 136.9, 136.0, 128.5, 128.3, 128.2, 121.0, 117.9, 113.1, 99.9, 77.4, 77.2, 76.9, 71.1, 70.1, 69.2, 66.7, 66.2, 64.7, 61.3, 36.4, 33.5, 26.0, 21.0, 20.8, 20.6, 20.5, 18.4, -5.2 ppm.

LC-MS (ESI): (m/z) = 788 [M+H]

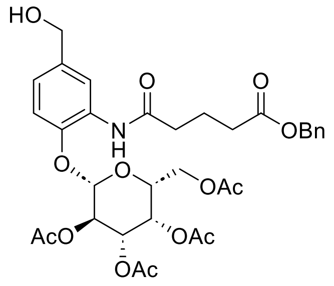

Exact Mass: 673.24

**(2R,3S,4S,5R,6S)-2-(acetoxymethyl)-6-(2-(5-(benzyloxy)-5-oxopentanamido)-4-(hydroxymethyl)phenoxy)tetrahydro-2H-pyran-3,4,5-triyl triacetate (S4):** To a solution of **S3** (1.1 g, 1.396 mmol) in MeOH (10 mL), was added p-toluenesulfonic acid monohydrate (26.55 mg, 0.1396 mmol) at RT. The resulting solution was stirred at RT for 30 min. After completion of the reaction as indicated by LCMS, the reaction mixture was neutralized by addition of $Et_3N$ (0.1 mL) at 0 °C. The solution was concentrated to give crude product and the crude product was purified by flash chromatography on silica gel eluting with ethyl acetate in hexane (0 to 100% gradient) to give the title compound as a thick oil (0.89 g, 1.32 mmol, 95%).

$^{1}$H NMR (500 MHz, $CDCl_3$) δ 8.31 (dd, *J* = 5.5, 2.0 Hz, 1H), 7.83 (s, 1H), 7.31 – 7.21 (m, 5H), 6.97 (dt, *J* = 8.4, 2.5 Hz, 1H), 6.87 (dd, *J* = 8.4, 1.8 Hz, 1H), 5.43 – 5.35 (m, 2H), 5.12 (dd, *J* = 10.6, 3.4 Hz, 1H), 5.04 (d, *J* = 3.6 Hz, 2H), 4.98 (d, *J* = 7.8 Hz, 1H), 4.51 (d, *J* = 5.4 Hz, 2H), 4.20 – 3.98 (m, 4H), 2.47 – 2.38 (m, 4H), 2.09 (d, *J* = 3.8 Hz, 3H), 2.05 – 1.91 (m, 12H) ppm.

$^{13}$C NMR (126 MHz, $CDCl_3$) δ 172.9, 171.0, 170.9, 170.4, 170.2, 169.9, 144.3, 136.6, 136.0, 122.0, 119.0, 113.3, 99.8, 99.8, 77.4, 77.4, 77.2, 77.1, 76.9, 76.9, 71.1, 71.1, 70.1, 69.2, 66.7, 66.2, 64.7, 61.3, 60.4, 36.3, 33.5, 21.0, 21.0, 20.7, 20.6, 20.6, 20.5, 14.2 ppm.

LC-MS (ESI): (m/z) = 674 [M+H]

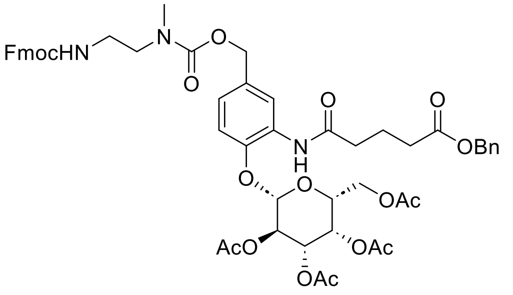


Exact Mass: 995.37

**(2S,3R,4S,5S,6R)-2-(4-(10-(9H-fluoren-9-yl)-4-methyl-3,8-dioxo-2,9-dioxa-4,7-diazadecyl)-2-(5-(benzyloxy)-5-oxopentanamido)phenoxy)-6-(acetoxymethyl)tetrahydro-2H-pyran-3,4,5-triyl triacetate (S6):** To a solution of **S4** (0.6 g, 0.89 mmol) in anhydrous DCM (5 mL) cooled at 0 °C, was sequentially added dropwise pyridine (179 µL, 2.22 mmol) and 4-nitrophenyl chloroformate (360 mg, 1.78 mmol) dissolved in anhydrous DCM (5 mL). The mixture was stirred at RT under argon atmosphere for 2 h, then quenched with saturated solution of $NaHCO_3$ (20 mL) and extracted with DCM (3 x 25 mL). The combined organic layers were dried over anhydrous sodium sulfate, filtered and concentrated under reduced pressure to give crude product. The crude product was purified by flash chromatography on silica gel eluting with ethyl acetate in DCM (0 to 50% gradient) to give activated carbonate as a quantitative yield.

The resulting carbonate (0.89 mmol) was dissolved in DMF (5 mL), was added subsequently (9H-fluoren-9-yl)methyl (2-(methylamino)ethyl)carbamate (**S5**), HCl (395 mg, 1.33 mmol) and DIPEA (0.3 mL, 1.78 mmol) at RT under argon atmosphere. The reaction was stirred for 30 min at RT. After completion of the reaction (as indicated by LCMS) crude material was purified by reverse phase flash chromatography without workup using a gradient of 10-85% acetonitrile in water with the addition of 0.05% trifluoroacetic acid to get the corresponding product (628 mg, 0.63 mmol, 71%) as a white solid.

$^{1}$H NMR (500 MHz, $CDCl_3$) δ 8.49 – 8.40 (m, 1H), 7.81 (d, *J* = 6.0 Hz, 1H), 7.67 (d, *J* = 7.6 Hz, 1H), 7.48 (d, *J* = 8.0 Hz, 2H), 7.33 – 7.14 (m, 8H), 6.95 (dt, *J* = 8.5, 1.9 Hz, 1H), 6.90 – 6.74 (m, 2H), 5.42 – 5.30 (m, 3H), 5.20 (d, *J* = 1.5 Hz, 3H), 5.10 – 4.91 (m, 6H), 4.45 (d, *J* = 1.6 Hz, 2H), 4.24 (d, *J* = 7.1 Hz, 1H), 4.18 – 4.00 (m, 4H), 3.33 (dq, *J* = 36.4, 7.1 Hz, 2H), 2.88 (s, 1H), 2.47 – 2.36 (m, 4H), 2.11 – 2.05 (m, 12H), 2.02 – 1.92 (m, 14H) ppm.

$^{13}C$ NMR (126 MHz, $CDCl_3$) δ 206.9, 172.9, 171.0, 170.9, 170.3, 170.3, 170.1, 169.9, 144.7, 144.0, 141.3, 136.0, 132.9, 128.7, 128.6, 128.5, 128.2, 128.2, 127.7, 127.1, 123.6, 120.4, 120.0, 113.2, 99.6, 71.2, 70.0, 69.1, 66.6, 66.2, 66.2, 61.3, 61.3, 53.5, 46.1, 36.3, 33.5, 30.9, 21.0, 21.0, 20.7, 20.6, 20.6, 20.6, 20.6 ppm.

LC-MS (ESI): (m/z) = 996 [M+H]

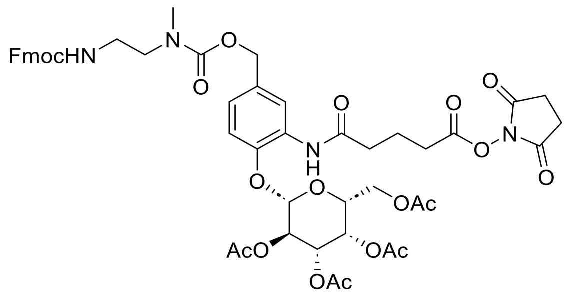


Exact Mass: 1002.34

**(2S,3R,4S,5S,6R)-2-(4-(10-(9H-fluoren-9-yl)-4-methyl-3,8-dioxo-2,9-dioxa-4,7-diazadecyl)-2-(5-((2,5-dioxopyrrolidin-1-yl)oxy)-5-oxopentanamido)phenoxy)-6-(acetoxymethyl) tetrahydro-2H-pyran-3,4,5-triyl triacetate (S7):** To a stirred solution of the compound **S6** (320 mg, 0.35 mmol) in MeOH (10 mL), was added Pd/C (64 mg, 10%w/w) at RT. The resulting solution was stirred at RT for 4 h under hydrogen pressure (hydrogen balloon was used for hydrogen pressure). After complete consumption of starting compound, the reaction mixture was filtered to remove the solids, and filtrate was concentrated to get the crude product which was pure enough and used for the next step without purification.

The resulting acid (0.35 mmol) was dissolved in DMF (3 mL), was added sequentially TSTU (160 mg, 0.53 mmol) and DIPEA (0.12 mL, 0.7 mmol) at 0 °C. The resulting solution was stirred at RT for 1 h. After completion of the reaction (as indicated by LCMS), the crude material was purified by reverse phase flash chromatography without workup using a gradient of 10-75% acetonitrile in water with the addition of 0.05% TFA to get the corresponding product (300 mg, 0.29 mmol, 85%).

$^{1}H$ NMR (500 MHz, DMSO) δ 8.60 (s, 1H), 7.86 – 7.72 (m, 3H), 7.61 (d, $J$ = 6.6 Hz, 2H), 7.32 (dt, $J$ = 11.7, 6.7 Hz, 3H), 7.25 (t, $J$ = 7.4 Hz, 2H), 7.09 – 6.94 (m, 2H), 5.38 – 5.31 (m, 1H), 5.29 (s, 1H), 5.26 – 5.13 (m, 2H), 4.88 (d, $J$ = 8.3 Hz, 2H), 4.41 – 4.32 (m, 1H), 4.23 (d, $J$ = 6.8 Hz, 2H), 4.13 (t, $J$ = 7.7 Hz, 1H), 4.03 (q, $J$ = 6.8 Hz, 2H), 3.19 (d, $J$ = 6.8 Hz, 2H), 3.05 (q, $J$ = 6.3 Hz, 2H), 2.76 (d, $J$ = 10.6 Hz, 7H), 2.68 (d, $J$ = 6.9 Hz, 2H), 2.38 (s, 2H), 2.07 (s, 3H), 2.02 – 1.82 (m, 12H) ppm.

LC-MS (ESI): (m/z) = 1003 [M+H]

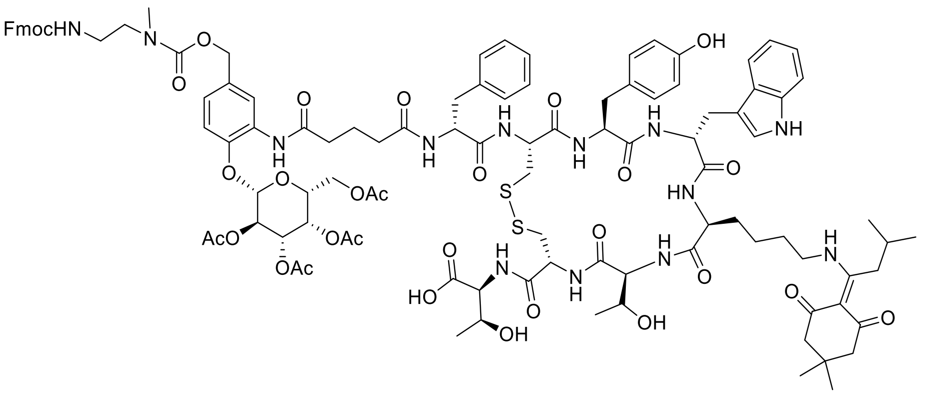


Exact Mass: 2141.86

**((4R,7S,10S,13R,16S,19R)-13-((1H-indol-3-yl)methyl)-19-((R)-2-(5-((5-(10-(9H-fluoren-9-yl)-4-methyl-3,8-dioxo-2,9-dioxa-4,7-diazadecyl)-2-(((2S,3R,4S,5S,6R)-3,4,5-triacetoxy-6-(acetoxymethyl)tetrahydro-2H-pyran-2-yl)oxy)phenyl)amino)-5-oxopentanamido)-3-phenylpropanamido)-10-(4-((1-(4,4-dimethyl-2,6-dioxocyclohexylidene)-3-methylbutyl) amino)butyl)-16-(4-hydroxybenzyl)-7-((R)-1-hydroxyethyl)-6,9,12,15,18-pentaoxo-1,2-dithia-5,8,11,14,17-pentaazacycloicosane-4-carbonyl)-L-allothreonine (S9):** To a solution of compound **S7** (55 mg, 0.05489 mmol) in acetonitrile and 0.1M phosphate buffer (pH = 7.3), (2 mL, 1:1 v/v), was added **S8***(59)* (75 mg, 0.06 mmol) at 0 °C. The resulting solution was stirred at RT for 16 h. After completion of the reaction (as indicated by LCMS), the crude material was purified by reverse phase chromatography without workup using a gradient of 10-65% acetonitrile in water with the addition of 0.05% TFA to get the corresponding product (75 mg, 0.035 mmol, 65%).

LC-MS (ESI): (m/z) = 2143 [M+H]

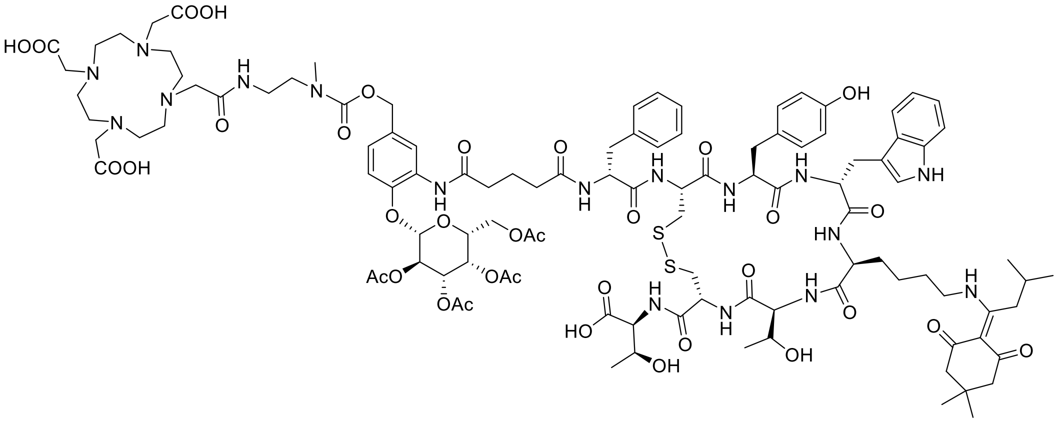


Exact Mass: 2305.97

**2,2',2''-(10-(2-((2-((((3-(5-(((R)-1-(((4R,7S,10S,13R,16S,19R)-13-((1H-indol-3-yl)methyl)-4-(((1S,2S)-1-carboxy-2-hydroxypropyl)carbamoyl)-10-(4-((1-(4,4-dimethyl-2,6-dioxocyclohexylidene)-3-methylbutyl)amino)butyl)-16-(4-hydroxybenzyl)-7-((R)-1-hydroxyethyl)-6,9,12,15,18-pentaoxo-1,2-dithia-5,8,11,14,17-pentaazacycloicosan-19-yl)amino)-1-oxo-3-phenylpropan-2-yl)amino)-5-oxopentanamido)-4-(((2S,3R,4S,5S,6R)-**

**3,4,5-triacetoxy-6-(acetoxymethyl)tetrahydro-2H-pyran-2-yl)oxy)benzyl)oxy)carbonyl)(methyl)amino)ethyl)amino)-2-oxoethyl)-1,4,7,10-tetraazacyclododecane-1,4,7-triyl) triacetic acid (S10):** Compound **S9** (30 mg, 0.014. mmol) was dissolved in 20% piperidine in DMF, and the solution was stirred for 30 min at RT. After completion of the reaction (as indicated by LCMS), the crude material was purified by reverse phase chromatography without workup using a gradient of 10-50% acetonitrile in water with the addition of 0.05% TFA to get the primary amine (**S10a**), which was used for the next step without further analysis. The resulting amine (0.014 mmol) was dissolved in DMF (1 mL), was added 2,2',2''-(10-(2-((2,5-dioxopyrrolidin-1-yl)oxy)-2-oxoethyl)-1,4,7,10-tetraazacyclododecane-1,4,7-triyl)triacetic acid (21 mg, 0.042 mmol) and DIPEA (12.2 µL, 0.07 mmol) at 0 °C. The resulting solution was stirred at RT for 3 h. After completion of the reaction (as indicated by LCMS) crude material was purified by reverse phase chromatography without workup using a gradient of 10-55% acetonitrile in water with the addition of 0.05% TFA to get the corresponding product (19 mg, 0.0082 mmol, 60%).

LC-MS (ESI): (m/z) = 2307 [M+H]

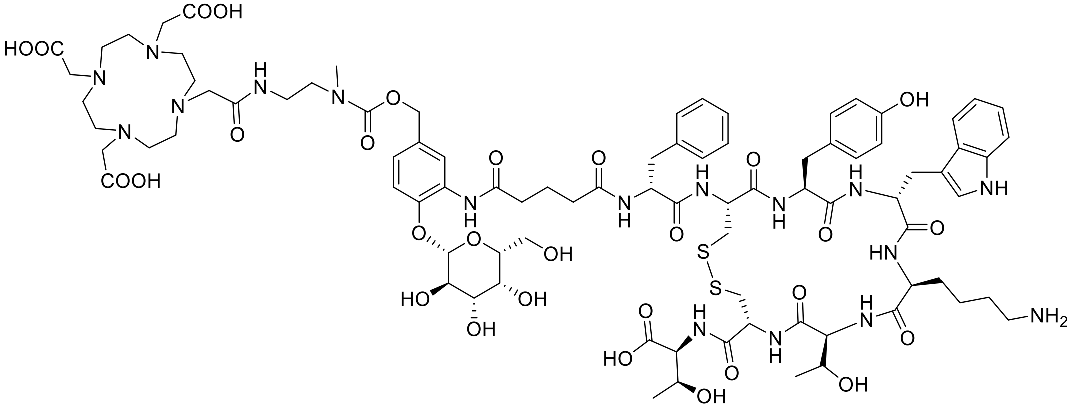


Exact Mass: 1931.80

**DOTA-β-Gal-TATE (S11):** Compound **S10** (15 mg, 0.0065 mmol) was dissolved in 10% hydrazine hydrate in DMF and the resulting solution was stirred at RT for 30 min. After completion of the reaction (as indicated by LCMS) crude solution was purified directly without workup by reverse phase HPLC (HPLC conditions: Column: X-Terra® (Waters corp.) C18 RP; 150 x 30 mm; 5.0 microns; solvent A: water with 0.1% TFA (v/v) and solvent B: acetonitrile with 0.1% TFA (v/v); Elution rate: 50.0 mL/min; Gradient: 10%B – 65%B over 35 min; Detection @ 220 nm. Fractions with the required mass and purity of >95% were pooled and freeze-dried to yield the peptides as colorless fluffy solids) to get the corresponding product (6.1 mg, 0.0029 mmol, 45%).

LC-MS (ESI): (m/z) = 1932 [M+H]

**DOTA-β-Glu-TATE Synthesis:**

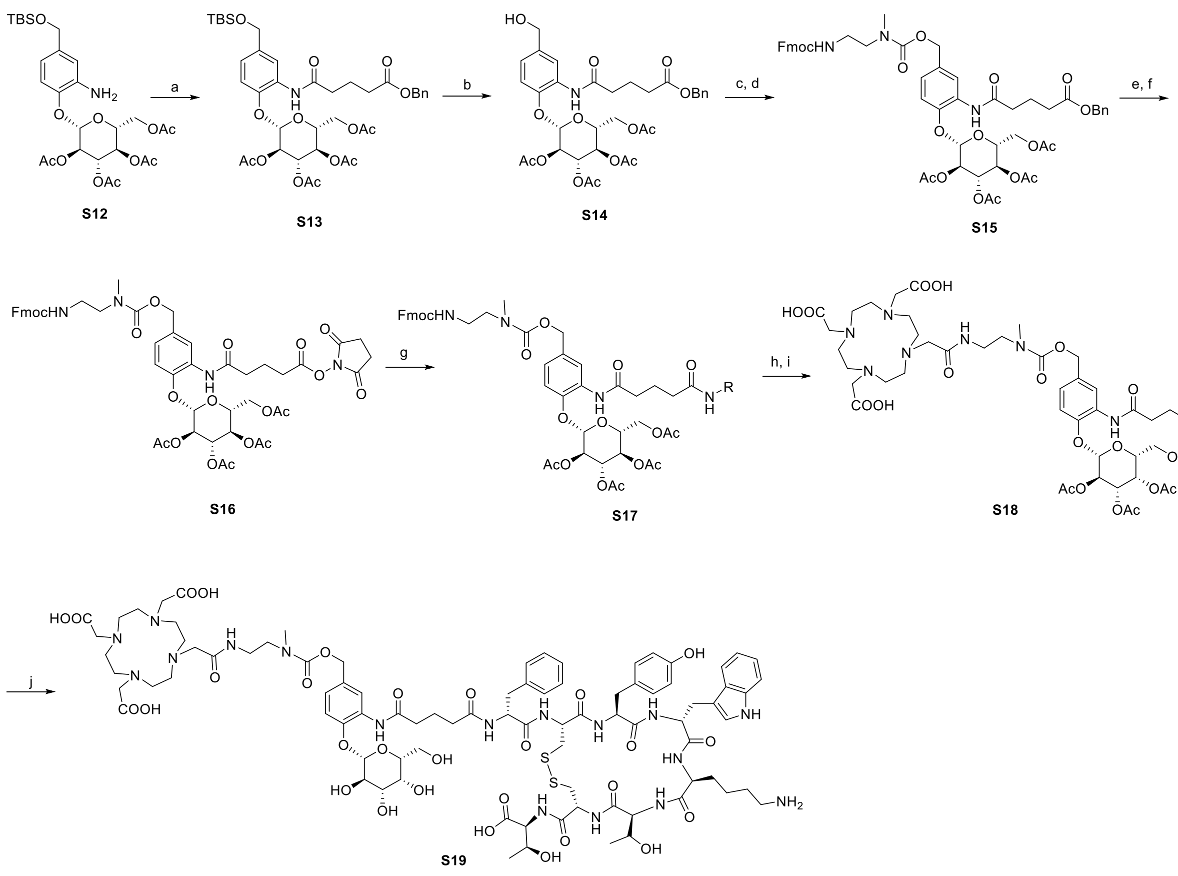

**Scheme S2.** *Reagents and conditions*: a) 5-(benzyloxy)-5-oxopentanoic acid, HATU, DIPEA, DMF, RT, overnight, 78%; b) TsOH, MeOH, RT, 30 min, 96%; c) 4-nitrophenyl chloroformate, pyridine, DCM, 0 °C -RT, 2 h, d) **S5**, DIPEA, DMF, RT, 30 min, 70%; e) Pd/C (10 wt. %), MeOH, $H_2$, RT, 1 h; f) TSTU, DIPEA, DMF, RT, 4 h, 81%; g) **S8**, acetonitrile and 0.1M phosphate buffer (pH = 7.3), RT, 20 h, 62%; h) piperidine, DMF, RT; i) DOTA-NHS-ester, DIPEA, DMF, RT, 2 h, 56%; j) Hydrazine hydrate, DMF, RT, 30 min, 40%.

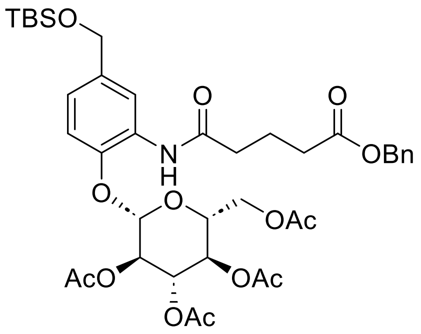


Exact Mass: 787.32

**(2R,3R,4S,5R,6S)-2-(acetoxymethyl)-6-(2-(5-(benzyloxy)-5-oxopentanamido)-4-(((tert-butyldimethylsilyl)oxy)methyl)phenoxy)tetrahydro-2H-pyran-3,4,5-triyl triacetate (S13):**

To a stirred solution of the (2R,3R,4S,5R,6S)-2-(acetoxymethyl)-6-(2-amino-4-(((tert-butyl dimethylsilyl)oxy)methyl)phenoxy)tetrahydro-2H-pyran-3,4,5-triyl triacetate **S12***(60)* (1.0 g, 1.71 mmol) in DMF (10 mL), was added sequentially 5-(benzyloxy)-5-oxopentanoic acid (488 mg, 2.06 mmol) HATU (1.026 g, 2.7 mmol), and DIPEA (0.64 mL 3.6 mmol) at 0 °C. The resulting mixture was stirred at RT for overnight. After completion of the reaction, added water to the reaction mixture at 0 °C and extracted with ethyl acetate (3 x 150 mL). The combined organics were washed with water (50 mL), brine solution (50 mL), and dried over anhydrous $Na_2SO_4$. The combined organic layers were concentrated in vacuo to give crude compound. The crude product was purified by flash chromatography on silica gel eluting with ethyl acetate in hexane (0 to 100% gradient) to give the title compound as a white solid (1.05 g, 1.33 mmol, 78%).

$^1$H NMR (400 MHz, $CDCl_3$) δ 8.24 (d, *J* = 2.1 Hz, 1H), 7.75 (s, 1H), 7.28 – 7.20 (m, 5H), 6.96 (dd, *J* = 8.4, 2.1 Hz, 1H), 6.82 (d, *J* = 8.4 Hz, 1H), 5.30 – 5.17 (m, 2H), 5.10 – 5.01 (m, 3H), 4.92 (d, *J* = 7.7 Hz, 1H), 4.58 (s, 2H), 4.25 (dd, *J* = 12.4, 5.4 Hz, 1H), 4.07 (dd, *J* = 12.3, 2.4 Hz, 1H), 3.80 (ddd, *J* = 10.1, 5.4, 2.4 Hz, 1H), 2.41 (ddd, *J* = 10.1, 5.8, 2.3 Hz, 4H), 2.04 – 1.89 (m, 14H), 0.84 (s, 9H), 0.00 (s, 6H) ppm.

$^{13}$C NMR (101 MHz, $CDCl_3$) δ 172.9, 170.7, 170.5, 170.4, 170.0, 169.4, 143.9, 137.2, 136.0, 128.6, 128.6, 128.5, 128.2, 121.1, 118.1, 113.6, 99.9, 72.1, 71.9, 71.4, 68.3, 66.2, 64.6, 61.7, 36.4, 33.5, 26.0, 20.9, 20.8, 20.6, 20.6, 20.6, 18.4, -5.2 ppm.

LC-MS (ESI): (m/z) = 788 [M+H]

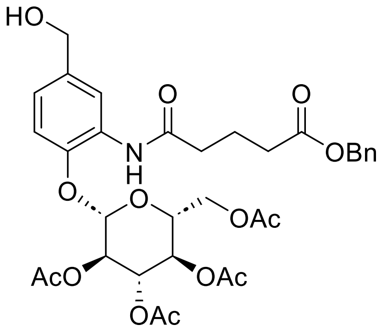


Exact Mass: 673.24

**(2R,3R,4S,5R,6S)-2-(acetoxymethyl)-6-(2-(5-(benzyloxy)-5-oxopentanamido)-4-(hydroxy methyl)phenoxy)tetrahydro-2H-pyran-3,4,5-triyl triacetate (S14):** Followed **S4** synthetic procedure on 0.9 g (1.14 mmol) scale of **S13**. Obtained title compound **S14** (736 mg, 1.09 mmol, 96%) as thick oil.

$^{1}$H NMR (400 MHz, $CDCl_3$) δ 8.41 (d, *J* = 2.1 Hz, 1H), 7.38 – 7.29 (m, 5H), 7.06 (dd, *J* = 8.4, 2.1 Hz, 1H), 6.94 (d, *J* = 8.4 Hz, 1H), 5.38 (t, *J* = 9.5 Hz, 1H), 5.30 (dd, *J* = 9.8, 7.7 Hz, 1H), 5.19 – 5.12 (m, 3H), 5.04 (d, *J* = 7.7 Hz, 1H), 4.63 (s, 2H), 4.34 (dd, *J* = 12.4, 5.4 Hz, 1H), 4.17 (dd, *J* = 12.4, 2.4 Hz, 1H), 3.92 (ddd, *J* = 10.1, 5.4, 2.4 Hz, 1H), 2.51 (t, *J* = 7.2 Hz, 4H), 2.18 – 2.02 (m, 16H) ppm.

$^{13}$C NMR (101 MHz, $CDCl_3$) δ 172.9, 170.9, 170.5, 170.5, 170.0, 169.5, 144.4, 136.7, 136.0, 128.7, 128.6, 128.2, 122.1, 119.1, 113.8, 99.7, 72.2, 71.9, 71.5, 68.2, 66.3, 64.9, 61.7, 36.3, 33.4, 20.9, 20.7, 20.7, 20.6, 20.6 ppm.

LC-MS (ESI): (m/z) = 674 [M+H]

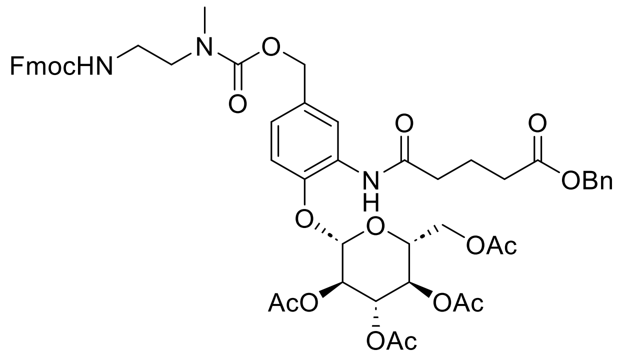


Exact Mass: 995.37

**(2S,3R,4S,5R,6R)-2-(4-(10-(9H-fluoren-9-yl)-4-methyl-3,8-dioxo-2,9-dioxa-4,7-diazadecyl)-2-(5-(benzyloxy)-5-oxopentanamido)phenoxy)-6-(acetoxymethyl)tetrahydro-2H-pyran-3,4,5-triyl triacetate (S15):** Followed **S6** synthetic procedure on 0.7 g (1.04 mmol) scale of **S14**. Obtained title compound **S15** (724 mg, 0.73 mmol, 70%) as a white solid.

$^{1}$H NMR (500 MHz, $CDCl_3$) δ 8.41 (d, *J* = 30.5 Hz, 1H), 7.87 – 7.68 (m, 1H), 7.68 (s, 1H), 7.49 (qd, *J* = 7.9, 5.6 Hz, 3H), 7.26 (ddt, *J* = 37.4, 20.7, 8.7 Hz, 12H), 6.88 – 6.73 (m, 2H), 5.31 – 5.14 (m, 2H), 5.10 – 4.92 (m, 6H), 4.84 (dd, *J* = 17.2, 7.9 Hz, 1H), 4.25 (d, *J* = 7.2 Hz, 2H), 4.19 (dd, *J* = 12.5, 5.4 Hz, 1H), 4.15 – 3.97 (m, 2H), 3.76 – 3.66 (m, 1H), 3.38 (t, *J* = 6.3 Hz, 2H), 3.31 (q, *J* = 6.2 Hz, 2H), 2.89 (d, *J* = 3.8 Hz, 3H), 2.39 – 2.26 (m, 3H), 2.02 – 1.90 (m, 14H) ppm.

$^{13}$C NMR (126 MHz, $CDCl_3$) δ 172.9, 172.8, 172.8, 171.3, 171.1, 170.5, 170.5, 170.4, 170.0, 169.4, 156.8, 156.7, 156.0, 144.3, 144.0, 141.4, 141.3, 136.0, 135.9, 132.4, 128.8, 128.6, 128.5, 128.3, 128.2, 128.2, 127.7, 127.7, 127.1, 125.2, 125.1, 122.6, 121.9, 120.0, 119.9, 119.4, 118.6, 113.4, 99.5, 72.1, 71.8, 71.5, 68.2, 66.7, 66.6, 66.3, 66.2, 61.7, 48.5, 47.3, 39.1, 36.4, 35.4, 33.5, 33.4, 33.3, 20.9, 20.7, 20.7, 20.6 ppm.

LC-MS (ESI): (m/z) = 996 [M+H]

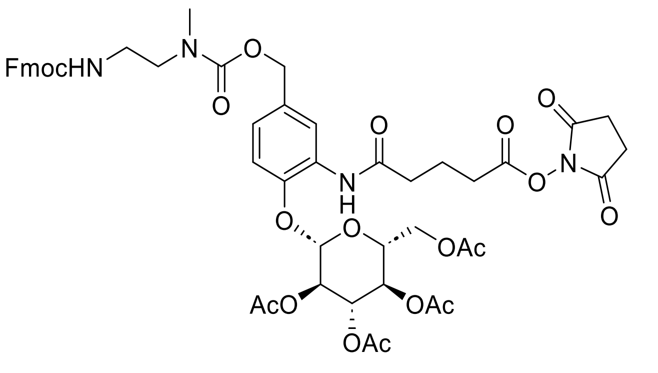


**(2S,3R,4S,5R,6R)-2-(4-(10-(9H-fluoren-9-yl)-4-methyl-3,8-dioxo-2,9-dioxa-4,7-diazadecyl)-2-(5-((2,5-dioxopyrrolidin-1-yl)oxy)-5-oxopentanamido)phenoxy)-6-(acetoxymethyl) tetrahydro-2H-pyran-3,4,5-triyl triacetate (S16):** Followed **S7** synthetic procedure on 0.5 g (0.74 mmol) scale of **S15**. Obtained title compound **S16** (600 mg, 0.56 mmol, 81%) white solid. $^{1}$H NMR (500 MHz, DMSO) δ 8.64 (s, 1H), 7.81 (d, *J* = 7.5 Hz, 3H), 7.61 (d, *J* = 6.6 Hz, 2H), 7.32 (dt, *J* = 11.9, 6.7 Hz, 2H), 7.25 (t, *J* = 7.4 Hz, 2H), 7.07 – 6.96 (m, 2H), 5.47 – 5.32 (m, 2H), 5.05 (t, *J* = 8.9 Hz, 1H), 4.97 – 4.85 (m, 3H), 4.23 (d, *J* = 6.8 Hz, 2H), 4.15 (dd, *J* = 19.7, 11.7 Hz, 4H), 4.00 (t, *J* = 10.1 Hz, 1H), 3.05 (q, *J* = 6.3 Hz, 2H), 2.76 (d, *J* = 9.7 Hz, 6H), 2.68 (s, 1H), 2.37 (s, 2H), 1.93 (dd, *J* = 12.6, 3.2 Hz, 12H).

LC-MS (ESI): (m/z) = 1003 [M+H]

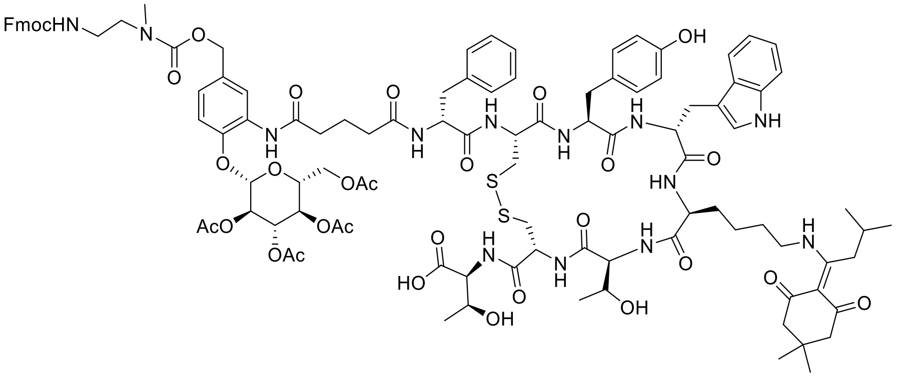


**((4R,7S,10S,13R,16S,19R)-13-((1H-indol-3-yl)methyl)-19-((R)-2-(5-((5-(10-(9H-fluoren-9-yl)-4-methyl-3,8-dioxo-2,9-dioxa-4,7-diazadecyl)-2-(((2S,3R,4S,5R,6R)-3,4,5-triacetoxy-6-(acetoxymethyl)tetrahydro-2H-pyran-2-yl)oxy)phenyl)amino)-5-oxopentanamido)-3-phenylpropanamido)-10-(4-((1-(4,4-dimethyl-2,6-dioxocyclohexylidene)-3-methylbutyl) amino)butyl)-16-(4-hydroxybenzyl)-7-((R)-1-hydroxyethyl)-6,9,12,15,18-pentaoxo-1,2-dithia-5,8,11,14,17-pentaazacycloicosane-4-carbonyl)-L-allothreonine (S17):** Followed **S9** synthetic procedure on 100 mg (0.099 mol) scale of **S16**. Obtained title compound **S17** (132 mg, 0.062 mmol, 62%) white solid.

LC-MS (ESI): (m/z) = 2143 [M+H]

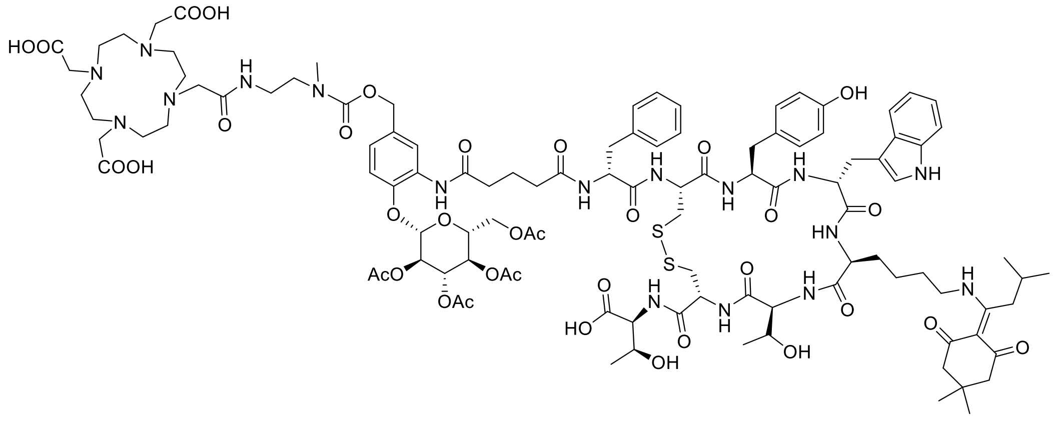


Exact Mass: 2305.97

**2,2',2''-(10-(2-((2-((((3-(5-(((R)-1-(((4R,7S,10S,13R,16S,19R)-13-((1H-indol-3-yl)methyl)-4-(((1S,2S)-1-carboxy-2-hydroxypropyl)carbamoyl)-10-(4-((1-(4,4-dimethyl-2,6-dioxocyclo hexylidene)-3-methylbutyl)amino)butyl)-16-(4-hydroxybenzyl)-7-((R)-1-hydroxyethyl)-6,9,12,15,18-pentaoxo-1,2-dithia-5,8,11,14,17-pentaazacycloicosan-19-yl)amino)-1-oxo-3-phenylpropan-2-yl)amino)-5-oxopentanamido)-4-(((2S,3R,4S,5R,6R)-3,4,5-triacetoxy-6-(acetoxymethyl)tetrahydro-2H-pyran-2-yl)oxy)benzyl)oxy)carbonyl)(methyl)amino)ethyl) amino)-2-oxoethyl)-1,4,7,10-tetraazacyclododecane-1,4,7-triyl)triacetic acid (S18):**

Followed **S10** synthetic procedure on 50 mg (0.0233 mmol) scale of **S17**. Obtained title compound **S18** (30 mg, 0.0131 mmol, 56%) as a white solid.

LC-MS (ESI): (m/z) = 2307 [M+H]

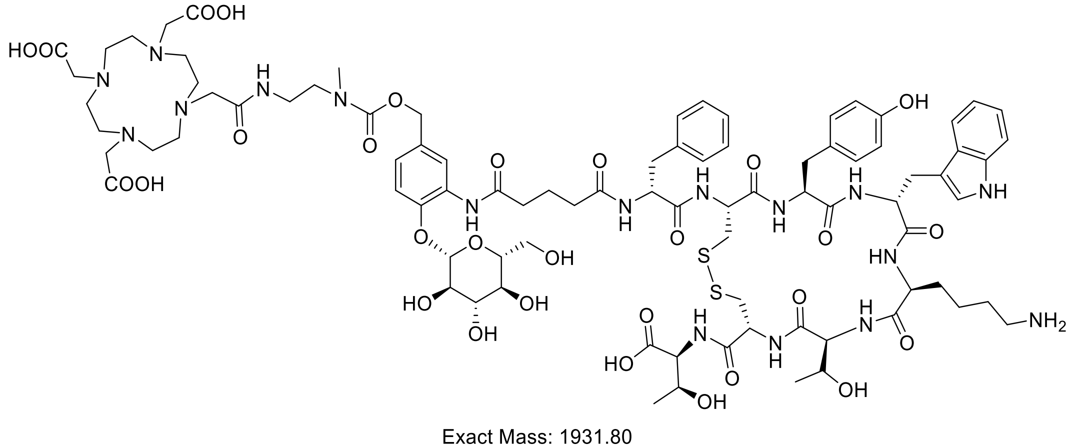


Exact Mass: 1931.80

**DOTA-β-Glu-TATE:** Followed **S11** synthetic procedure on 20 mg (0.0087 mmol) scale of **S18**. Obtained title compound **S19** (6.7 mg, 0.0035 mmol, 40%) white solid.

LC-MS (ESI): (m/z) = 1932 [M+H]

**DOTA-β-GlcA-TATE Synthesis:**

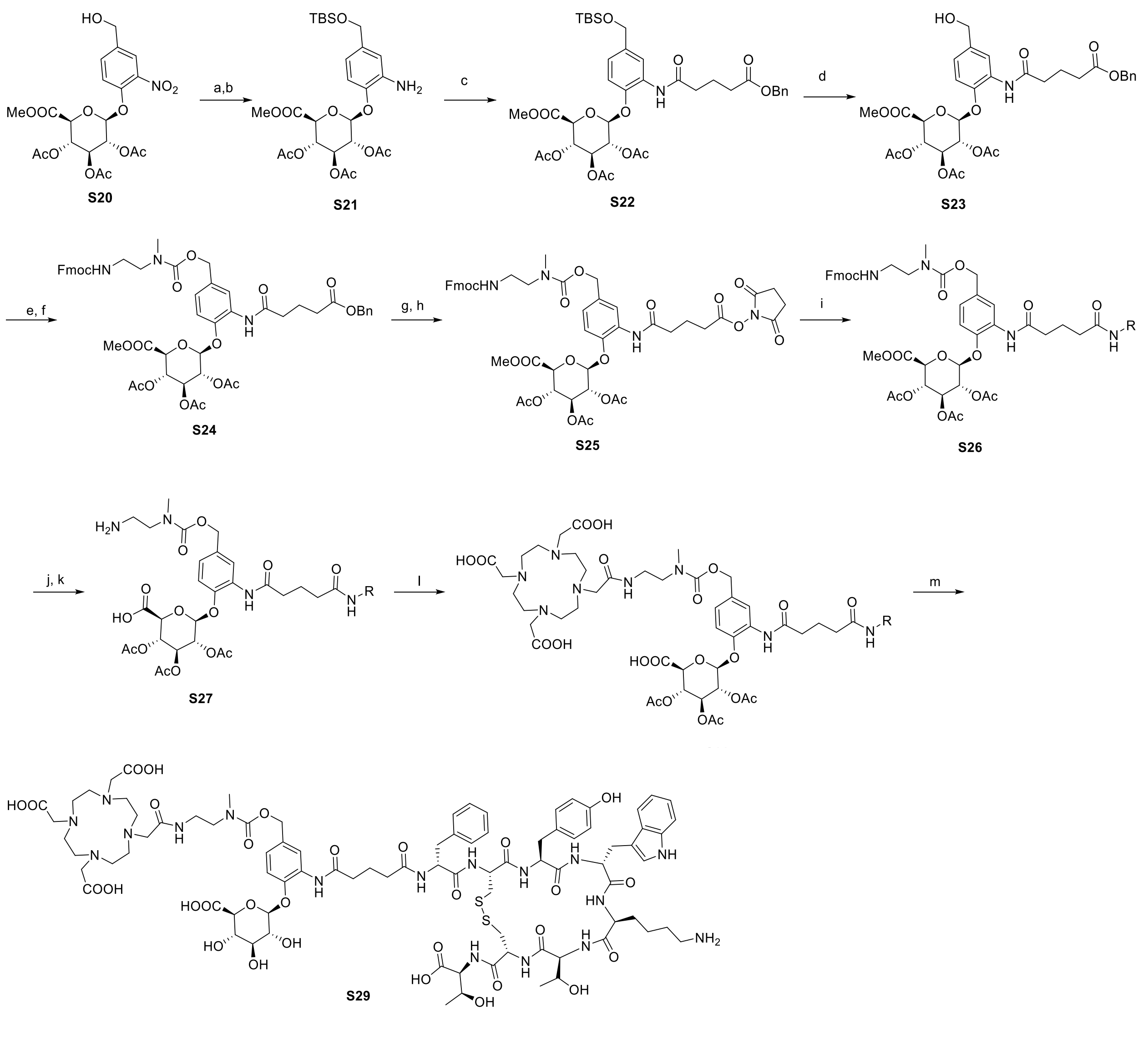

**Scheme S3.** *Reagents and conditions*: a) TBSCl, imidazole, DCM, RT, overnight, b) Zn powder, $NH_4Cl$, MeOH, 2 h, 0 °C - RT 80%; c) 5-(benzyloxy)-5-oxopentanoic acid, HATU, DIPEA, DMF, RT, overnight, 71%; d) $TsOH.H_2O$, MeOH, RT, 30 min, 90%; e) 4-nitrophenyl chloroformate, pyridine, DCM, 0 °C -RT, 2 h, f) **S5**, DIPEA, DMF, RT, 30 min, 67%; g) Pd/C (10 wt. %), MeOH, RT, 1 h; h) TSTU, DIPEA, DMF, RT, 1 h, 74%; i) **S8**, acetonitrile and 0.1M phosphate buffer (pH = 7.3), RT, 20 h, 47%; j) $Me_3SnOH$, DCE, Mw 100 °C, 2 h, k) piperidine, DMF, RT; 30 min, 60%; l) DOTA-NHS-ester**,** DIPEA, DMF, RT, 2 h, 50%; m) Hydrazine hydrate, DMF, RT, 30 min, 45%.

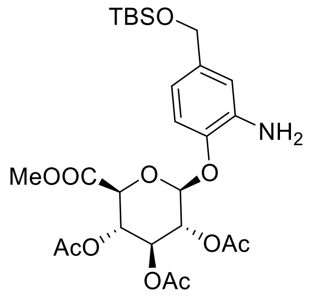


**(2S,3R,4S,5S,6S)-2-(2-amino-4-(((tert-butyldimethylsilyl)oxy)methyl)phenoxy)-6-(methoxy carbonyl)tetrahydro-2H-pyran-3,4,5-triyl triacetate (S21):** The (2S,3R,4S,5S,6S)-2-(4-(hydroxymethyl)-2-nitrophenoxy)-6-(methoxycarbonyl)tetrahydro-2H-pyran-3,4,5-triyl triacetate **S20** (1.0 g, 2.06 mmol) was dissolved in DCM (20 mL), was added sequentially TBSCl (452 mg, 3.09 mmol) and imidazole (204 mg, 3.09 mmol) at 0 °C. The resulting reaction stirred overnight. After completion, added water to the reaction mixture and extracted with DCM (2 x 100 mL). The combined organic layers were washed with brine solution (20 mL) and concentrated to give crude product. The crude product was purified by flash chromatography on silica gel eluting with ethyl acetate in hexane (0 to 30% gradient) which was used for next step without further characterization. The resulting TBS protected compound (2.06 mmol) was dissolved in methanol (40 mL), was added sequentially zinc (1.33 g, 20.6 mmol) and ammonium chloride (1.09 g, 20.6 mmol) at 0 °C. The reaction was stirred on ice for 15 min then the ice bath was removed, and stirring was continued at RT for 2 hours. The reaction was filtered through celite with methanol, and the filtrate was concentrated in vacuo. The crude residue was re-suspended in DCM (100 mL) and saturated $NaHCO_3$ solution (50 mL). The layers were separated, and the aqueous layers were extracted with DCM (2 x 100 mL). The combined organic layers dried over sodium sulfate and concentrated in vacuo to give crude product. The crude product was purified by flash chromatography on silica gel eluting with ethyl acetate in hexane (0 to 50% gradient) to give the title compound as a white solid (0.93 g, 1.6 mmol, 80%).

$^1$H NMR (400 MHz, $CDCl_3$) δ 8.29 (d, $J$ = 2.0 Hz, 1H), 7.96 (s, 1H), 7.34 – 7.28 (m, 5H), 7.02 (dd, $J$ = 8.4, 2.1 Hz, 1H), 6.91 (d, $J$ = 8.3 Hz, 1H), 5.44 – 5.34 (m, 1H), 5.33 – 5.22 (m, 2H), 5.11 (s, 2H), 5.05 (d, $J$ = 7.6 Hz, 1H), 4.56 (s, 2H), 4.20 (d, $J$ = 9.7 Hz, 1H), 3.70 (s, 3H), 2.48 (td, $J$ = 7.4, 1.9 Hz, 4H), 2.38 (s, 2H), 2.10 – 1.98 (m, 11H) ppm.

$^{13}$C NMR (101 MHz, $CDCl_3$) δ 172.9, 171.1, 170.2, 169.9, 169.4, 166.6, 144.5, 137.3, 136.0, 129.0, 128.5, 128.2, 122.4, 119.5, 115.0, 100.1, 72.3, 71.1, 69.2, 66.2, 64.7, 53.1, 36.2, 33.4, 20.7, 20.7, 20.6, 20.4 ppm.

LC-MS (ESI): (m/z) = 570 [M+H]

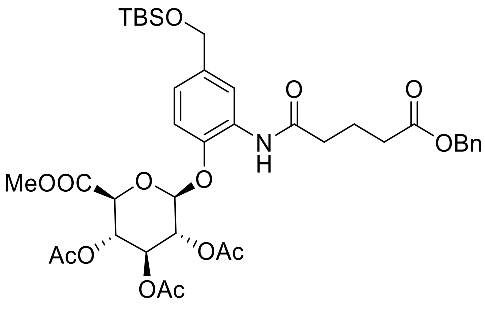


**(2S,3R,4S,5S,6S)-2-(2-(5-(benzyloxy)-5-oxopentanamido)-4-(((tert-butyldimethylsilyl)oxy) methyl)phenoxy)-6-(methoxycarbonyl)tetrahydro-2H-pyran-3,4,5-triyl triacetate (S22):** Compound **S21** (0.9 g, 1.58 mmol) in DMF (10 mL), was added sequentially 5-(benzyloxy)-5-oxopentanoic acid (421 mg, 1.89 mmol), HATU (0.9 g, 2.37 mmol), and DIPEA (0.64 mL 3.9 mmol) at 0 °C. The resulting mixture was stirred at RT overnight. After completion, water added to the reaction mixture at 0 °C and extracted with ethyl acetate (3 x 150 mL). The combined organics were washed with water (50 mL), brine solution (50 mL), and dried over anhydrous $Na_2SO_4$. The combined organic layers were concentrated in vacuo to give crude compound. The crude product was purified by flash chromatography on silica gel eluting with ethyl acetate in hexane (0 to 50% gradient) to give the title compound (867 mg, 1.12 mmol, 71%) as thick oil.

$^1$H NMR (500 MHz, $CDCl_3$) δ 8.22 (d, *J* = 2.2 Hz, 1H), 7.87 (s, 1H), 7.30 – 7.17 (m, 5H), 6.96 (dd, *J* = 8.5, 2.1 Hz, 1H), 6.84 (d, *J* = 8.4 Hz, 1H), 5.27 – 5.17 (m, 2H), 5.04 (s, 2H), 4.98 (d, *J* = 7.7 Hz, 1H), 4.58 (s, 2H), 4.13 (d, *J* = 9.7 Hz, 1H), 3.63 (s, 3H), 2.41 (t, *J* = 7.4 Hz, 4H), 0.84 (s, 9H), -0.00 (d, *J* = 1.5 Hz, 6H) ppm.

$^{13}$C NMR (126 MHz, $CDCl_3$) δ 172.9, 170.8, 170.1, 169.8, 169.4, 166.6, 144.1, 137.6, 136.1, 129.0, 128.5, 128.1, 121.2, 118.3, 114.8, 100.3, 72.4, 71.2, 69.2, 66.2, 64.6, 60.3, 53.0, 36.3, 33.5, 26.1, 26.0, 25.8, 21.0, 20.8, 20.7, 20.6, 20.4, 18.4, 14.2, -5.0, -5.2, -5.5 ppm.

LC-MS (ESI): (m/z) = 774 [M+H]

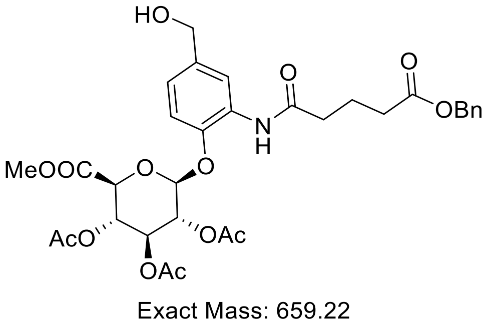


**(2R,3S,4R,5R,6R)-6-(2-(5-(benzyloxy)-5-oxopentanamido)-4-(hydroxymethyl)phenoxy) tetrahydro-2H-pyran-2,3,4,5-tetrayl tetraacetate (S23):** Followed **S4** synthetic procedure on 800 mg (1.0 mmol) scale of **S22**. Obtained title compound **S23** (613 mg, 0.93 mmol, 90%) white solid.

$^1$H NMR (400 MHz, $CDCl_3$) δ 8.29 (d, *J* = 2.0 Hz, 1H), 7.96 (s, 1H), 7.34 – 7.25 (m, 5H), 7.02 (dd, *J* = 8.4, 2.1 Hz, 1H), 6.91 (d, *J* = 8.3 Hz, 1H), 5.44 – 5.34 (m, 1H), 5.33 – 5.22 (m, 2H), 5.11 (s, 2H), 5.04 (s, 1H), 4.56 (s, 2H), 4.20 (d, *J* = 9.7 Hz, 1H), 3.70 (s, 3H), 2.48 (td, *J* = 7.4, 1.9 Hz, 4H), 2.10 – 1.98 (m, 11H) ppm.

$^{13}$C NMR (101 MHz, $CDCl_3$) δ 172.9, 171.1, 170.2, 169.9, 169.4, 166.6, 144.5, 137.3, 136.0, 129.0, 128.5, 128.2, 122.4, 119.5, 115.0, 100.1, 72.3, 71.1, 69.2, 66.2, 64.7, 53.1, 36.2, 33.4, 20.7, 20.7, 20.6, 20.4 ppm.

LC-MS (ESI): (m/z) = 660 [M+H]

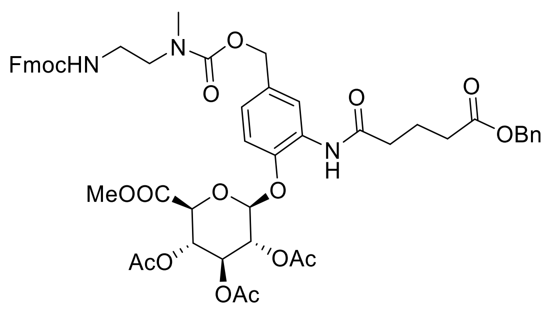


Exact Mass: 981.35

**(2S,3R,4S,5S,6S)-2-(4-(10-(9H-fluoren-9-yl)-4-methyl-3,8-dioxo-2,9-dioxa-4,7-diazadecyl)-2-(5-(benzyloxy)-5-oxopentanamido)phenoxy)-6-(methoxycarbonyl)tetrahydro-2H-pyran-3,4,5-triyl triacetate (S24):** Followed **S6** synthetic procedure on 0.5 g (0.75 mmol) scale of **S23**. Obtained title compound **S24** (498 mg, 0.5 mmol, 67%) white solid.

$^1$H NMR (500 MHz, $CDCl_3$) δ 8.39 (d, *J* = 32.3 Hz, 1H), 7.91 (d, *J* = 33.5 Hz, 1H), 7.70 – 7.60 (m, 3H), 7.49 (t, *J* = 8.0 Hz, 2H), 7.35 – 7.10 (m, 8H), 6.83 (ddd, *J* = 42.2, 16.7, 8.5 Hz, 2H), 5.99 (s, 1H), 5.30 (t, *J* = 9.6 Hz, 1H), 5.20 (tt, *J* = 9.6, 4.5 Hz, 2H), 5.08 – 4.82 (m, 5H), 4.25 (d, *J* = 7.0 Hz, 2H), 4.11 (q, *J* = 5.8 Hz, 1H), 4.00 (t, *J* = 10.8 Hz, 1H), 3.62 (s, 3H), 3.44 – 3.35 (m, 2H), 3.31 (q, *J* = 6.2 Hz, 2H), 2.89 (d, *J* = 3.9 Hz, 3H), 2.46 – 2.31 (m, 5H), 1.98 (q, *J* = 8.5 Hz, 12H) ppm.

$^{13}$C NMR (126 MHz, $CDCl_3$) δ 172.9, 172.8, 171.4, 171.1, 170.2, 169.8, 169.4, 166.5, 156.9, 156.7, 156.0, 144.5, 144.3, 144.1, 144.1, 143.4, 141.3, 140.2, 138.0, 136.0, 136.0, 133.0, 129.3, 128.8, 128.6, 128.5, 128.2, 127.7, 127.7, 127.1, 127.0, 125.2, 125.1, 122.7, 122.0, 121.0, 120.0, 119.8, 119.5, 118.8, 114.7, 114.6, 107.8, 100.0, 72.4, 71.1, 71.0, 69.2, 69.2, 66.7, 66.6, 66.3, 66.2, 53.1, 53.1, 48.5, 47.3, 39.1, 36.4, 35.4, 35.2, 33.5, 33.5, 33.4, 20.8, 20.6, 20.5 ppm.

LC-MS (ESI): (m/z) = 982 [M+H]

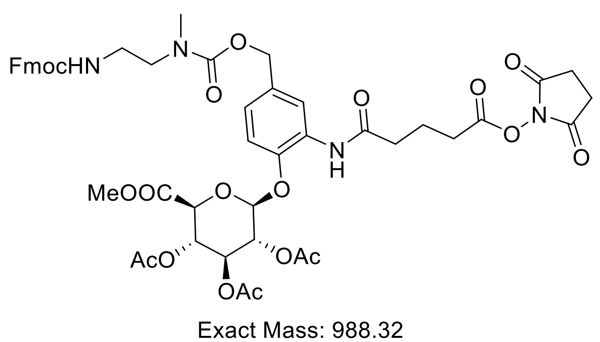


Exact Mass: 988.32

**(2S,3R,4S,5S,6S)-2-(4-(10-(9H-fluoren-9-yl)-4-methyl-3,8-dioxo-2,9-dioxa-4,7-diazadecyl)-2-(5-((2,5-dioxopyrrolidin-1-yl)oxy)-5-oxopentanamido)phenoxy)-6-(methoxycarbonyl)tetrahydro-2H-pyran-3,4,5-triyl triacetate (S25):** Followed **S7** synthetic procedure on 300 mg (0.3 mmol) scale of **S24**. Obtained title compound **S25** (223 mg, 0.22 mmol, 74%) white solid.

$^1$H NMR (500 MHz, DMSO): δ 8.78 (s, 1H), 7.89 (d, *J* = 7.6 Hz, 2H), 7.67 (t, *J* = 6.5 Hz, 2H), 7.40 (dt, *J* = 12.9, 6.7 Hz, 3H), 7.32 (t, *J* = 7.4 Hz, 2H), 7.09 (dt, *J* = 23.2, 8.1 Hz, 2H), 5.59 (d, *J* = 7.7 Hz, 1H), 5.49 (t, *J* = 9.6 Hz, 1H), 5.19 (t, *J* = 8.7 Hz, 1H), 5.07 (t, *J* = 9.7 Hz, 1H), 4.95 (d, *J* = 9.1 Hz, 2H), 4.72 (d, *J* = 9.8 Hz, 1H), 4.31 (d, *J* = 6.8 Hz, 2H), 4.20 (d, *J* = 7.6 Hz, 1H), 3.63 (d, *J* = 3.6 Hz, 3H), 3.13 (q, *J* = 6.3 Hz, 2H), 2.83 (d, *J* = 9.4 Hz, 7H), 2.76 (d, *J* = 8.7 Hz, 2H), 2.46 (s, 2H), 2.02 (d, *J* = 2.7 Hz, 9H), 1.94 (d, *J* = 7.4 Hz, 2H) ppm.

$^{13}$C NMR (126 MHz, DMSO): δ 174.6, 170.8, 170.7, 170.1, 170.0, 169.8, 169.2, 167.5, 156.6, 155.9, 147.2, 144.4, 141.2, 132.1, 128.4, 128.1, 127.5, 125.6, 125.6, 123.1, 120.6, 115.3, 98.2, 71.4, 71.2, 71.1, 69.4, 66.2, 65.7, 53.1, 48.7, 48.2, 47.2, 35.4, 34.8, 30.1, 25.9, 20.9, 20.8, 20.7, 20.5 ppm.

LC-MS (ESI): (m/z) = 989 [M+H]

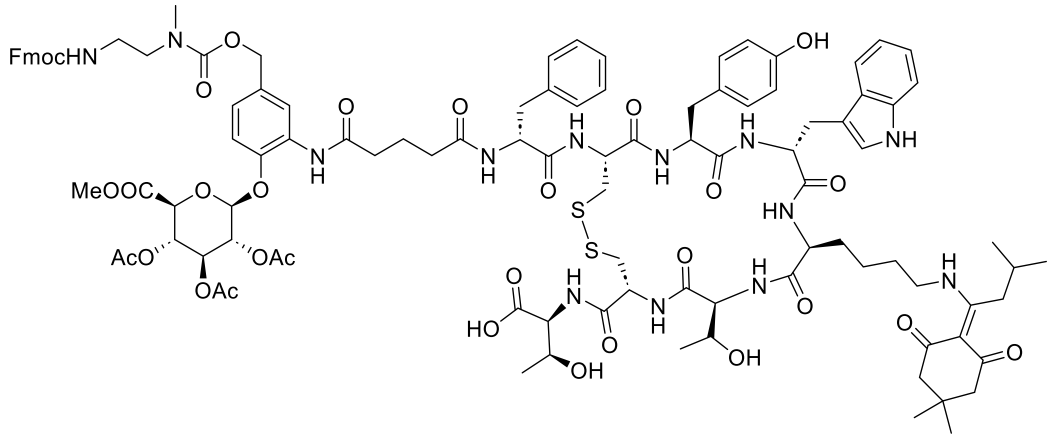


Exact Mass: 2127.84

**((4R,7S,10S,13R,16S,19R)-13-((1H-indol-3-yl)methyl)-19-((R)-2-(5-((5-(10-(9H-fluoren-9-yl)-4-methyl-3,8-dioxo-2,9-dioxa-4,7-diazadecyl)-2-(((2S,3R,4S,5S,6S)-3,4,5-triacetoxy-6-(methoxycarbonyl)tetrahydro-2H-pyran-2-yl)oxy)phenyl)amino)-5-oxopentanamido)-3-phenylpropanamido)-10-(4-((1-(4,4-dimethyl-2,6-dioxocyclohexylidene)-3-methylbutyl) amino)butyl)-16-(4-hydroxybenzyl)-7-((R)-1-hydroxyethyl)-6,9,12,15,18-pentaoxo-1,2-dithia-5,8,11,14,17-pentaazacycloicosane-4-carbonyl)-L-allothreonine (S26):**

Followed **S9** synthetic procedure on 100 mg (0.101 mmol) scale of **S25**. Obtained title compound **S26** (101 mg, 0.047 mmol, 47%) white solid.

LC-MS (ESI): (m/z) = 2129 [M+H]

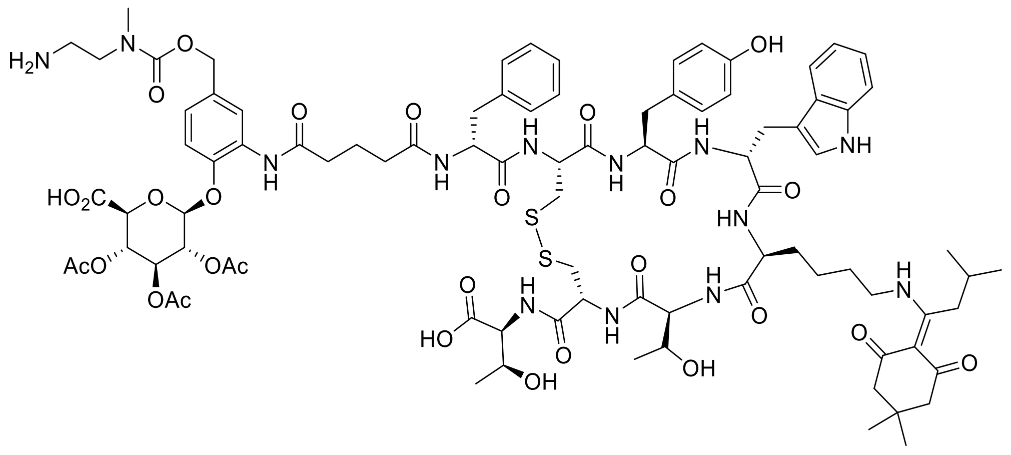


**((4R,7S,10S,13R,16S,19R)-13-((1H-indol-3-yl)methyl)-19-((R)-2-(5-((5-((((2-aminoethyl)(methyl)carbamoyl)oxy)methyl)-2-(((2S,3R,4S,5S,6S)-3,4,5-triacetoxy-6-carboperoxy tetrahydro-2H-pyran-2-yl)oxy)phenyl)amino)-5-oxopentanamido)-3-phenylpropanamido)-10-(4-((1-(4,4-dimethyl-2,6-dioxocyclohexylidene)-3-methylbutyl)amino)butyl)-16-(4-hydroxybenzyl)-7-((R)-1-hydroxyethyl)-6,9,12,15,18-pentaoxo-1,2-dithia-5,8,11,14,17-penta azacycloicosane-4-carbonyl)-L-allothreonine (S27):** The carboxylic ester **S26** (80 mg, 0.0376 mmol) was dissolved in 1,2-dichloroethane, trimethyltinhydroxide (41 mg, 0.225 mmol). The mixture was subjected to microwave irradiation at 100 °C for 2 h. After completion of the reaction as indicated by LCMS, the mixture was concentrated in vacuo, and the residue was purified by reverse phase chromatography without workup using a gradient of 10-70% acetonitrile in water with the addition of 0.05% TFA to get the primary amine, this compound was used for the next step without further analysis. The resulting compound (0.0376, mmol) was dissolved in 20% piperidine in DMF (1 mL), and the solution was stirred for 30 min at RT. After completion of the reaction (as indicated by LCMS) crude material was purified by reverse phase chromatography without workup using a gradient of 10-50% acetonitrile in water with the addition of 0.05% TFA to get title compound (43 mg, 0.023 mmol, 60%) white solid.

LC-MS (ESI): (m/z) = 1893 [M+H]

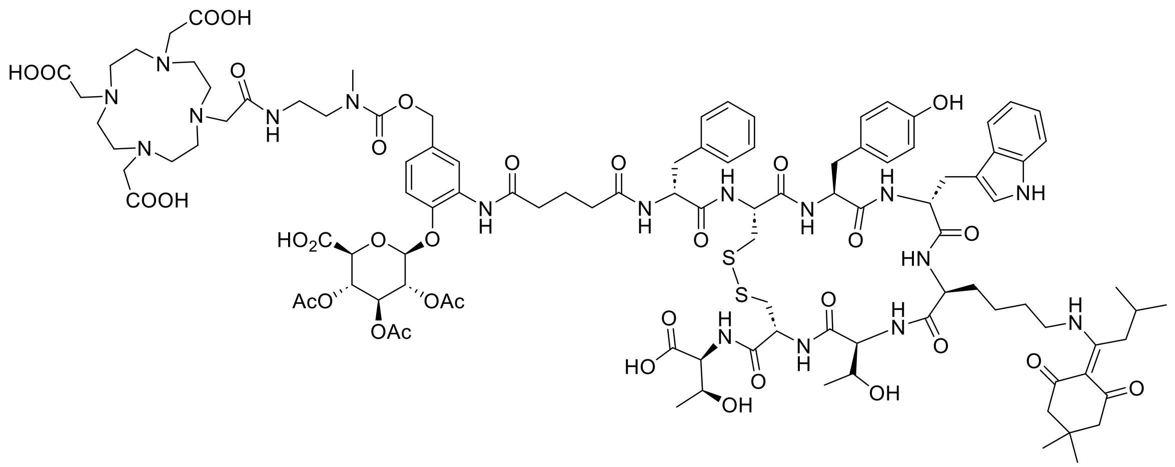

**2,2',2''-(10-(2-((2-((((3-(5-(((R)-1-(((4R,7S,10S,13R,16S,19R)-13-((1H-indol-3-yl)methyl)-4-(((1S,2S)-1-carboxy-2-hydroxypropyl)carbamoyl)-10-(4-((1-(4,4-dimethyl-2,6-dioxocyclo hexylidene)-3-methylbutyl)amino)butyl)-16-(4-hydroxybenzyl)-7-((R)-1-hydroxyethyl)-6,9,12,15,18-pentaoxo-1,2-dithia-5,8,11,14,17-pentaazacycloicosan-19-yl)amino)-1-oxo-3-phenylpropan-2-yl)amino)-5-oxopentanamido)-4-(((2S,3R,4S,5S,6S)-3,4,5-triacetoxy-6-carboxytetrahydro-2H-pyran-2-yl)oxy)benzyl)oxy)carbonyl)(methyl)amino)ethyl)amino)-2-oxoethyl)-1,4,7,10-tetraazacyclododecane-1,4,7-triyl)triacetic acid (S28).** Followed **S10** synthetic procedure on 30 mg (0.0158 mmol) **S27**. Obtained title compound **S28** (18 mg, 0.0079 mmol, 50%) white solid.

LC-MS (ESI): (m/z) = 2279 [M+H]

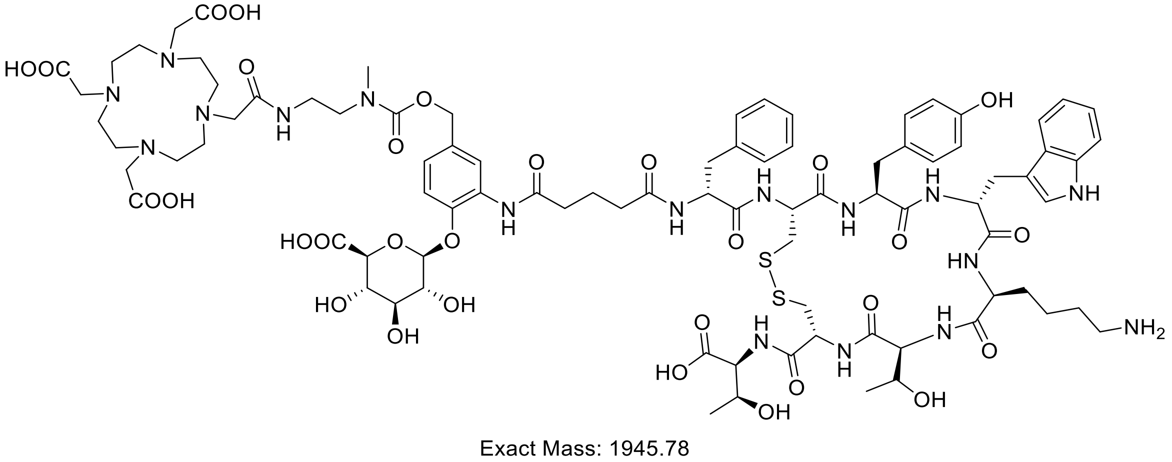


**DOTA-β-GlcA-TATE (S29):** Followed **S11** synthetic procedure on 17 mg (0.0075 mmol) scale of **S28**. Obtained title compound **S29** (6.5 mg, 0.0034 mmol, 45%) white solid.

LC-MS (ESI): (m/z) = 1947 [M+H]

**DOTA-G-Chol-G-TATE Synthesis:**

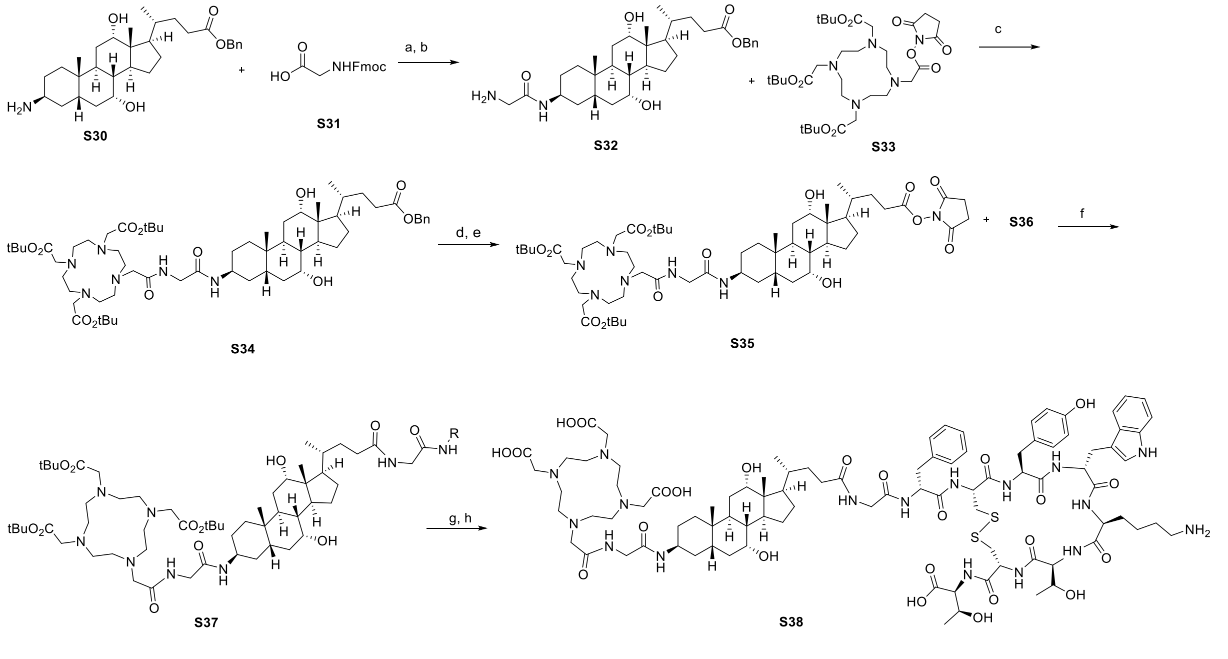


**Scheme S4.** *Reagents and conditions*: a) HATU, DIPEA, DMF, RT, 3 h; b) pieridine, DCM, RT 30 min, 65% over two steps; c) DIPEA, DMF, RT, 24 h, 80%; d) Pd/C (10 wt. %), $H_2$, MeOH, RT, 1 h; e) TSTU, DIPEA, DMF, RT, 1 h, 68% over two steps; f) DIPEA, DMF, RT, 2 h, 46%; g) TFA:TIPS:$H_2O$ (98:1:1), 1h; h); 5% Hydrazine hydrate, DMF, RT, 30 min, over two steps 38%.

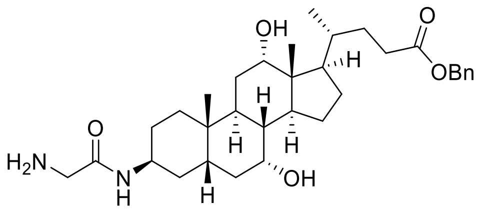


Exact Mass: 554.3720

**Benzyl (*R*)-4-((3*S*,5*S*,7*R*,8*R*,9*S*,10*S*,12*S*,13*R*,14*S*,17*R*)-3-(2-aminoacetamido)-7,12-dihydroxy-10,13-dimethylhexadecahydro-1H-cyclopenta[a]phenanthren-17-yl)pentanoate (S32)**: To a solution of compound **S30***(61)* (0.5 g, 1.0 mmol) in DMF (5 mL), was added sequentially **S31** (358 mg, 1.2 mmol), HATU (494 mg, 1.3 mol), and DIPEA (0.43 mL, 2.5 mmol) at RT. The resulting solution was stirred at RT for 3 h. After completion of the reaction (as indicated by LCMS), the reaction mixture was cooled 0 °C and added water (10 mL), the product was precipitated. The product was filtered up and dried to give crude product, which was used for the next step without further purification.

The above crude product was (1.0 mmol) was dissolved in 20% piperidine in DCM (5 mL), and the solution was stirred for 30 min at RT. After completion of the reaction (as indicated by LCMS), the volatiles were removed and the crude product was purified by flash chromatography on silica gel eluting with methanol in DCM (0 to 10% gradient) to give the title compound (387 mg, 0.7 mmol, 65% over two steps) as a thick oil.

$^1$H NMR (500 MHz, $CDCl_3$) δ 7.75 (s, 1H), 7.39 – 7.27 (m, 5H), 5.16 – 5.01 (m, 2H), 4.08 (s, 1H), 3.89 (d, *J* = 56.5 Hz, 2H), 3.58 – 3.38 (m, 2H), 3.15 (t, *J* = 5.6 Hz, 4H), 2.54 (t, *J* = 13.6 Hz, 1H), 2.41 (ddd, *J* = 14.5, 9.4, 4.5 Hz, 1H), 2.29 (dt, *J* = 15.0, 7.6 Hz, 1H), 2.22 – 2.17 (m, 1H), 2.04 – 1.03 (m, 22H), 1.01 – 0.85 (m, 6H), 0.65 (s, 3H) ppm.

$^{13}$C NMR (126 MHz, $CDCl_3$) δ 174.1, 136.1, 128.5, 128.2, 128.2, 73.0, 68.3, 66.1, 53.5, 47.2, 46.5, 45.3, 44.7, 41.9, 39.4, 37.2, 35.2, 34.3, 33.6, 31.4, 31.1, 30.9, 28.5, 27.5, 26.0, 24.7, 23.2, 23.0, 22.6, 22.4, 17.3, 12.5 ppm.

LC-MS (ESI): (m/z) = 555 [M+H]

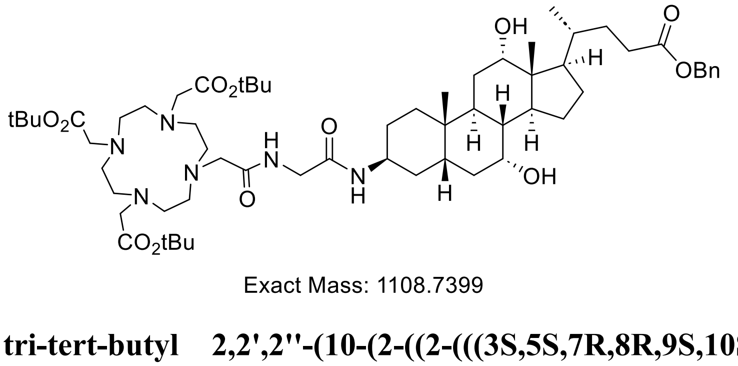


Exact Mass: 1108.7399

**tri-tert-butyl 2,2',2''-(10-(2-((2-(((3S,5S,7R,8R,9S,10S,12S,13R,14S,17R)-17-((R)-5-(benzyl oxy)-5-oxopentan-2-yl)-7,12-dihydroxy-10,13-dimethylhexadecahydro-1H-cyclopenta[a] phenanthren-3-yl)amino)-2-oxoethyl)amino)-2-oxoethyl)-1,4,7,10-tetraazacyclododecane-1,4,7-triyl)triacetate (S34):**To a solution of compound **S32** (0.35 g, 0.63 mmol) in DMF (5 mL), was added sequentially **S33** (405 mg, 0.75 mmol), and DIPEA (0.274 mL, 1.57 mmol) at 0 °C temperature. The resulting solution was stirred at RT for 2 h. After completion of the reaction (as indicated by LCMS), the crude material was purified by reverse phase chromatography without workup using a gradient of 10-60% acetonitrile in water with the addition of 0.05% TFA to get the corresponding product (560 mg, 0.05 mmol, 80% over two steps).

LC-MS (ESI): (m/z) = 1109 [M+H]

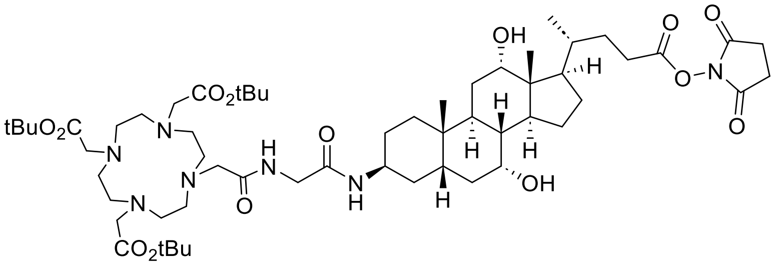


Exact Mass: 1115.7094

**tri-tert-butyl 2,2',2''-(10-(2-((2-(((3S,5S,7R,8R,9S,10S,12S,13R,14S,17R)-17-((R)-5-((2,5-dioxopyrrolidin-1-yl)oxy)-5-oxopentan-2-yl)-7,12-dihydroxy-10,13-dimethylhexadecahydro -1H-cyclopenta[a]phenanthren-3-yl)amino)-2-oxoethyl)amino)-2-oxoethyl)-1,4,7,10-tetraazacyclododecane-1,4,7-triyl)triacetate (S35):** Followed **S7** synthetic procedure on 300 mg (0.27 mmol) scale of **S34**. Obtained title compound **S35** (205 mg, 0.184 mmol, 68% over two steps) white solid.

LC-MS (ESI): (m/z) = 1116 [M+H]

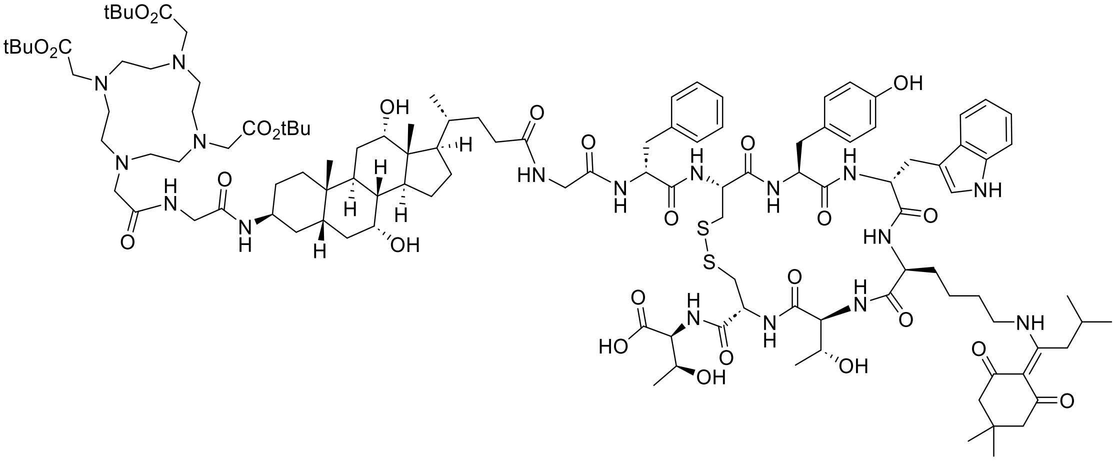


Exact Mass: 2312.2492

**((4R,7S,10S,13R,16S,19R)-13-((1H-indol-3-yl)methyl)-19-((R)-2-(2-((R)-4-((3S,5S,7R,8R,9S, 10S,12S,13R,14S,17R)-7,12-dihydroxy-10,13-dimethyl-3-(2-(2-(4,7,10-tris(2-(tert-butoxy)-2-oxoethyl)-1,4,7,10-tetraazacyclododecan-1-yl)acetamido)acetamido)hexadecahydro-1H-cyclopenta[a]phenanthren-17-yl)pentanamido)acetamido)-3-phenylpropanamido)-10-(4-((1-(4,4-dimethyl-2,6-dioxocyclohexylidene)-3-methylbutyl)amino)butyl)-16-(4-hydroxy**

**benzyl)-7-((R)-1-hydroxyethyl)-6,9,12,15,18-pentaoxo-1,2-dithia-5,8,11,14,17-pentaaza cycloicosane-4-carbonyl)-L-allothreonine (S37):** A solution of compound **S35** (84 mg, 0.076 mmol) in DMF (1 mL), was added sequentially **S36** (100 mg, 0.076 mmol), and DIPEA (0.026 mL, 0.15 mmol) at 0 °C temperature. The resulting solution was stirred at RT for 2 h. After completion of the reaction (as indicated by LCMS), the crude material was purified by reverse phase chromatography without workup using a gradient of 10-50% acetonitrile in water with the addition of 0.05% TFA to get the corresponding product (84 mg, 0.035 mmol, 46%).

LC-MS (ESI): (m/z) = 2313 [M+H]

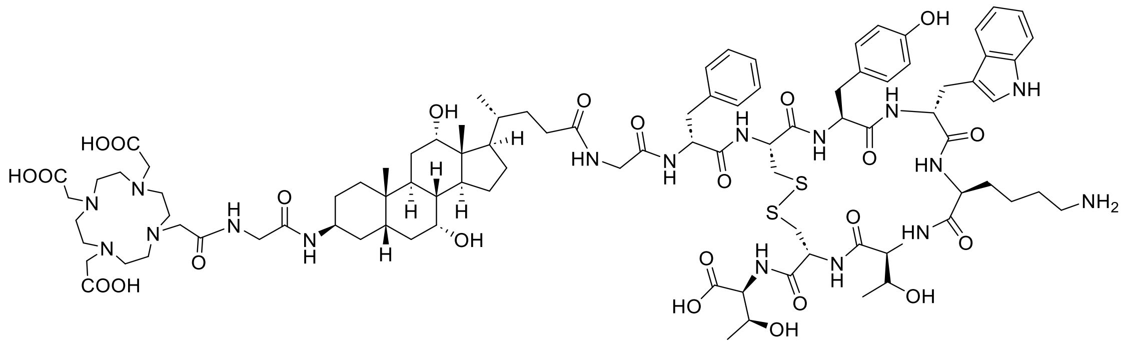


Exact Mass: 1937.9307

**DOTA-G-Chol-G-TATE (S38):**

A cleavage cocktail of 98:1:1 - TFA:TIPS:water was added to the compound **S37** (25 mg, 0.011 mmol) and stirred for 1 h. After the completion of the reaction the volatile were evaporated under reduced pressure. The residue was triturated with anhydrous ether (20.0 ml) and decanted. This process was repeated two more times and the fully deprotected peptide residues were used next ivDde deprotection. The crude compound (0.011 mmol) was dissolved in 5% hydrazine hydrate in DMF (1 mL) and the resulting solution was stirred at RT for 30 min. After completion of the reaction (as indicated by LCMS) crude solution was purified directly by reverse phase HPLC (HPLC conditions: Column: X-Terra® (Waters corp.) C18 RP; 150 x 30 mm; 5.0 microns; solvent A: water with 0.1% TFA (v/v) and solvent B: acetonitrile with 0.1% TFA (v/v); Elution rate: 50.0 ml/min; Gradient: 10%B – 65%B over 35 min; Detection @ 220 nm. Fractions with the required mass and purity of >95% were pooled and freeze-dried to yield the peptides as colorless fluffy solids) get the corresponding product (7.9 mg, 0.0041 mmol, 38% over two steps).

LC-MS (ESI): (m/z) = 1938 [M+H]

**DOTA-NH-NHMe Synthesis:**

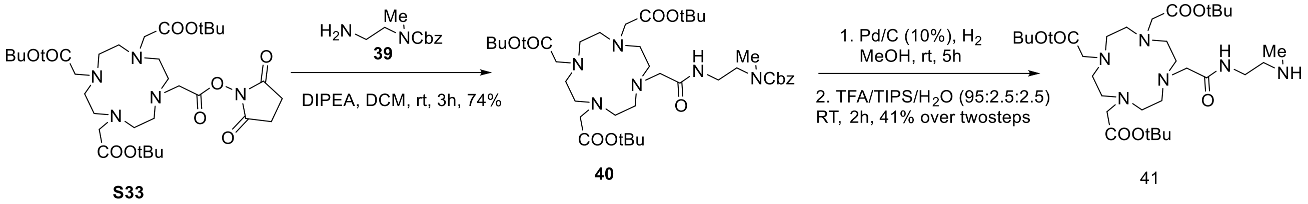


**Scheme S5.** *Reagents and conditions*: a) DIPEA, DCM, RT, 3 h, 74%; b) Pd/C (10 wt.%), $H_2$, MeOH, RT, 5 h; c) TFA:TIPS:$H_2O$ (98:1:1), RT, 2 h; 41% over two steps.

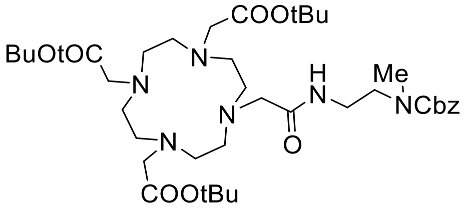


**tri-tert-butyl 2,2',2''-(10-(2-((2-(((benzyloxy)carbonyl)(methyl)amino)ethyl)amino)-2-oxoethyl)-1,4,7,10-tetraazacyclododecane-1,4,7-triyl)triacetate (S40):** To a stirred solution of S33 (1.0 g, 1.5 mol) and Ncbz N-methyl ethelene diamine (400 mg, 1.62 mol) in DCM (20 mL) and added and DIPEA (0.716 ml, 4.47 mmol) at 0 °C. Reaction mixture was stirred at RT for 3 h. After completion of the reaction (as indicated by LCMS), the volatiles were removed, and the crude product was purified by flash chromatography on silica gel eluting with methanol in DCM (0 to 10% gradient) to give the title compound (845 mg, 74%) as a thick oil.

LC-MS (ESI): (m/z) = 763 [M+H]

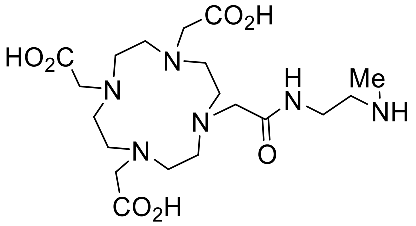


**DOTA-NH-NHMe (S41):** To a stirred solution of the compound **S40** (500.0 mg, 1 Eq, 0.65 mol) in MeOH (10 mL), was added Pd/C (75 mg, 10%w/w) at RT. The resulting solution was stirred at RT for 5 h under hydrogen pressure (hydrogen balloon was used for hydrogen pressure). After complete consumption of starting compound, the reaction mixture was filtered to remove the solids, and filtrate was concentrated to get the crude product which was pure enough and used for the next step without purification. The crude product dissolved in TFA:TIPS:water (95:2.5:2.5, 5 mL) and stirred for 2 h. After completion of the reaction (as indicated by LCMS), the volatiles were removed, and the crude product was purified by prep. HPLC to get title product (122 mg, 41%) as a white solid

LC-MS (ESI): (m/z) = 461 [M+H].

### UPLC-MS result of synthesized DOTA-TATE variants:

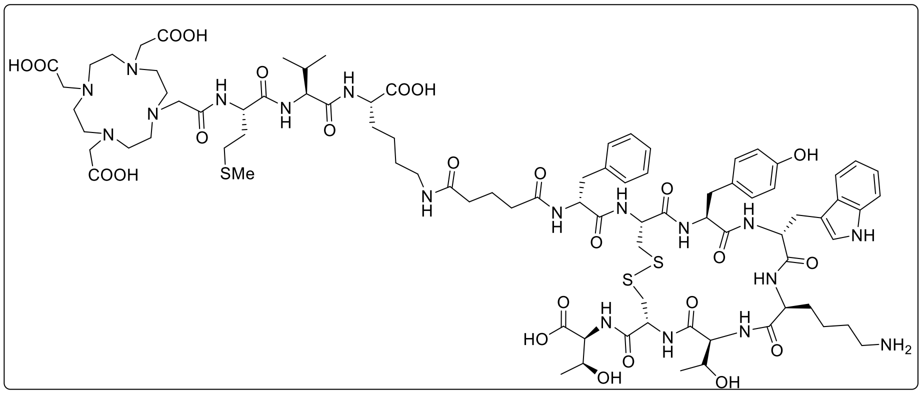


Exact Mass: 1906.83

DOTA-MVK(ε)-TATE

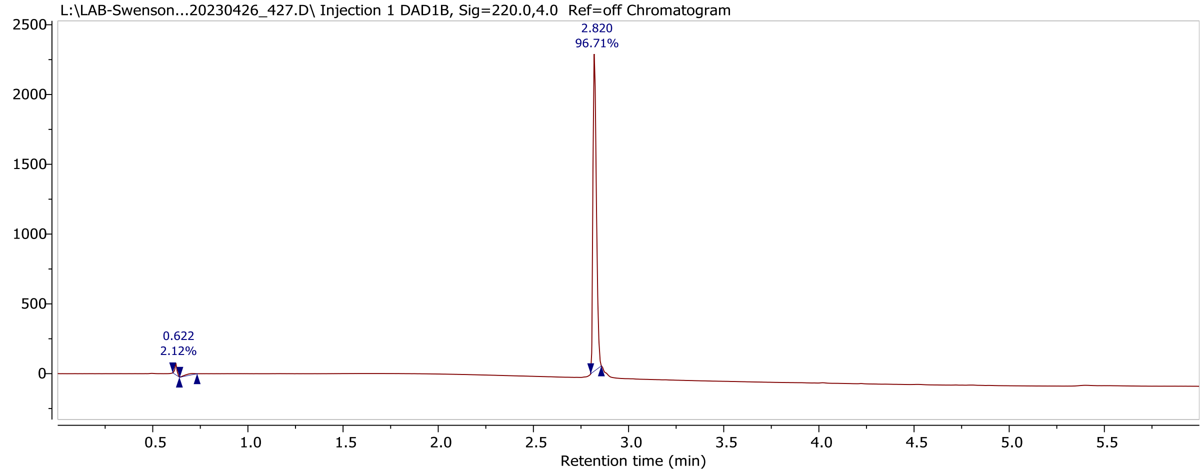


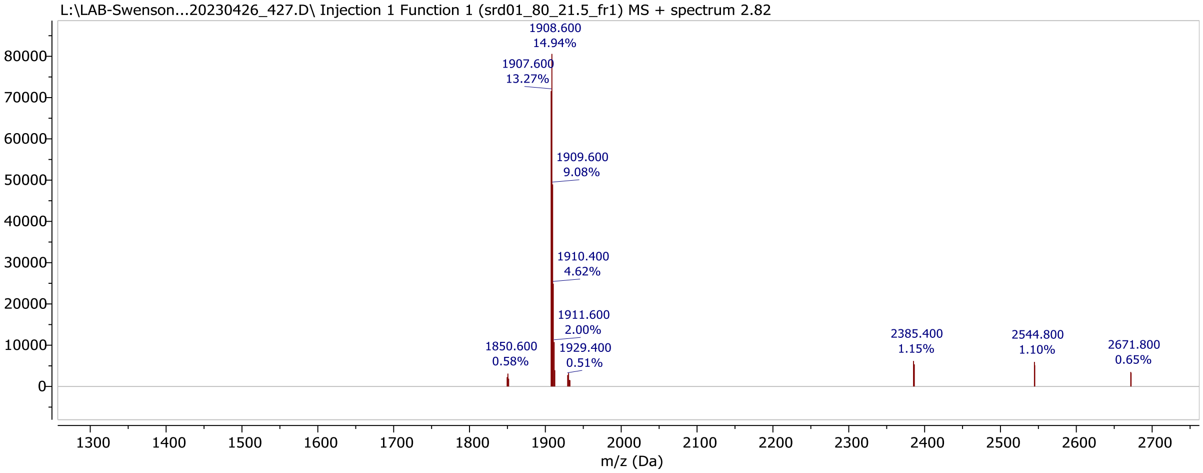


**Figure S1**. UPLC–MS chromatogram and mass spectrum of DOTA-MVK(ε)-TATE.

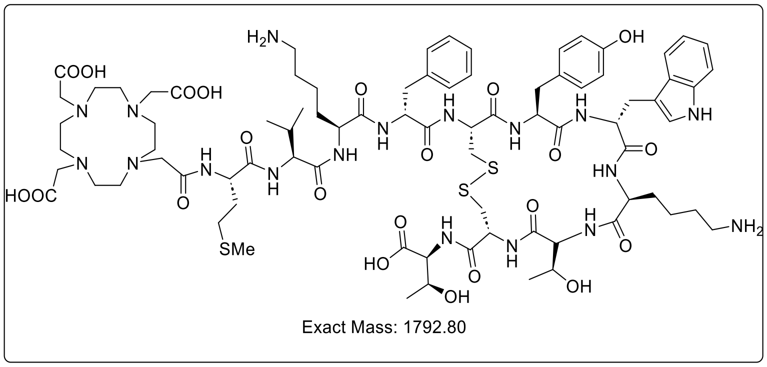


DOTA-MVK-TATE

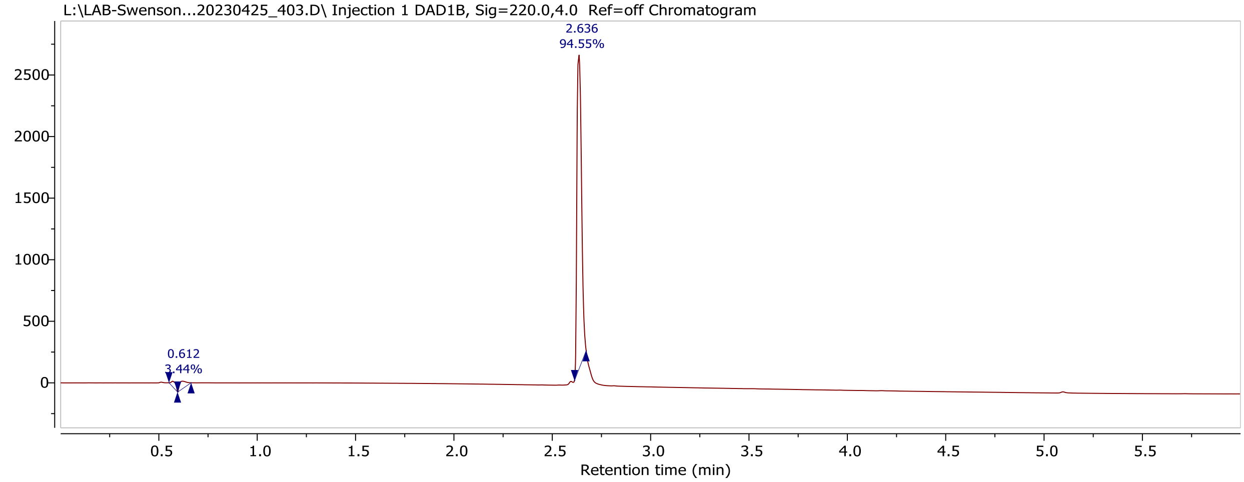


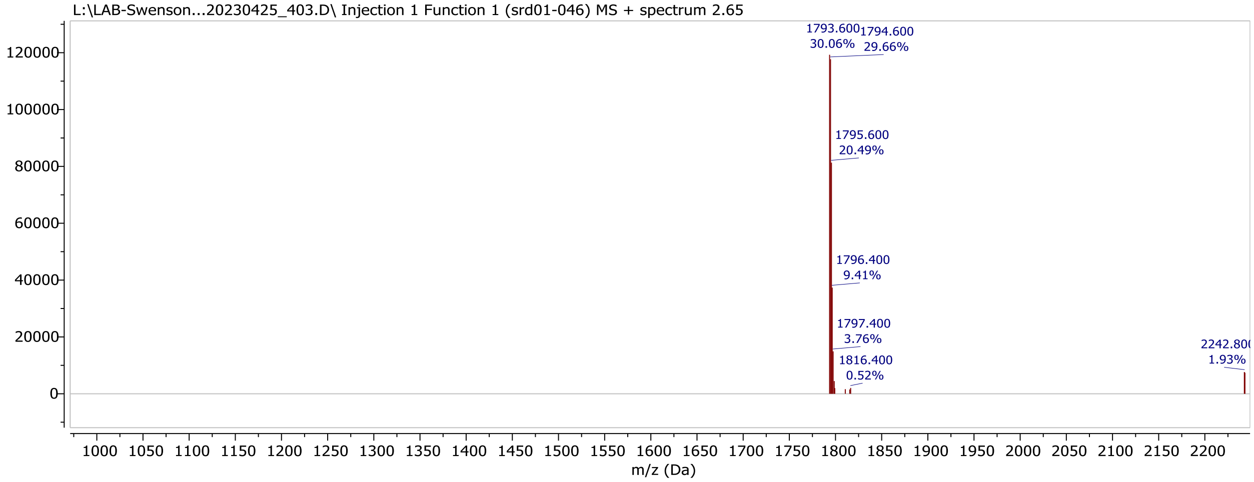


**Figure S2**. UPLC–MS chromatogram and mass spectrum of DOTA-MVK-TATE.

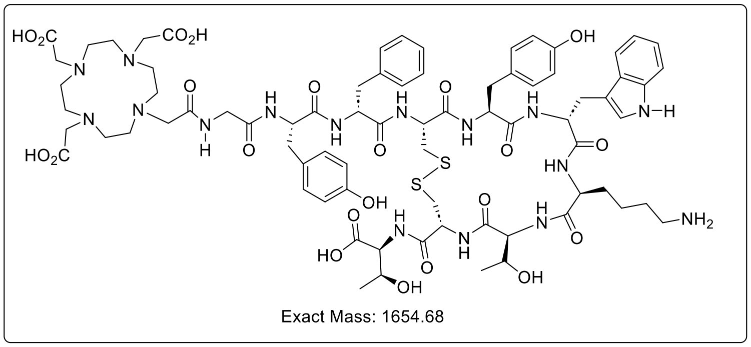


DOTA-GY-TATE

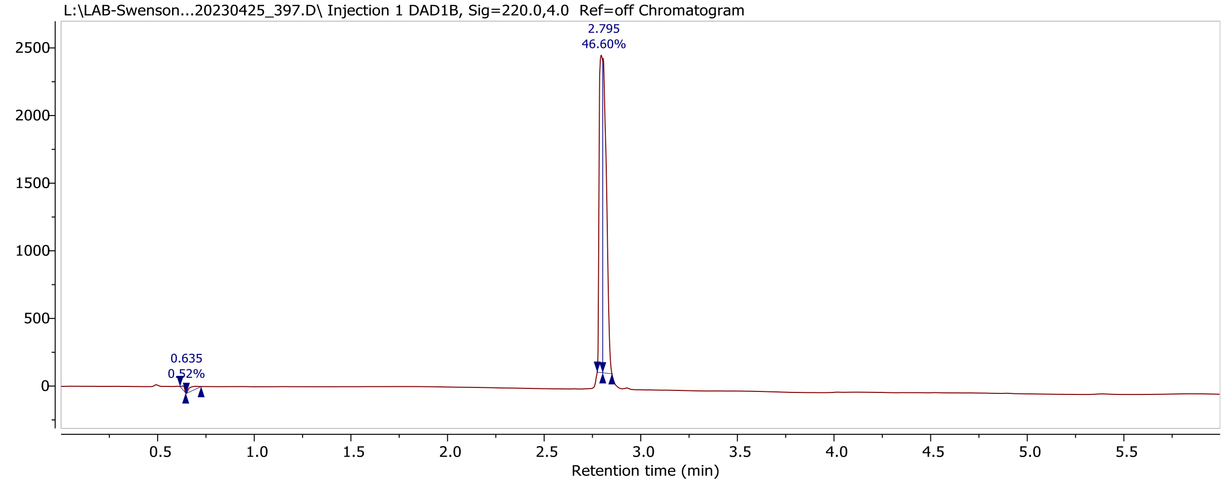


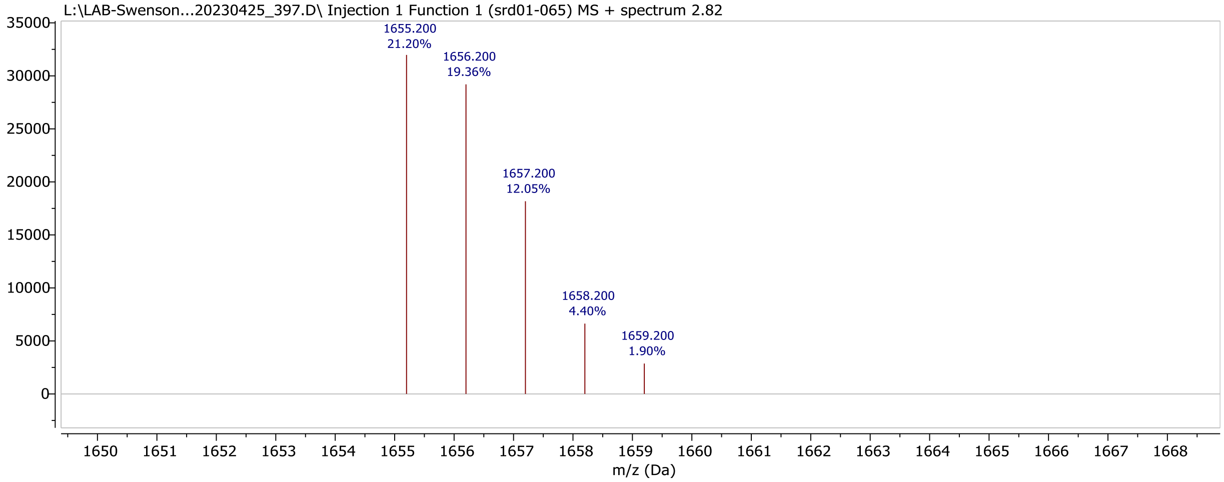


**Figure S3**. UPLC–MS chromatogram and mass spectrum of DOTA-GY-TATE.

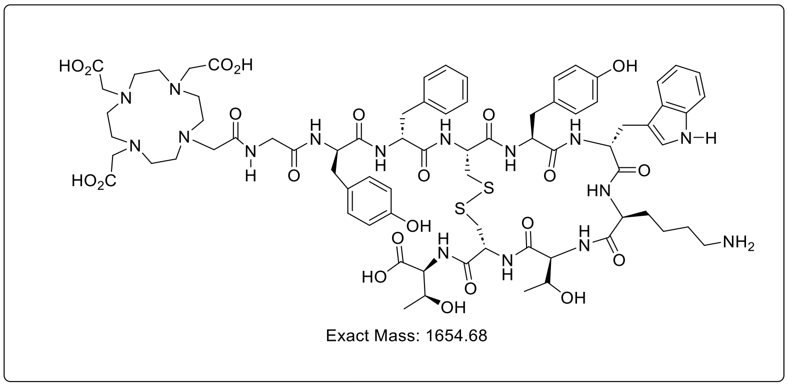


DOTA-GY*-TATE

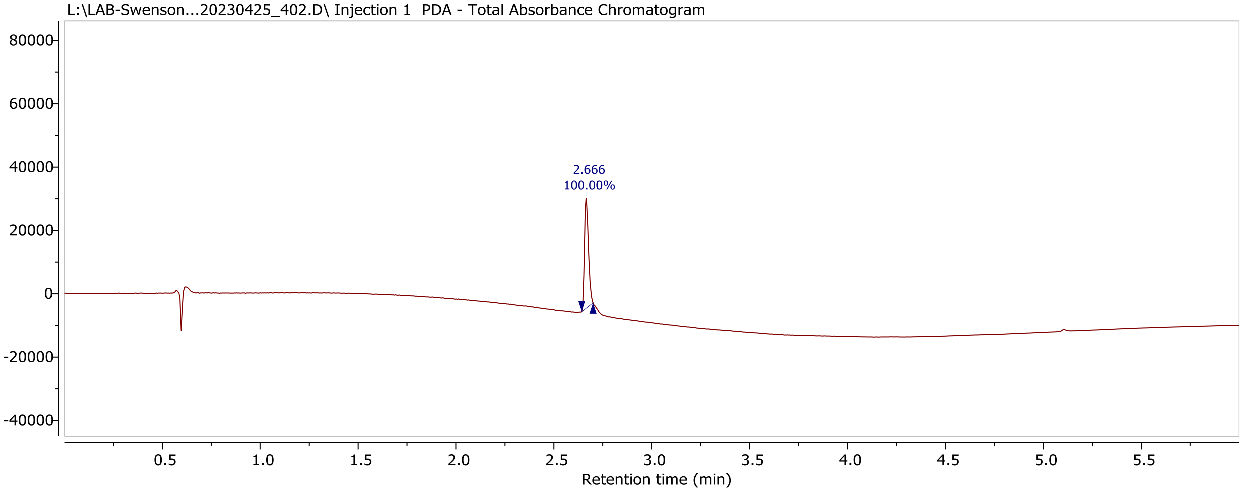


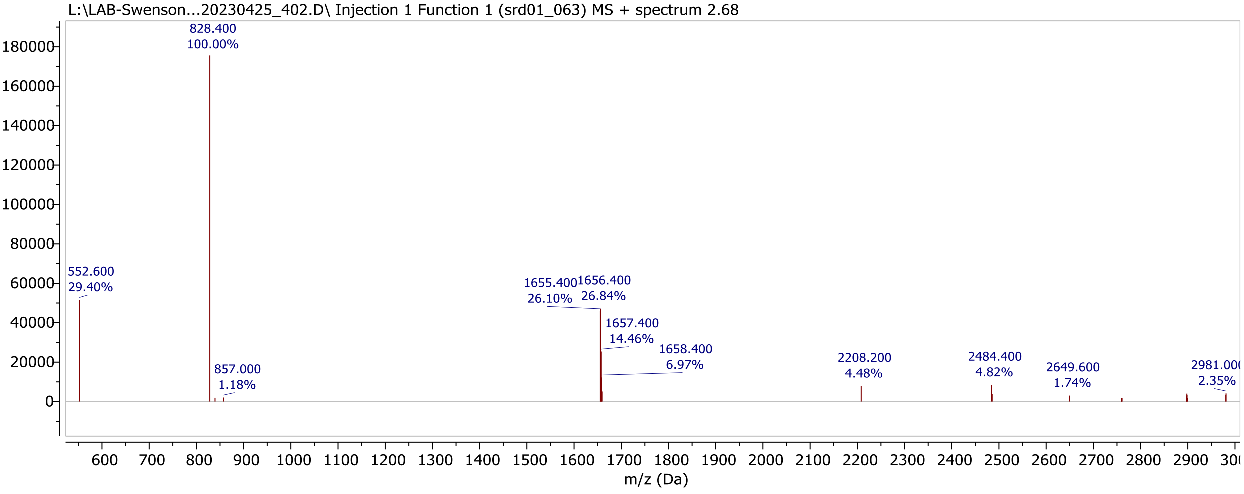


**Figure S4**. UPLC–MS chromatogram and mass spectrum of DOTA-GY*-TATE.

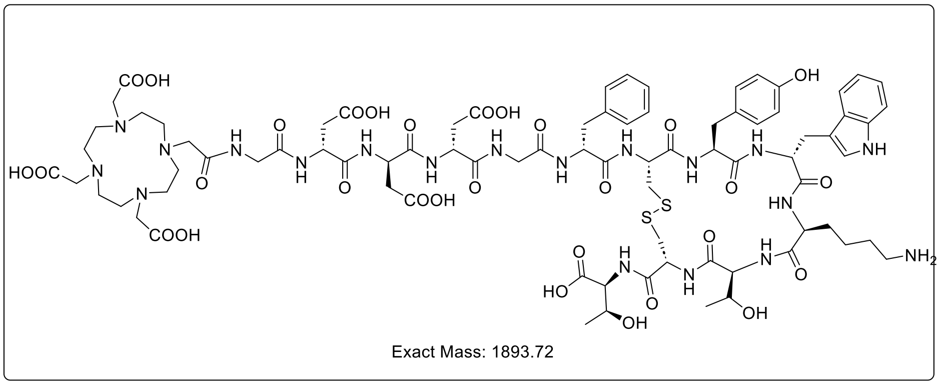


DOTA-GDDDG-TATE

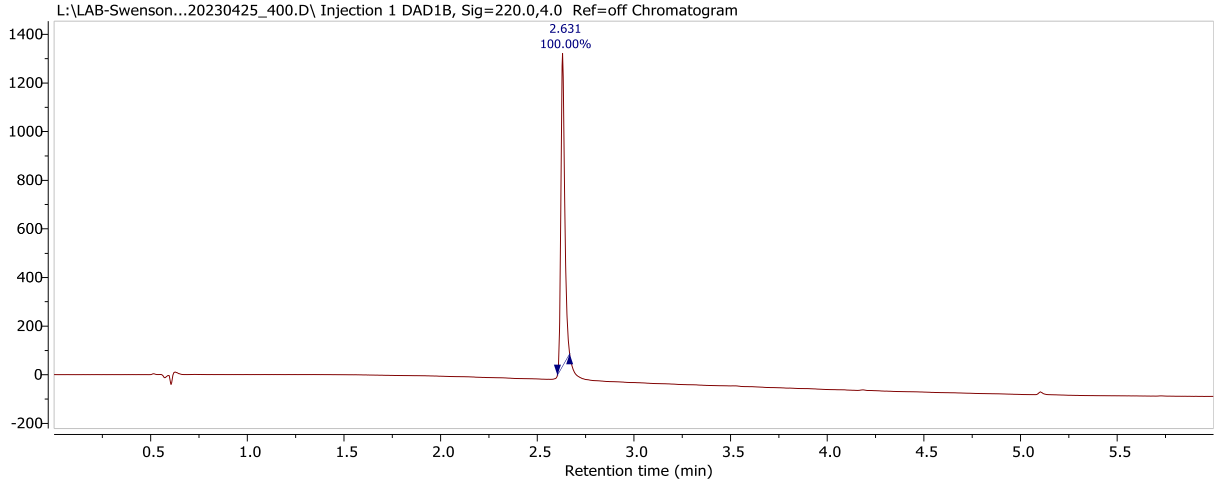


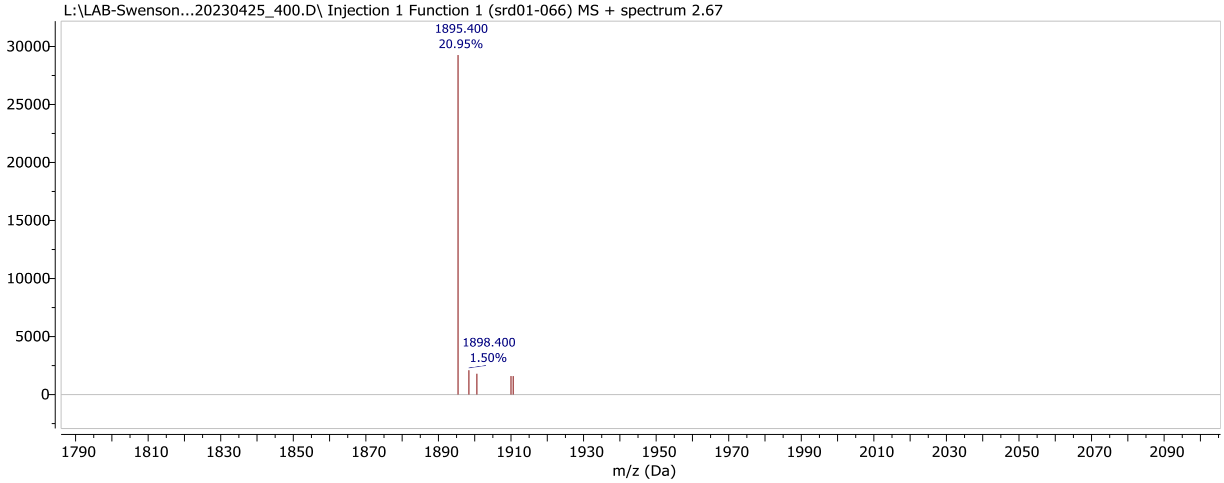


**Figure S5**. UPLC–MS chromatogram and mass spectrum of DOTA-GDDDG-TATE.

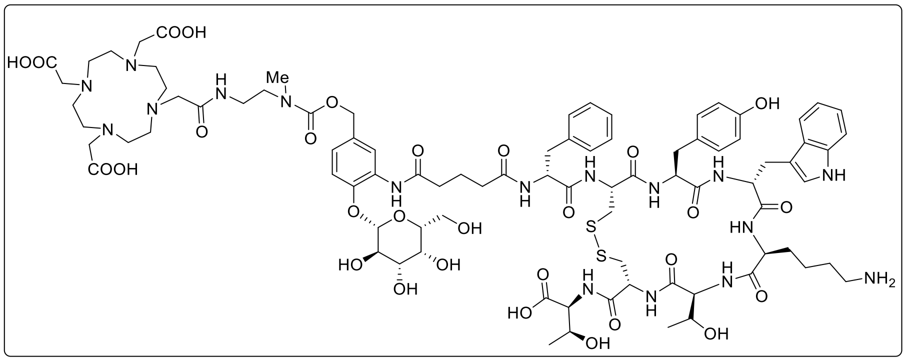


Exact Mass: 1931.80

DOTA-β-Gal-TATE

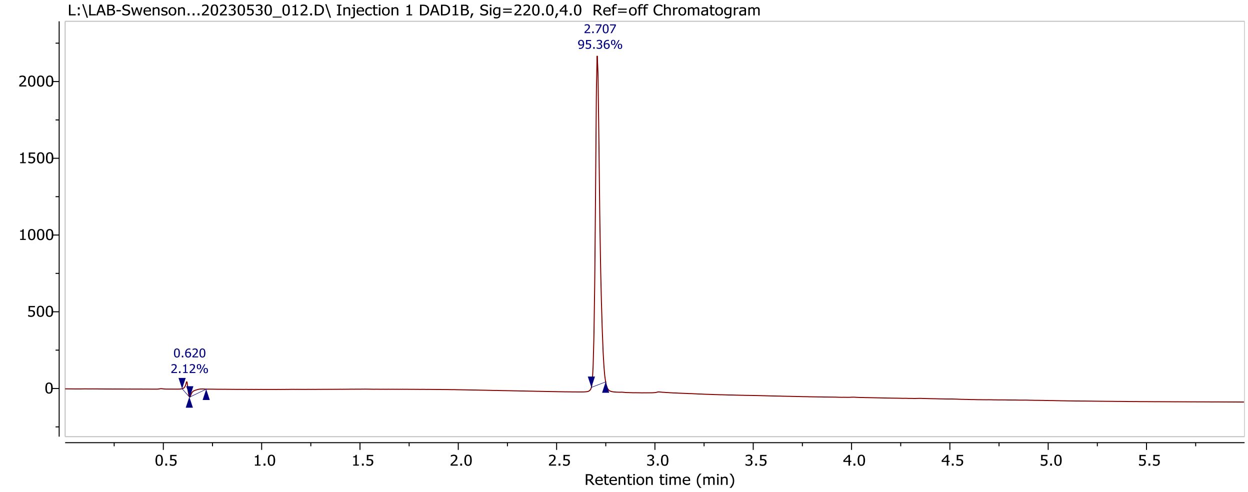


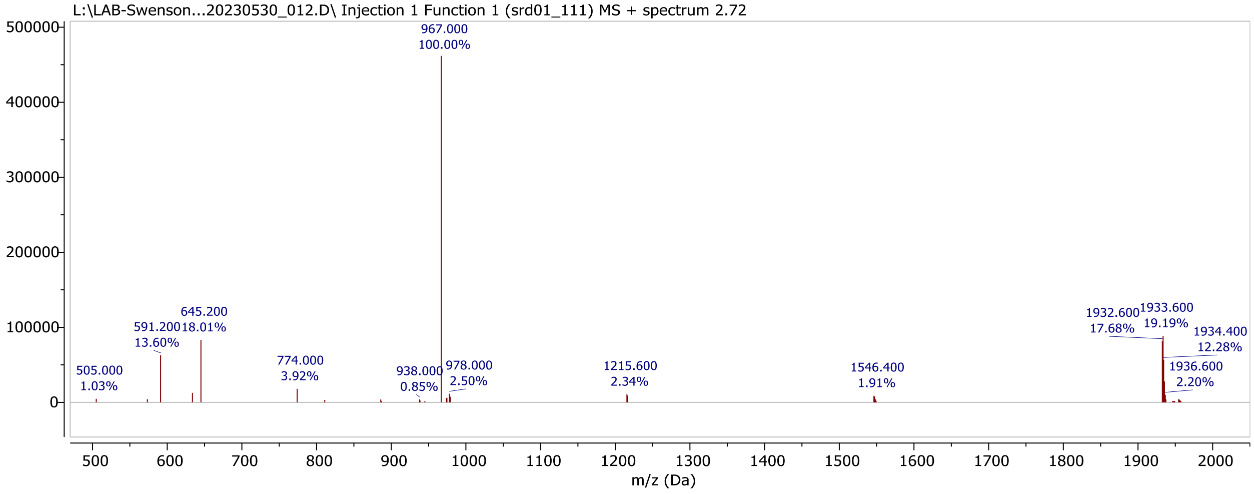


**Figure S6**. UPLC–MS chromatogram and mass spectrum of DOTA-β-Gal-TATE.

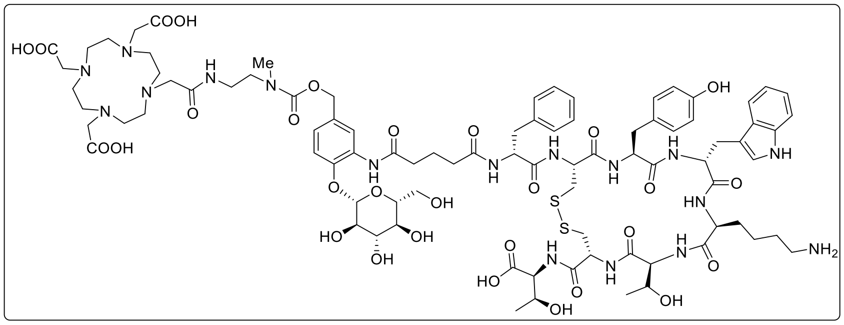

Exact Mass: 1931.80

DOTA-β-Glu-TATE

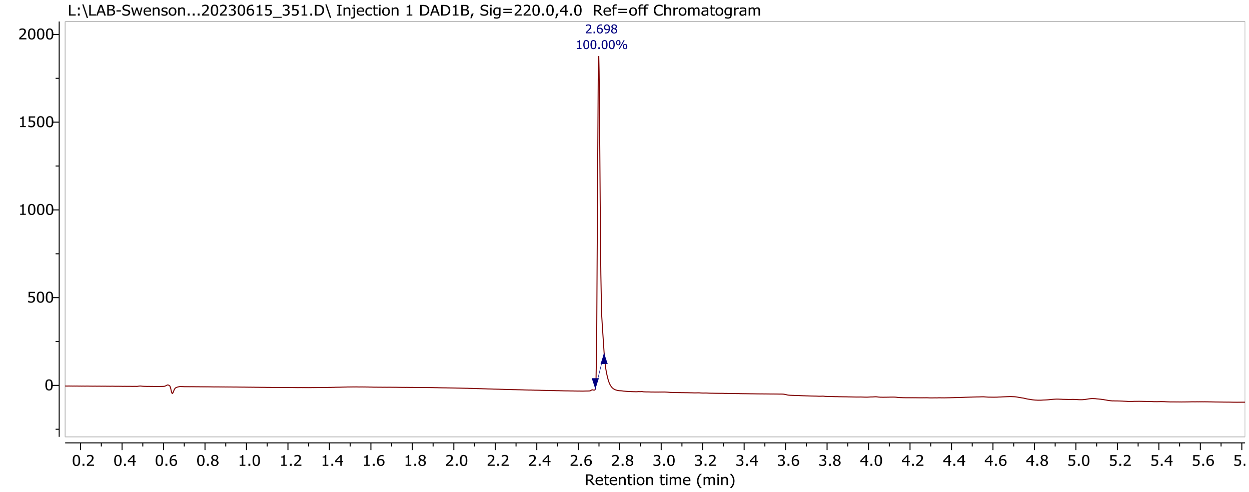


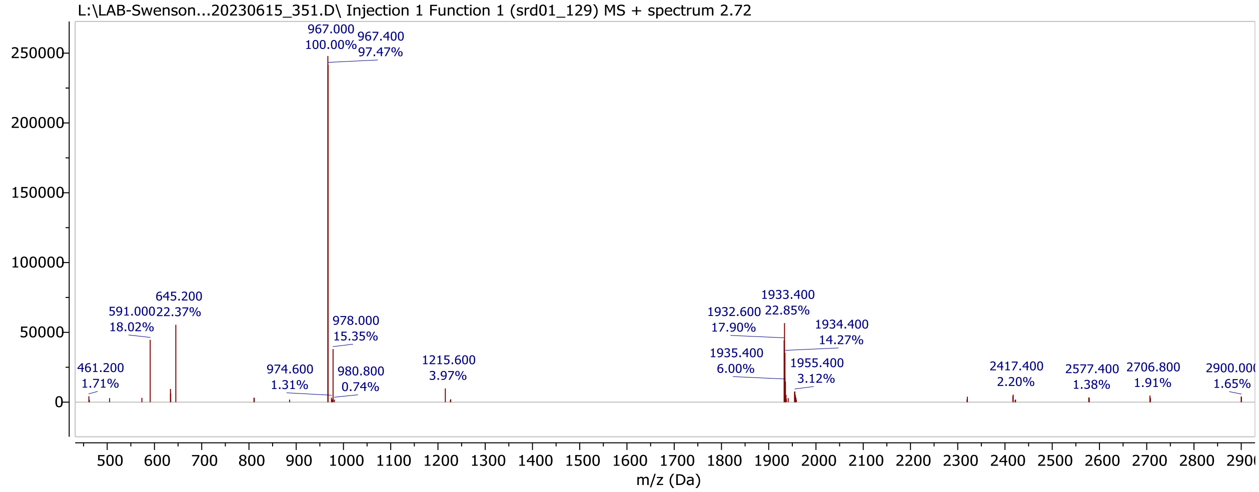


**Figure S7**. UPLC–MS chromatogram and mass spectrum of DOTA-β-Glu-TATE.

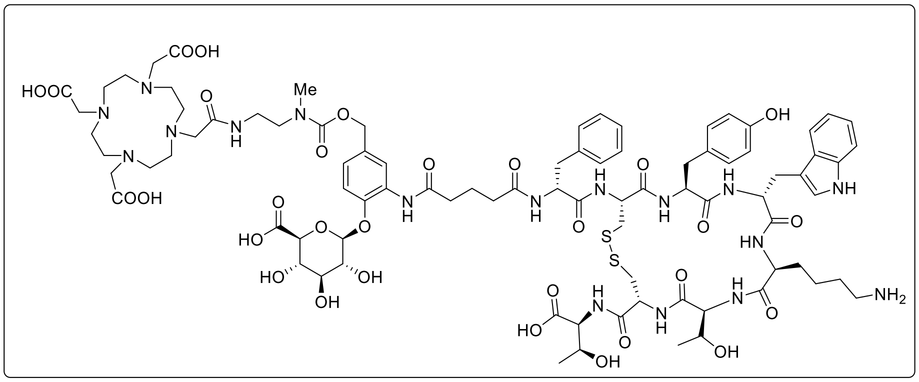

Exact Mass: 1945.78

DOTA-β-GlcA-TATE:

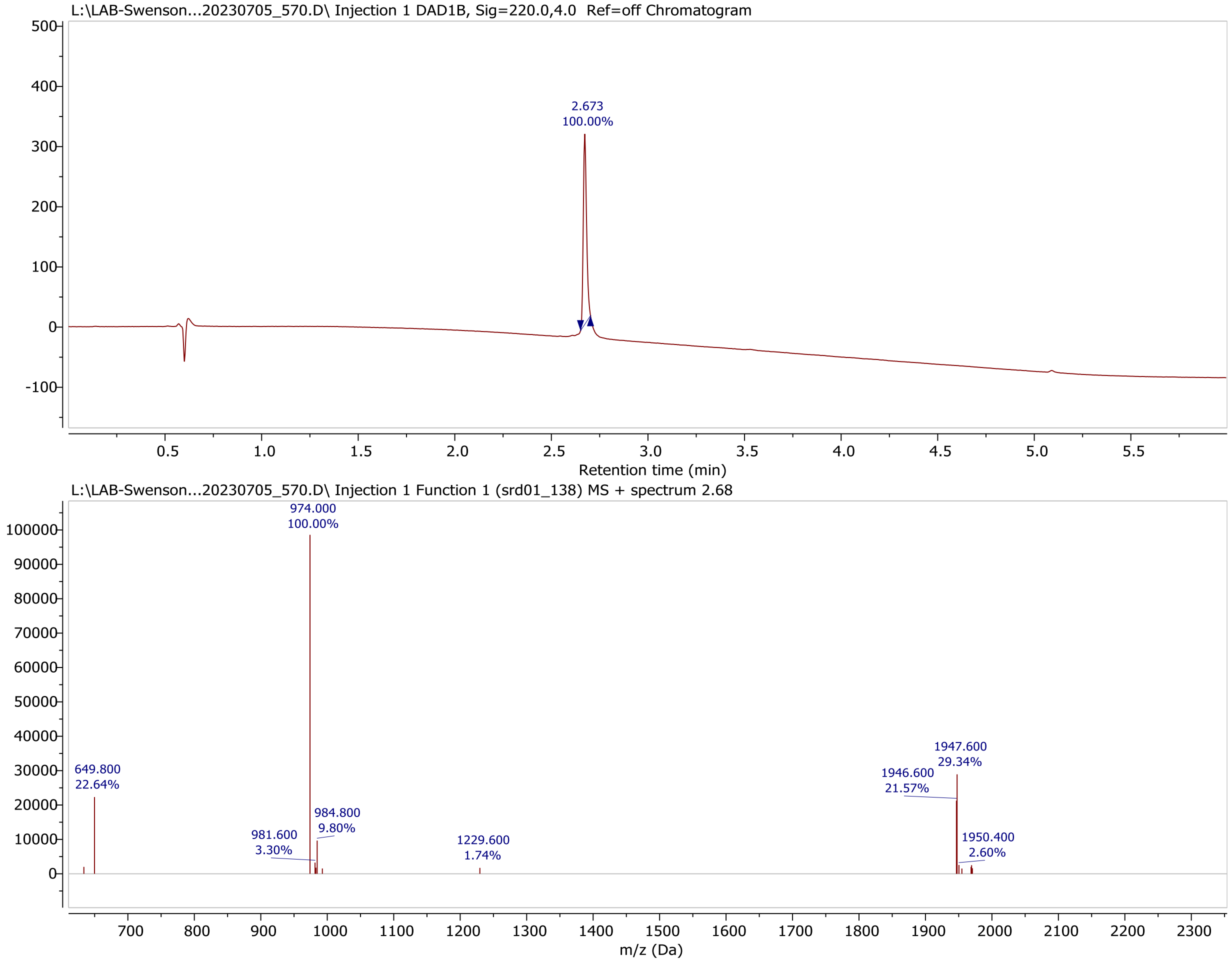


**Figure S8**. UPLC–MS chromatogram and mass spectrum of DOTA-β-GlcA-TATE.

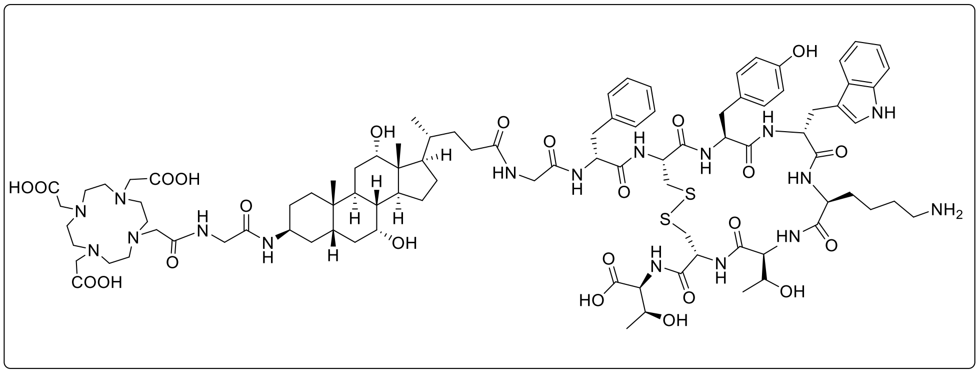


Exact Mass: 1937.93

DOTA-G-Chol-G-TATE

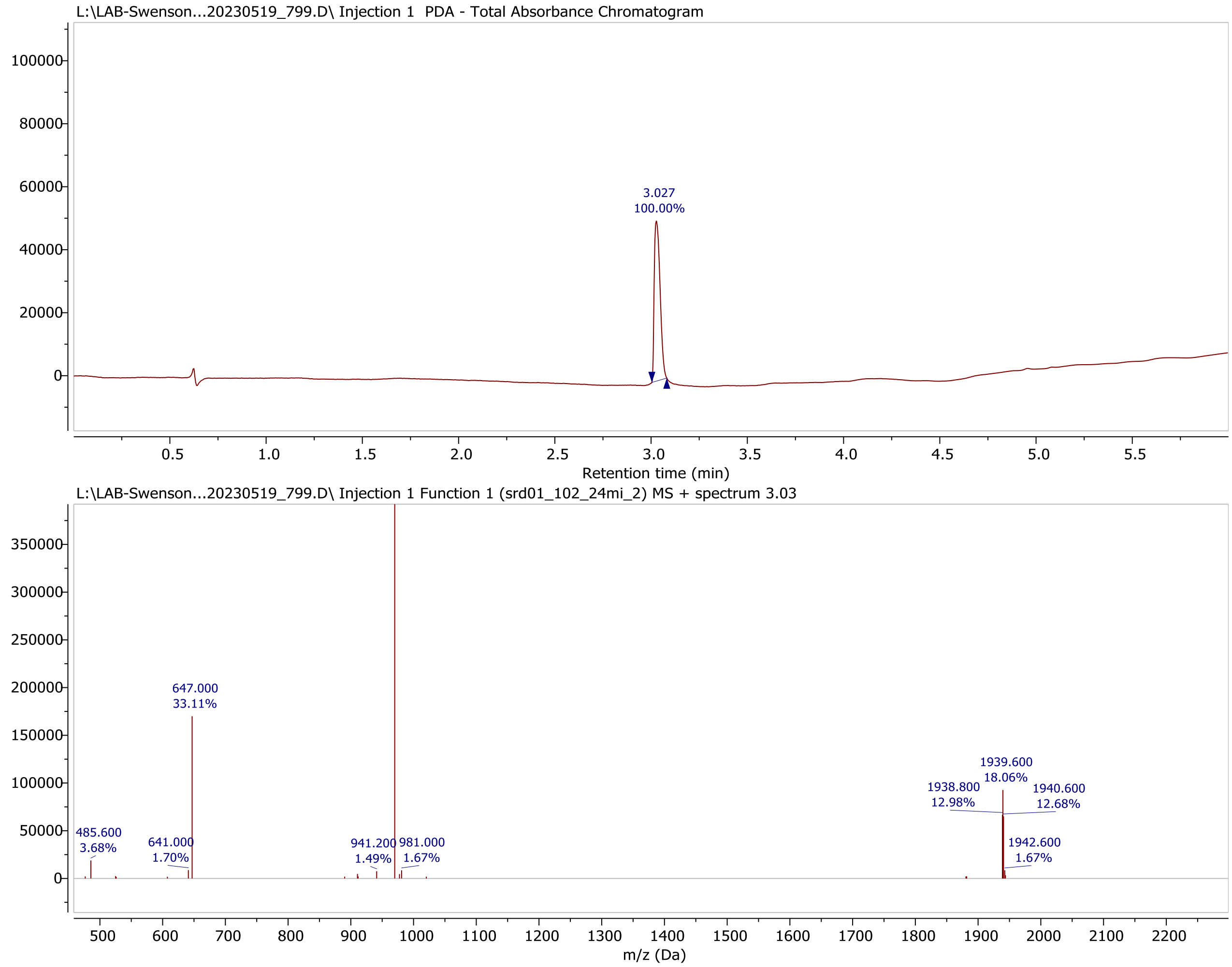


**Figure S9**. UPLC–MS chromatogram and mass spectrum of DOTA-G-Chol-G-TATE.

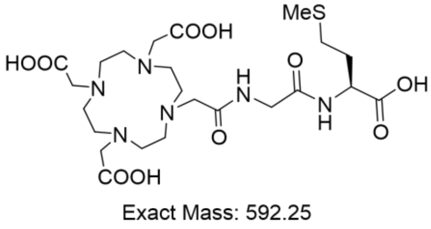


DOTA-G-M-OH

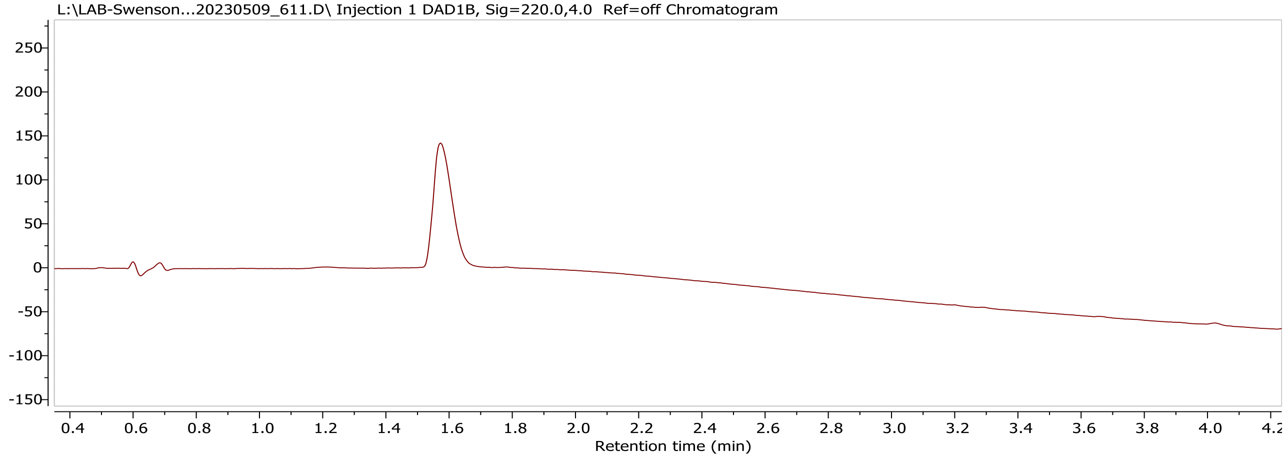


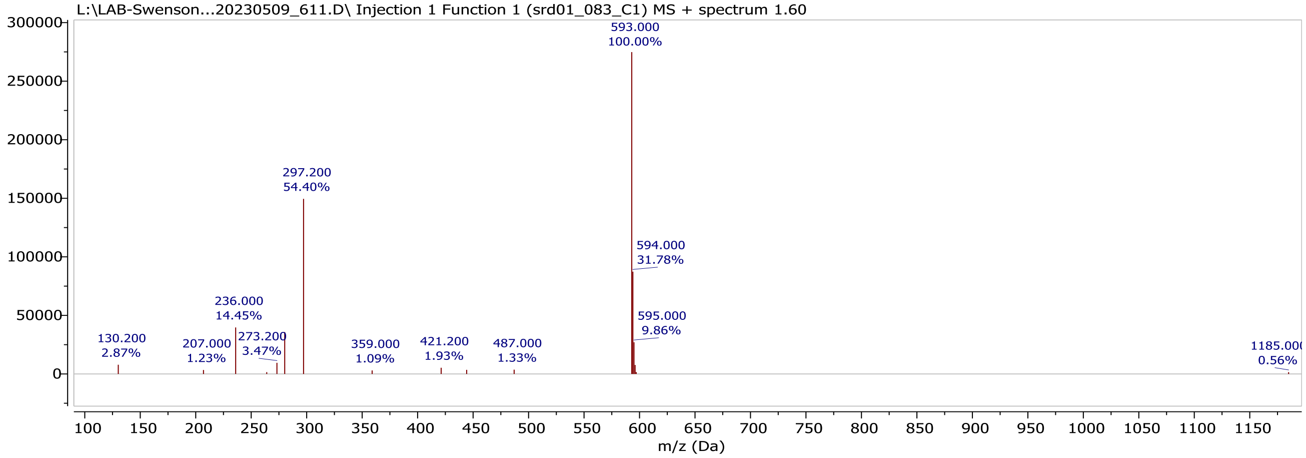


**Figure S10**. UPLC–MS chromatogram and mass spectrum of DOTA-G-M-OH.

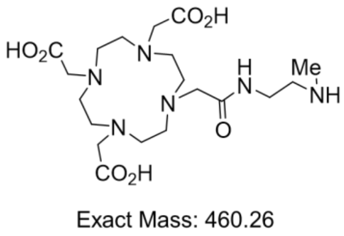


DOTA-NH-NHMe

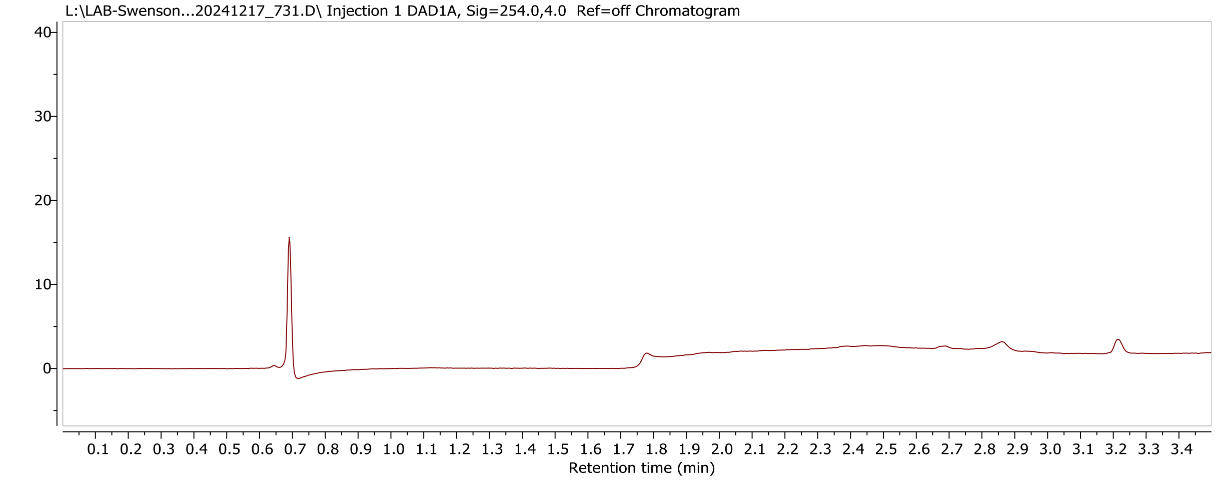


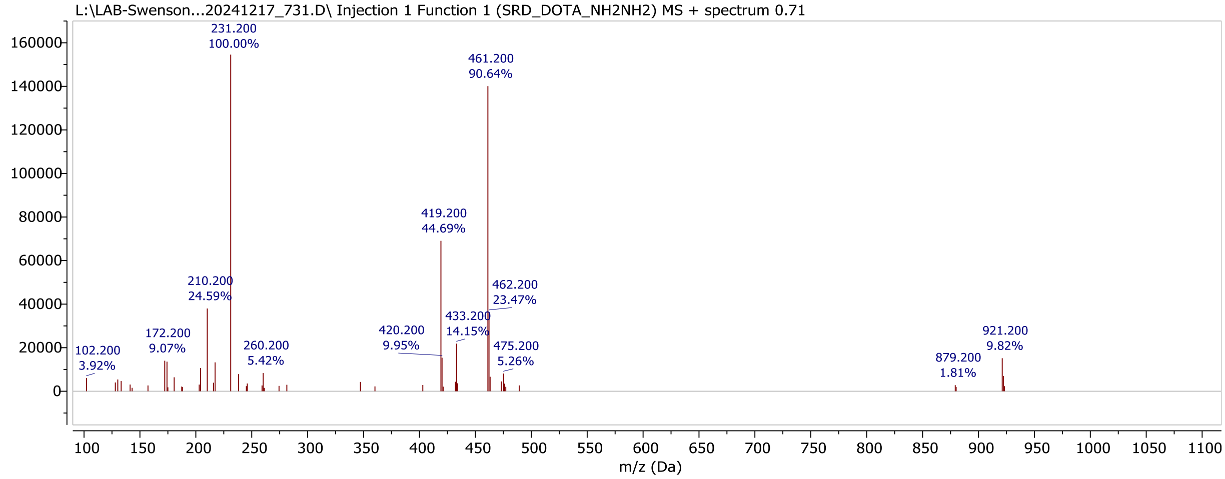


**Figure S11**. UPLC–MS chromatogram and mass spectrum of DOTA-NH-NHMe.

### Radiolabeling and In Vivo Evaluation of DOTA-TATE Variants: Biodistribution, PET Quantification, and Dosimetry

**Radiolabeling:** To perform the initial biodistribution screen, radiolabeling of 10 compounds, including DOTA-TATE, was carried out using $^{111}$In (Cardinal Health, Arbutus, MD, USA), an affordable and readily available gamma-emitting radioisotope with a moderate half-life (2.83 days), which affords efficient DOTA radiolabeling and facilitates handling. $^{111}InCl_3$ (28 – 36 MBq) was diluted with 0.4 M $NH_4OAc$ buffer (pH 5.2) and combined with DOTA-TATE variants (20 µg). The solution was mixed well and then incubated at 90˚C for 30 min. Radiolabeling efficiency and radiochemical purity were determined by radio-TLC and radio-HPLC. Radiochemical yield was >95% and the material was used without further purification. The specific activity was 1.55 ± 0.12 TBq/g (41.9 ± 3.2 Ci/g).

In addition, to the DOTA-TATE reference compound, two agents from the biodistribution screen, DOTA-MVK(ε)-TATE and DOTA-β-Gal-TATE, were selected for further investigation.

To assess tumor and kidney uptake using PET imaging in a tumor model and to compare these findings with DOTA-TATE, radiolabeling was performed with the positron-emitting radioisotope $^{86}$Y ($t_{1/2}$ = 14.7 h, National Institutes of Health Cyclotron Facility, Bethesda, MD, USA). DOTA-TATE variants (15 - 30 µg) were added to the $^{86}$Y solution (46 – 104 MBq in 1 M $HNO_3$) containing 2,5-dihydroxybenzoic acid (10 µL, 5 mg/mL) that had been neutralized to pH 5 with 5 M $NH_4OAc$ buffer (pH 7). The solution was mixed well and then incubated at 90˚C for 30 min. Radiolabeling efficiency and radiochemical purity was determined by radio-TLC and radio-HPLC. Like the $^{111}$In-labeled variants, the overall radiochemical yield and purity for the $^{86}$Y-labeled compounds were >95% and were used without additional purification. The specific activity was 3.85 ± 1.11 TBq/g (104 ± 30 Ci/g).

To perform targeted alpha particle therapy studies, $^{225}$Ac-labeled versions of three variants (DOTA-TATE, DOTA-MVK(ε)-TATE, and DOTA-β-Gal-TATE) were prepared. DOTA-TATE variants (15 - 20 µg) were added to an $^{225}$Ac solution (2.1 – 3.2 MBq in 0.1 M $HNO_3$, Oak Ridge National Laboratory, Oak Ridge, TN, USA) containing 2,5-dihydroxybenzoic acid (10 µL, 5 mg/mL), which had been neutralized to pH 5 using a 5 M $NH_4OAc$ buffer (pH 7). The solution was mixed thoroughly and incubated at 90°C for 30 min. Radiolabeling efficiency was determined by radio-TLC and radio-HPLC. The overall radiochemical yield was greater than 95%, and the material was used without further purification. The specific activity was 140 ± 23 GBq/g (3.78 ± 0.63 Ci/g).

Radio-instant thin-layer chromatography (radio-iTLC) was performed using silicic acid-impregnated glass microfiber paper strips (iTLC-SA, Varian, Lake Forest, CA, USA) as the stationary phase, and 50 mM EDTA in 100 mM $NH_4OAc$ (pH 5.5) as the mobile phase, analyzed with a radio-TLC scanner (AR-2000, Eckert-Ziegler, Wilmington, MA, USA). For Ac-225, equilibration was allowed for at least 12 hours before scanning the TLC strips to confirm initial scans. Radio-HPLC analysis (Agilent 1260 Infinity II, Agilent Technologies Inc., Santa Clara, CA, USA) was conducted on a TSK- Octadecyl-4PW column (5 µm, 4.6 mm ID × 15 cm, Tosoh Co. Tokyo, Japan) with gradient elution using 0.1% trifluoroacetic acid in water as mobile phase A and 0.1% trifluoroacetic acid in acetonitrile as Mobile phase B. The elution sequence was 0-5 min of 5% B, 5-10 min of 5-50% B, 10-20 min of 50-100% B, 20-30 min of 100-5% B at a flow rate of 0.75 mL/min.

**PET imaging and biodistribution:** For initial biodistribution screen studies, normal mice were administered 1.4 – 1.8 MBq (1 μg) of $^{111}$In-DOTA-TATE variants via tail vein injection and biodistribution studies were conducted at 2 hours post-injection (n = 2 – 4 per cohort).

For studies involving three variants (DOTA-TATE, DOTA-MVK(ε)-TATE, and DOTA-β-Gal-TATE), AR42J cells (5 × $10^6$ cells per mouse) were subcutaneously inoculated into the right shoulder of the mice. Tumors were allowed to grow for 3 weeks, reaching a volume of 100–200 $mm^3$ before being used in subsequent experiments. For biodistribution or PET imaging studies, mice bearing subcutaneous AR42J tumors were administered 1.4 – 1.8 MBq (1 μg) of $^{111}$In-labeled DOTA-TATE variants or 2.7 – 4.9 MBq (1 μg) of $^{86}$Y-labeled DOTA-TATE variants via tail vein injection. Biodistribution studies with $^{111}$In-labeled DOTA-TATE variants were performed 1 hour post-injection (n = 3 per cohort), while studies with $^{86}$Y-labeled DOTA-TATE variants were conducted at 1, 4, and 24 hours post-injection (n = 3 per cohort). Mice were euthanized by $CO_2$ asphyxiation, and 10 tissues, including tumors, were collected. All samples were weighed and counted on a gamma counter (2480 Wizard[3], Perkin Elmer Inc., Waltham, MA, USA). Results are presented as percentage of injected activity per gram (%IA/g) and mean values ± standard deviations.

PET imaging to assess the uptake of three $^{86}$Y-labeled DOTA-TATE variants (2.7 – 4.9 MBq, 1 μg) in the kidney and tumor was performed at 1, 4, and 24 hours post-injection (n = 3 per cohort) in mice bearing AR42J tumors using the MRS*PET/CT 120 scanner (MR Solutions, Guildford, UK). PET/CT data acquisition was managed using Preclinical Scan software (version 4.2.4.0, MR Solutions). Mice were anesthetized with 2% isoflurane, and static PET scans were acquired over 10 min. CT scans were obtained for attenuation correction and anatomical co-registration. PET data were reconstructed using three-dimensional ordered-subsets expectation maximization, and were subsequently normalized, decay- and dead-time–corrected prior to analysis. PET/CT images were analyzed using MIM software (version 7.2.7, MIM Software Inc., Beachwood, OH, USA), and the quantitative evaluation of the maximum standardized uptake ($SUV_{max}$) and the average standardized uptake values ($SUV_{mean}$) in the kidney and tumor were also performed.

Prior to conducting alpha particle therapy, biodistribution studies with $^{225}$Ac-labeled DOTA-TATE variants (117 – 163 kBq, 1 μg) were performed at 24 hours and 4 days post-injection in athymic nu/nu mice bearing AR42J tumors (n = 4 – 5). Mice were euthanized by $CO_2$ asphyxiation, and 10 tissues, including tumors, were collected. All samples were weighed and counted on a gamma counter after equilibration for 12 hours. Results are presented as %IA/g and mean values ± standard deviations.

**Dosimetry:** The dosimetry was calculated by measuring the residence time per gram for each organ using the percent injected activity per gram (%IA/g) data collected for the biodistribution study. Because two biodistribution data sample groups were collected, one for the $^{86}$Y and the other for the $^{225}$Ac variant of the three DOTA-TATE variants (DOTA-TATE, DOTA-MVK(ε)-TATE, and DOTA-β-Gal-TATE) the %IA/g time activity curves (TAC) of the $^{86}$Y variant were merged with the $^{225}$Ac TACs. This was possible because of the difference in half lives between $^{86}$Y and $^{225}$Ac being 14.74 and 237.8 hours respectively setting the measurement time points to be 1, 4 and 24 hours for the $^{86}$Y ligands and 24, 96 hours for $^{225}$Ac. The full time activity curve is thus made up of the 1 and 4 hour time points of the $^{86}$Y %IA/g data points, an average of the common 24 hour time point of the $^{86}$Y and $^{225}$Ac %IA/g data points and the last 96 hour %IA/g time point of the $^{225}$Ac samples. The full 4 time points of 1, 4, 24, and 96 hours making up the TAC were then decayed by $^{86}$Y and $^{225}$Ac half-life's respectively to generate $^{86}$Y and $^{225}$Ac TAC from which the residence times were calculated.

The integration of the TAC was performed using trapezoidal interpolation between time points. The first time point assumed constant activity and a decay curve with the half life of the radioisotope was used to estimate activity beyond 96 hours. The average %IA/g was used for each tissue sample for each time point and the resulting standard deviation of the average was then used as the error estimate per time point to propagate through the residence time calculation.

Using the residence time for each TAC, the dose was calculated by looking up the dose per decay for each alpha, beta (positron and electron from nuclear beta decay) and electron emission listed in the nuclear data sheets provided by the Brookhaven National Laboratory National Nuclear Data Center. Since the $^{225}$Ac decays via a complicated decay chain, it is assumed that the dose from the daughters remains in the tissue assuming no biological dissemination once the $^{225}$Ac decays breaking the bonds of the DOTA-TATE ligand. The following mean dose per decay were used. For $^{225}$Ac, 27.58MeV/Bq-s for alpha emissions, 1.33MeV/Bq-s for beta plus electron emissions. For $^{86}$Y, 0.219 MeV/Bq-s for beta and electron emissions. Gamma emissions were ignored because of the small size of the tissue samples have small gamma absorption effects which would contribute negligibly to the dose calculation. Furthermore, dosimetry estimate packages like MIRDCalc do not offer mouse models in which proper gamma emission dose from inter organ activity can be calculated*(62)*.

To estimate the effective dose in mSv/MBq, a radiological factor of 5 was used to multiply the dose calculated from the alpha ionizations while a factor of 1 was used for the beta and electron ionizations to convert from mGy/MBq to mSv/MBq*(63, 64)*.

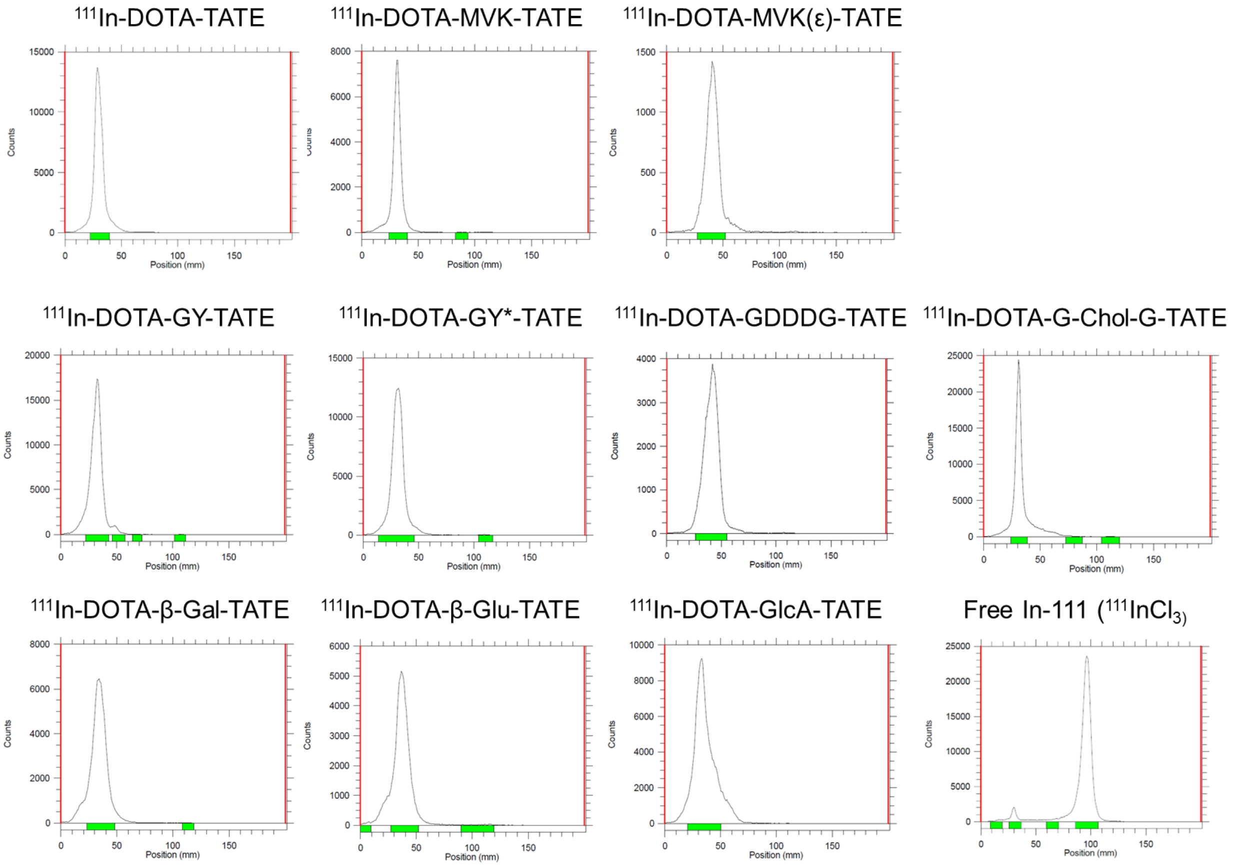


**Figure S12**. **Radio-TLC of $^{111}$In-labeled DOTA-TATE derivatives.** Radio-TLC was performed on ITLC-SA strips using 50 mM EDTA and 100 mM $NH_4OAc$ as the mobile phase. All $^{111}$In-labeled DOTA-TATE derivatives remained at the origin, in contrast to free $^{111}$In, which migrated with the solvent front.

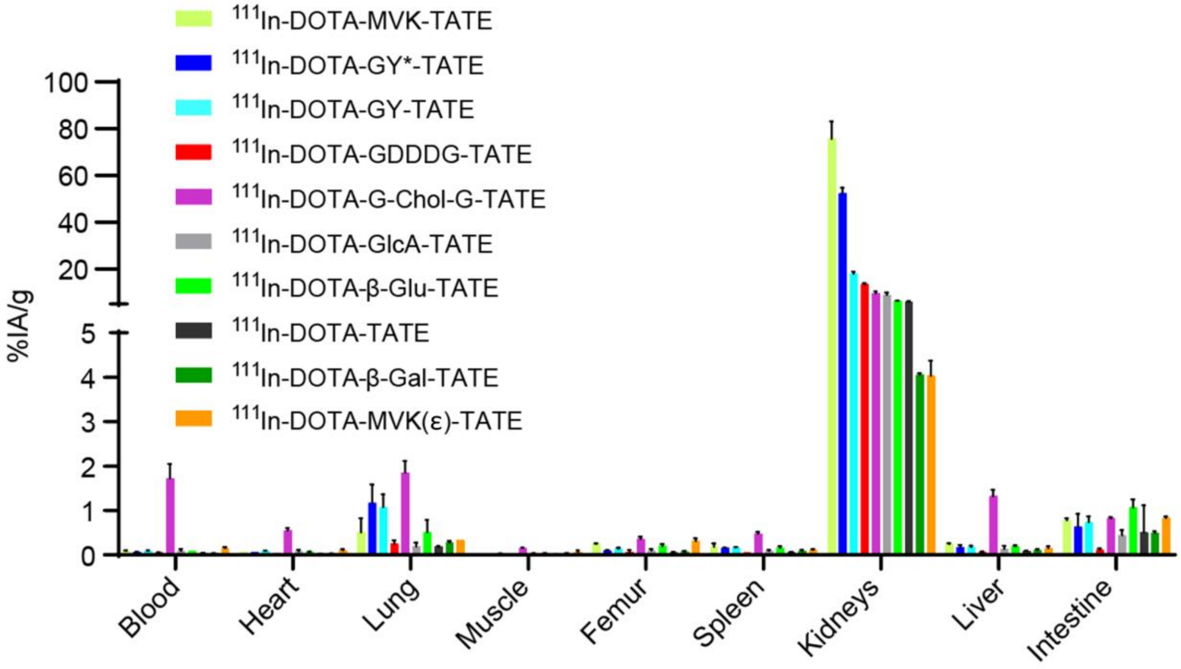


**Figure S13**. **Screening biodistribution of $^{111}$In-labeled DOTA-TATE derivatives in normal athymic mice.** Whole-organ radioactivity was measured at 2 post-injection to assess baseline distribution profiles. Data represent mean ± SD (n = 2 - 4 per cohort).

**Table S1. Screening biodistribution profiles of $^{111}$In-labeled DOTA-TATE derivatives in normal athymic mice.** Whole-organ radioactivity was quantified at 2 post-injection to evaluate baseline distribution patterns. Data are expressed as %IA/g (n = 2 - 4 per cohort).

| Organs | $^{111}$In-DOTA-MVK-TATE | | $^{111}$In-DOTA-GY*-TATE | | $^{111}$In-DOTA-GY-TATE | | $^{111}$In-DOTA-GDDDG-TATE | | $^{111}$In-DOTA-G-Chol-G-TATE | |
|---|---|---|---|---|---|---|---|---|---|---|
| | Average | SD | Average | SD | Average | SD | Average | SD | Average | SD |
| Blood | 0.09 | 0.01 | 0.06 | 0.01 | 0.09 | 0.02 | 0.05 | 0.01 | 1.72 | 0.32 |
| Heart | 0.07 | 0.00 | 0.06 | 0.00 | 0.09 | 0.00 | 0.04 | 0.00 | 0.55 | 0.05 |
| Lung | 0.51 | 0.32 | 1.17 | 0.42 | 1.07 | 0.30 | 0.25 | 0.07 | 1.86 | 0.26 |
| Muscle | 0.02 | 0.00 | 0.02 | 0.00 | 0.03 | 0.00 | 0.02 | 0.00 | 0.15 | 0.01 |
| Femur | 0.25 | 0.00 | 0.10 | 0.01 | 0.14 | 0.02 | 0.07 | 0.04 | 0.36 | 0.05 |
| Spleen | 0.18 | 0.08 | 0.16 | 0.00 | 0.17 | 0.01 | 0.05 | 0.00 | 0.48 | 0.03 |
| Kidney | 75.65 | 7.51 | 52.32 | 2.51 | 17.96 | 0.82 | 13.45 | 0.53 | 9.60 | 0.77 |
| Liver | 0.25 | 0.02 | 0.17 | 0.05 | 0.18 | 0.03 | 0.07 | 0.01 | 1.32 | 0.14 |
| Intestine | 0.78 | 0.05 | 0.63 | 0.29 | 0.74 | 0.13 | 0.11 | 0.03 | 0.83 | 0.02 |

| Organs | $^{111}$In-DOTA- GlcA-TATE | | $^{111}$In-DOTA-β-Glu-TATE | | $^{111}$In-DOTA-TATE | | $^{111}$In-DOTA-β-Gal-TATE | | $^{111}$In-DOTA-MVK(ε)-TATE | |
|---|---|---|---|---|---|---|---|---|---|---|
| | Average | SD | Average | SD | Average | SD | Average | SD | Average | SD |
| Blood | 0.08 | 0.05 | 0.09 | 0.00 | 0.04 | 0.00 | 0.04 | 0.00 | 0.15 | 0.03 |
| Heart | 0.08 | 0.04 | 0.07 | 0.01 | 0.03 | 0.00 | 0.03 | 0.00 | 0.11 | 0.02 |
| Lung | 0.19 | 0.09 | 0.51 | 0.28 | 0.18 | 0.02 | 0.28 | 0.03 | 0.33 | 0.00 |
| Muscle | 0.03 | 0.01 | 0.03 | 0.01 | 0.01 | 0.00 | 0.02 | 0.02 | 0.06 | 0.04 |
| Femur | 0.10 | 0.03 | 0.21 | 0.04 | 0.05 | 0.01 | 0.07 | 0.02 | 0.31 | 0.07 |
| Spleen | 0.08 | 0.03 | 0.16 | 0.04 | 0.06 | 0.01 | 0.08 | 0.03 | 0.12 | 0.01 |
| Kidney | 8.88 | 1.04 | 6.34 | 0.08 | 6.05 | 0.16 | 4.06 | 0.04 | 4.03 | 0.34 |
| Liver | 0.13 | 0.07 | 0.19 | 0.02 | 0.09 | 0.01 | 0.11 | 0.02 | 0.16 | 0.04 |
| Intestine | 0.44 | 0.12 | 1.07 | 0.18 | 0.51 | 0.61 | 0.49 | 0.04 | 0.83 | 0.04 |

**Table S2. Biodistribution of $^{111}$In-labeled DOTA-TATE derivatives at 1 h post-injection in AR42J tumor-bearing mice.** Radioactivity levels in tumors and major organs were measured at 1 h post-injection following intravenous administration of $^{111}$In-labeled DOTA-TATE derivatives. Data are expressed as %IA/g (n = 3 per cohort).

| Organs | $^{111}$In-DOTA-TATE | | $^{111}$In-DOTA-MVK(ε)-TATE | | $^{111}$In-DOTA-β-Gal-TATE | |
|---|---|---|---|---|---|---|
| | Average | SD | Average | SD | Average | SD |
| Blood | 0.27 | 0.06 | 0.73 | 0.14 | 0.56 | 0.06 |
| Tumor | 5.67 | 0.92 | 5.44 | 0.19 | 10.77 | 1.11 |
| Heart | 0.10 | 0.01 | 0.29 | 0.05 | 0.29 | 0.07 |
| Lung | 0.49 | 0.11 | 0.73 | 0.08 | 0.92 | 0.04 |
| Muscle | 0.06 | 0.01 | 0.27 | 0.09 | 0.17 | 0.03 |
| Femur | 0.12 | 0.02 | 0.55 | 0.03 | 0.34 | 0.03 |
| Spleen | 0.10 | 0.02 | 0.16 | 0.02 | 0.21 | 0.05 |
| Kidney | 7.33 | 0.04 | 5.56 | 0.29 | 5.19 | 0.18 |
| Liver | 0.23 | 0.07 | 0.31 | 0.03 | 0.45 | 0.01 |
| Intestine | 0.42 | 0.06 | 0.81 | 0.18 | 0.79 | 0.17 |

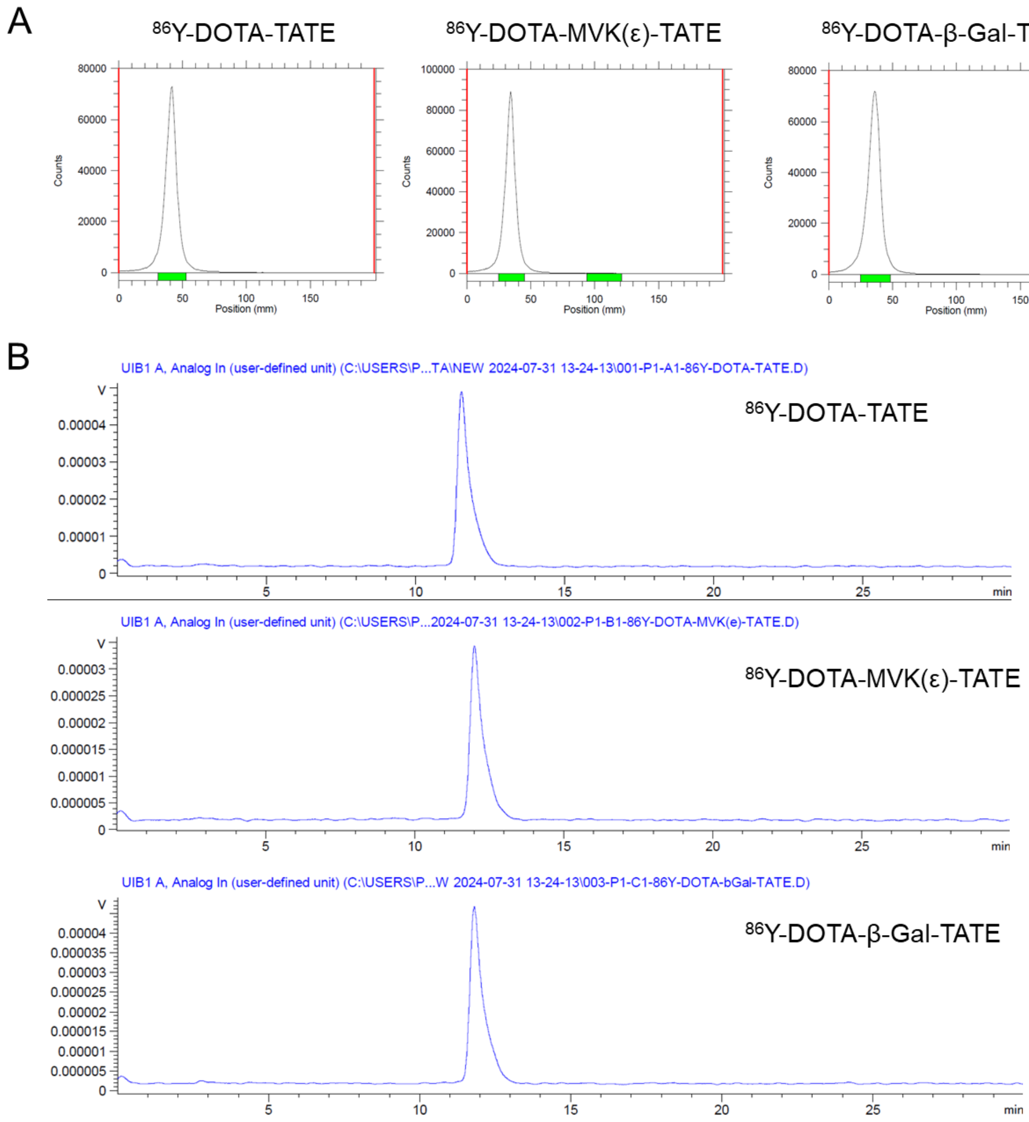


**Figure S14**. **Radio-TLC and radio-HPLC of $^{86}$Y-labeled DOTA-TATE derivatives.** (**A**) Radio-TLC was performed on ITLC-SA strips using 50 mM EDTA or 100 mM $NH_4OAc$ as the mobile phase. (**B**) Radio-HPLC analysis (Agilent 1260 Infinity II, TSK-Octadecyl-4PW column) was carried out with a water/acetonitrile gradient containing 0.1% trifluoroacetic acid. All derivatives exhibited a single radioactive peak in the radio-HPLC chromatogram, eluting between 11 and 12 min.

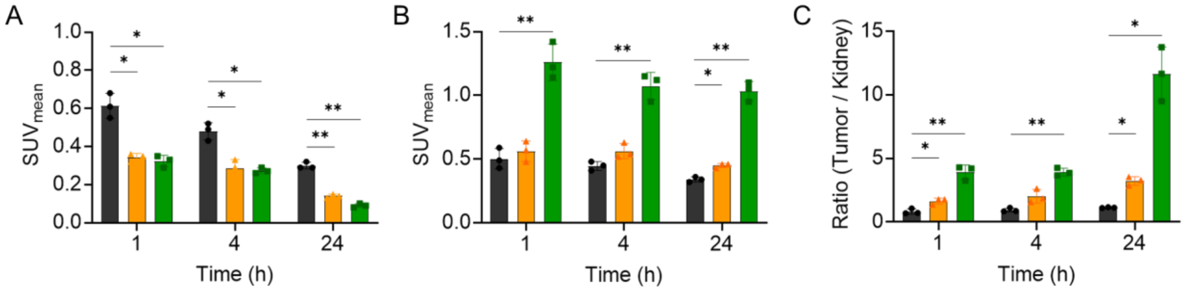


**Figure S15. PET quantification of $^{86}$Y-labeled DOTA-TATE variants in AR42J tumor-bearing mice.** Quantification was performed using mean standardized uptake values ($SUV_{mean}$) for (**A**) kidney, (**B**) tumor, and (**C**) tumor-to-kidney ratio at each time point. Data are presented as mean ± SD (n = 3 per cohort). Statistical significance was assessed using unpaired two-tailed t-tests (*$p < 0.05$, **$p < 0.01$).

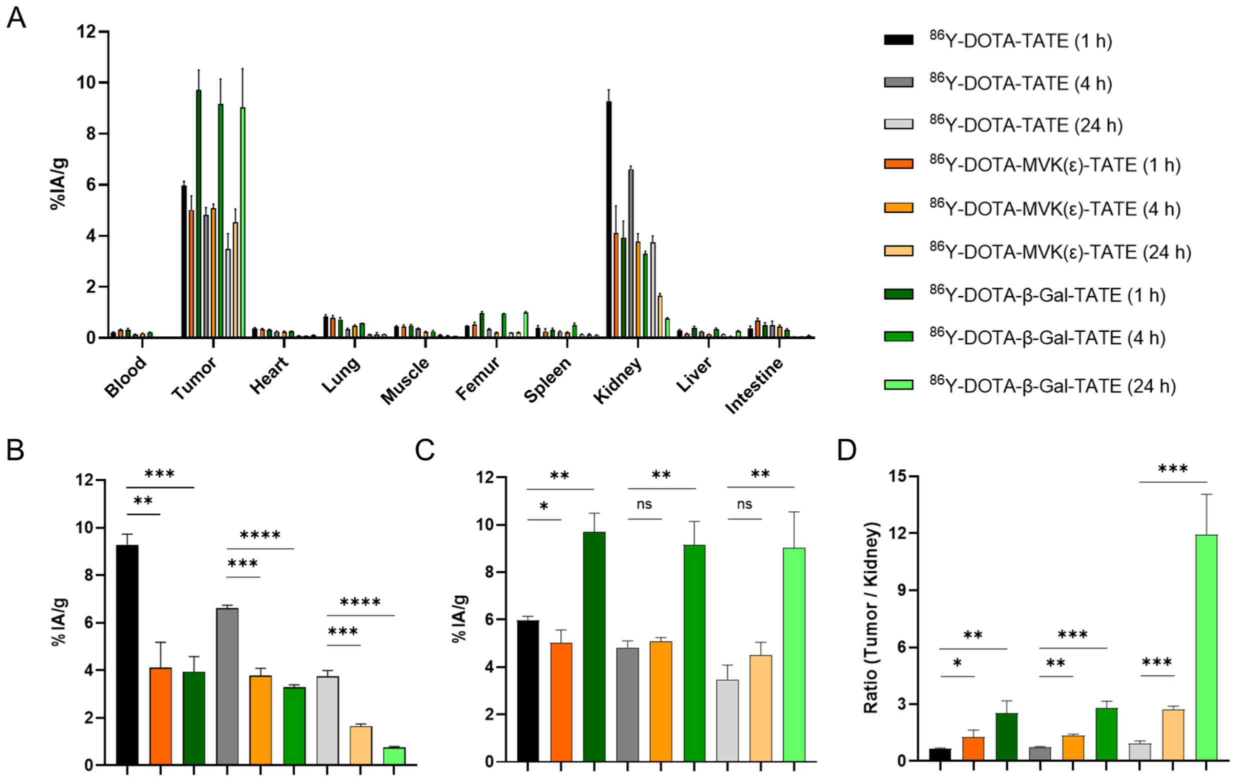


**Figure S16. Biodistribution of $^{86}$Y-labeled DOTA-TATE derivatives in AR42J tumor-bearing mice.** (**A**) Time-dependent biodistribution at 1, 4, and 24 h post-injection following intravenous administration of $^{86}$Y-labeled DOTA-TATE derivatives. (**B**) Kidney uptake across the three groups at each time point. (**C**) Tumor uptake across the three groups at each time point. (**D**) Tumor-to-kidney uptake ratios for each group at 1, 4, and 24 h. Data are presented as %IA/g (A–C) or ratios (D) and expressed as mean ± SD (n = 3 per cohort). Statistical significance was assessed using unpaired two-tailed t-tests: *$p < 0.05$, **$p < 0.01$, ***$p < 0.001$, ****$p < 0.0001$.

**Table S3. Biodistribution of $^{86}$Y-labeled DOTA-TATE derivatives at 1, 4, and 24 h post-injection in AR42J tumor-bearing mice.** Radioactivity in tumors and major organs at 1, 4, and 24 h post-injection of $^{86}$Y-labeled DOTA-TATE derivatives, expressed as %IA/g (n = 3).

| $^{86}$Y-DOTA-TATE | | | | | | |
|---|---|---|---|---|---|---|
| Organs | 1 h | | 4 h | | 24 h | |
| | Average | SD | Average | SD | Average | SD |
| Blood | 0.22 | 0.01 | 0.14 | 0.02 | 0.03 | 0.00 |
| Tumor | 5.98 | 0.15 | 4.83 | 0.28 | 3.48 | 0.61 |
| Heart | 0.37 | 0.03 | 0.25 | 0.02 | 0.09 | 0.01 |
| Lung | 0.83 | 0.08 | 0.34 | 0.04 | 0.13 | 0.02 |
| Muscle | 0.44 | 0.04 | 0.38 | 0.02 | 0.10 | 0.03 |
| Femur | 0.47 | 0.01 | 0.35 | 0.03 | 0.20 | 0.01 |
| Spleen | 0.40 | 0.09 | 0.25 | 0.03 | 0.15 | 0.01 |
| Kidney | 9.29 | 0.44 | 6.61 | 0.12 | 3.74 | 0.25 |
| Liver | 0.29 | 0.04 | 0.25 | 0.01 | 0.15 | 0.01 |
| Intestine | 0.37 | 0.10 | 0.51 | 0.15 | 0.05 | 0.00 |
| $^{86}$Y-DOTA-MVK(ε)-TATE | | | | | | |
| Organs | 1 h | | 4 h | | 24 h | |
| | Average | SD | Average | SD | Average | SD |
| Blood | 0.32 | 0.02 | 0.17 | 0.02 | 0.02 | 0.00 |
| Tumor | 5.02 | 0.56 | 5.08 | 0.17 | 4.53 | 0.52 |
| Heart | 0.35 | 0.03 | 0.24 | 0.03 | 0.08 | 0.01 |
| Lung | 0.79 | 0.10 | 0.47 | 0.05 | 0.13 | 0.09 |
| Muscle | 0.45 | 0.07 | 0.23 | 0.03 | 0.09 | 0.01 |
| Femur | 0.52 | 0.10 | 0.21 | 0.02 | 0.20 | 0.03 |
| Spleen | 0.25 | 0.11 | 0.22 | 0.02 | 0.14 | 0.03 |
| Kidney | 4.12 | 1.06 | 3.78 | 0.31 | 1.66 | 0.08 |
| Liver | 0.17 | 0.01 | 0.15 | 0.01 | 0.06 | 0.01 |
| Intestine | 0.69 | 0.09 | 0.46 | 0.05 | 0.05 | 0.01 |
| $^{86}$Y-DOTA-β-Gal-TATE | | | | | | |
| Organ | 1 h | | 4 h | | 24 h | |
| | Average | SD | Average | SD | Average | SD |
| Blood | 0.32 | 0.06 | 0.23 | 0.01 | 0.02 | 0.00 |
| Tumor | 9.71 | 0.79 | 9.16 | 0.99 | 9.03 | 1.53 |
| Heart | 0.33 | 0.02 | 0.26 | 0.01 | 0.11 | 0.02 |
| Lung | 0.72 | 0.08 | 0.58 | 0.01 | 0.15 | 0.01 |
| Muscle | 0.48 | 0.05 | 0.25 | 0.05 | 0.09 | 0.00 |
| Femur | 0.97 | 0.07 | 0.96 | 0.01 | 1.01 | 0.04 |
| Spleen | 0.32 | 0.07 | 0.50 | 0.08 | 0.12 | 0.01 |
| Kidney | 3.94 | 0.64 | 3.29 | 0.10 | 0.76 | 0.04 |
| Liver | 0.40 | 0.05 | 0.35 | 0.04 | 0.27 | 0.02 |
| Intestine | 0.49 | 0.11 | 0.33 | 0.04 | 0.10 | 0.02 |

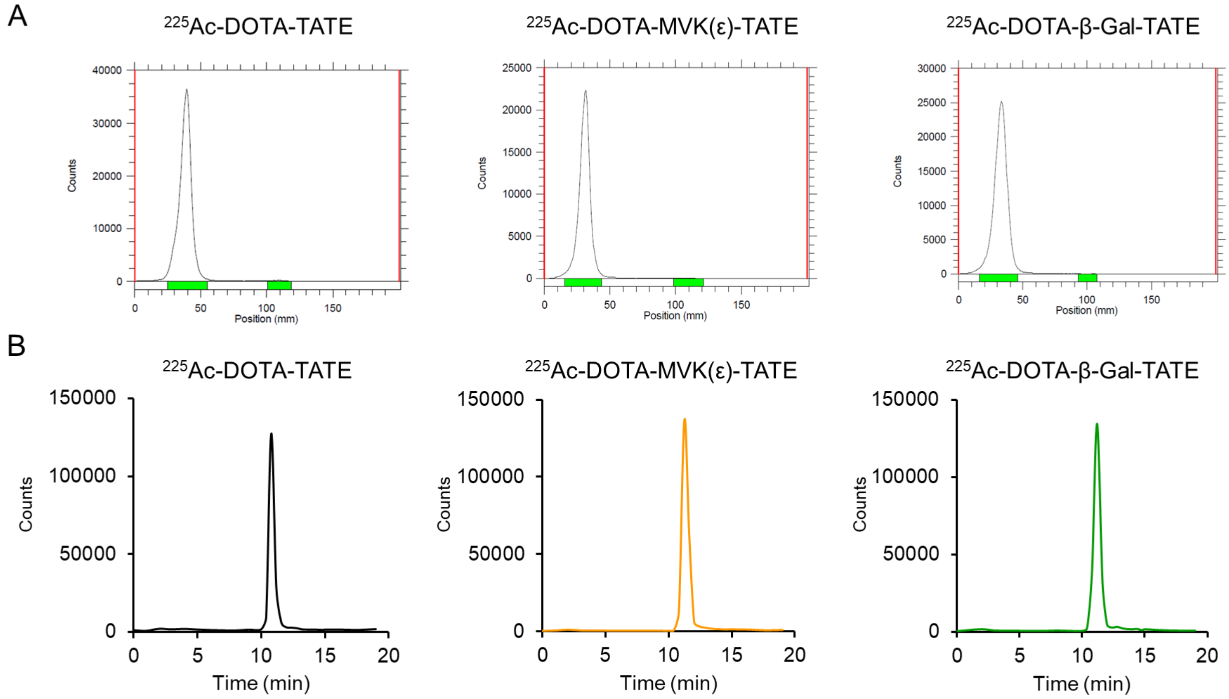


**Figure S17**. Radio-TLC and radio-HPLC of $^{225}$Ac-labeled DOTA-TATE derivatives. (**A**) Radio-TLC was performed on ITLC-SA strips using 50 mM EDTA or 100 mM $NH_4OAc$ as the mobile phase. (**B**) Radio-HPLC analysis (Agilent 1260 Infinity II, TSK-Octadecyl-4PW column) was carried out with a water/acetonitrile gradient containing 0.1% trifluoroacetic acid. For $^{225}$Ac analysis, fractions were collected every 0.4 min after sample injection, allowed to equilibrate for decay product ingrowth, and the radioactivity in each fraction was quantified using a gamma counter. All derivatives exhibited a single radioactive peak in the radio-HPLC chromatogram, eluting between 11 and 12 min.

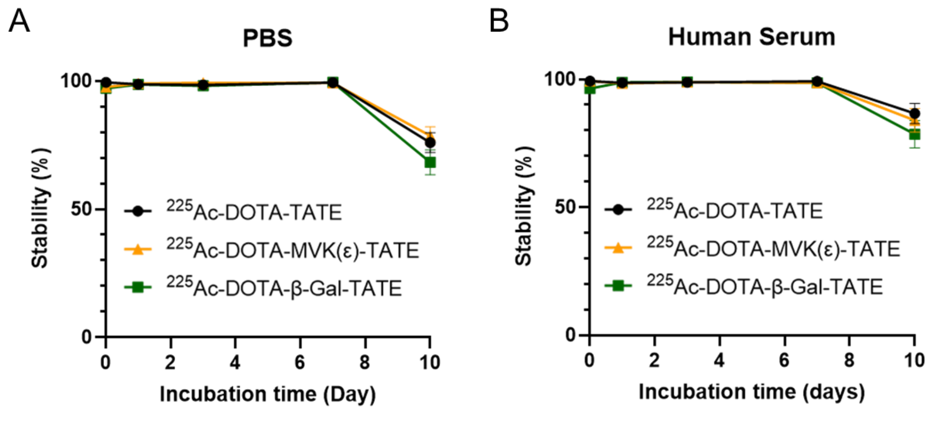


**Figure S18. Radiochemical stability of $^{225}$Ac-labeled DOTA-TATE variants over 10 days.** $^{225}$Ac-labeled DOTA-TATE variants were incubated in (A) phosphate-buffered saline (PBS) and (B) human serum at 37°C and analyzed by radio-TLC.

**Table S4. Biodistribution of $^{225}$Ac-labeled DOTA-TATE variants in AR42J tumor-bearing mice at 24 h and 4 days post-injection.** Radioactivity levels in tumors and major organs were measured at 24 h and 4 days following intravenous administration of $^{225}$Ac-labeled DOTA-TATE variants. Data are expressed as %IA/g (n = 4 - 5 per cohort).

| | $^{225}$Ac-DOTA-TATE | | | | $^{225}$Ac-DOTA-MVK(ε)-TATE | | | | $^{225}$Ac-DOTA-β-Gal-TATE | | | |
|---|---|---|---|---|---|---|---|---|---|---|---|---|
| Organs | 24 h | | 4 days | | 24 h | | 4 days | | 24 h | | 4 days | |
| | Average | SD | Average | SD | Average | SD | Average | SD | Average | SD | Average | SD |
| Blood | 0.03 | 0.01 | 0.04 | 0.02 | 0.03 | 0.01 | 0.03 | 0.02 | 0.04 | 0.03 | 0.03 | 0.00 |
| Tumor | 3.40 | 0.34 | 1.14 | 0.37 | 3.83 | 0.51 | 2.41 | 0.21 | 8.22 | 1.84 | 2.74 | 0.33 |
| Heart | 0.06 | 0.00 | 0.14 | 0.01 | 0.08 | 0.02 | 0.12 | 0.03 | 0.10 | 0.03 | 0.13 | 0.02 |
| Lung | 0.14 | 0.02 | 0.16 | 0.09 | 0.14 | 0.01 | 0.11 | 0.02 | 0.10 | 0.02 | 0.11 | 0.04 |
| Muscle | 0.10 | 0.01 | 0.16 | 0.04 | 0.09 | 0.01 | 0.15 | 0.01 | 0.10 | 0.03 | 0.22 | 0.07 |
| Femur | 0.20 | 0.02 | 0.41 | 0.11 | 0.33 | 0.07 | 0.32 | 0.04 | 0.48 | 0.11 | 0.42 | 0.13 |
| Spleen | 0.16 | 0.11 | 0.27 | 0.06 | 0.32 | 0.08 | 0.19 | 0.02 | 0.47 | 0.32 | 0.22 | 0.05 |
| Kidney | 4.19 | 0.28 | 1.02 | 0.13 | 1.27 | 0.25 | 0.43 | 0.05 | 0.89 | 0.08 | 0.28 | 0.04 |
| Liver | 0.29 | 0.02 | 0.65 | 0.14 | 0.61 | 0.06 | 0.42 | 0.06 | 0.89 | 0.49 | 0.50 | 0.03 |
| Intestine | 0.09 | 0.06 | 0.10 | 0.02 | 0.11 | 0.06 | 0.07 | 0.01 | 0.17 | 0.12 | 0.09 | 0.02 |

**Table S5**. Total Effective Dose per Injected Activity (mSv/MBq) of $^{225}$Ac-labeled DOTA-TATE variants in AR42J xenografts. The table summarizes the estimated total effective dose per unit of injected activity (mSv/MBq) for three $^{225}$Ac-labeled DOTA-TATE variants ($^{225}$Ac-DOTA-TATE, $^{225}$Ac-DOTA-MVK(ε)-TATE, and $^{225}$Ac-DOTA-β-Gal-TATE) based on biodistribution data obtained from AR42J xenografts. Values represent mean ± SD (n = 4 - 5).

| | $^{225}$Ac-DOTA-TATE | | $^{225}$Ac-DOTA-MVK(ε)-TATE | | $^{225}$Ac-DOTA-β-Gal-TATE | |
|---|---|---|---|---|---|---|
| Organ | Average (mSv/MBq) | SD (mSv/MBq) | Average (mSv/MBq) | SD (mSv/MBq) | Average (mSv/MBq) | SD (mSv/MBq) |
| Blood | 4.00E+03 | 4.15E+02 | 3.44E+03 | 3.24E+02 | 3.93E+03 | 7.18E+02 |
| Tumor | 4.30E+05 | 3.66E+04 | 6.64E+05 | 4.63E+04 | 1.06E+06 | 7.92E+04 |
| Heart | 1.42E+04 | 1.01E+03 | 1.06E+04 | 1.70E+03 | 1.20E+04 | 1.29E+03 |
| Lung | 3.85E+04 | 2.29E+03 | 1.95E+04 | 3.97E+03 | 1.91E+04 | 2.66E+03 |
| Muscle | 9.33E+03 | 9.30E+02 | 7.16E+03 | 6.37E+02 | 1.02E+04 | 2.38E+03 |
| Femur | 5.86E+04 | 1.72E+04 | 4.19E+04 | 6.69E+03 | 7.96E+04 | 1.04E+04 |
| Spleen | 3.41E+04 | 4.60E+03 | 2.55E+04 | 2.06E+03 | 3.37E+04 | 6.00E+03 |
| Kidney | 5.27E+05 | 2.40E+04 | 1.85E+05 | 9.85E+03 | 1.56E+05 | 6.61E+03 |
| Liver | 6.47E+04 | 9.91E+03 | 1.02E+05 | 3.55E+03 | 1.34E+05 | 5.92E+03 |
| Intestine | 3.04E+04 | 2.82E+03 | 2.36E+04 | 1.51E+03 | 2.87E+04 | 3.85E+03 |

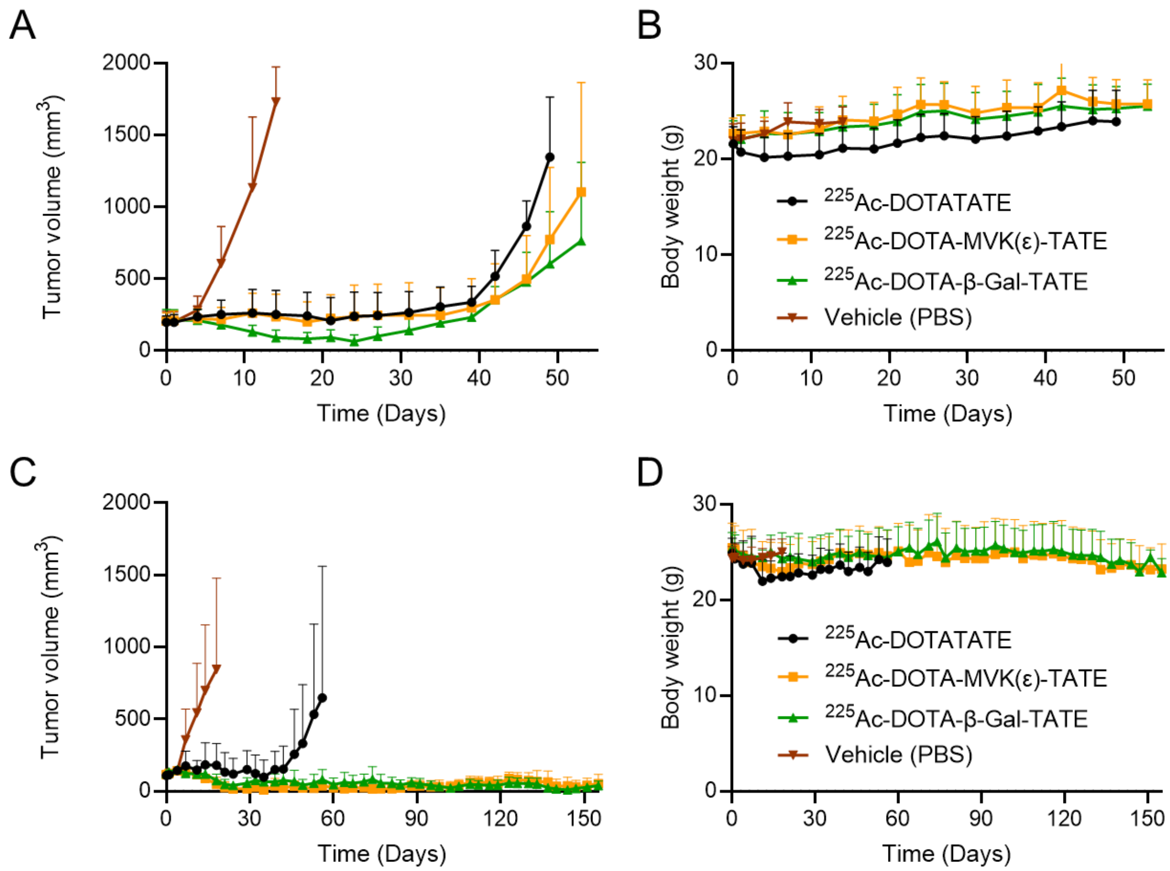


**Figure S19. Mean tumor volume and body weight changes following one- and two-cycles $^{225}$Ac-labeled DOTA-TATE variant treatments in AR42J tumor-bearing mice.** Data represent group means corresponding to the individual mouse data shown in the main text figures. Plots for each dataset are discontinued after first mouse death due to excessive tumor volume. Mean tumor volume (**A**) and mean body weight (**B**) in mice treated with a one-cycle regimen (148 kBq) of $^{225}$Ac-labeled DOTA-TATE variants or vehicle control (n = 8 per cohort). Mean tumor volume (**C**) and mean body weight (**D**) in mice receiving a two-cycle regimen (2 × 148 kBq, 8-day interval) of each agent or vehicle control (n = 6 per cohort). Data is presented as mean ± SD.

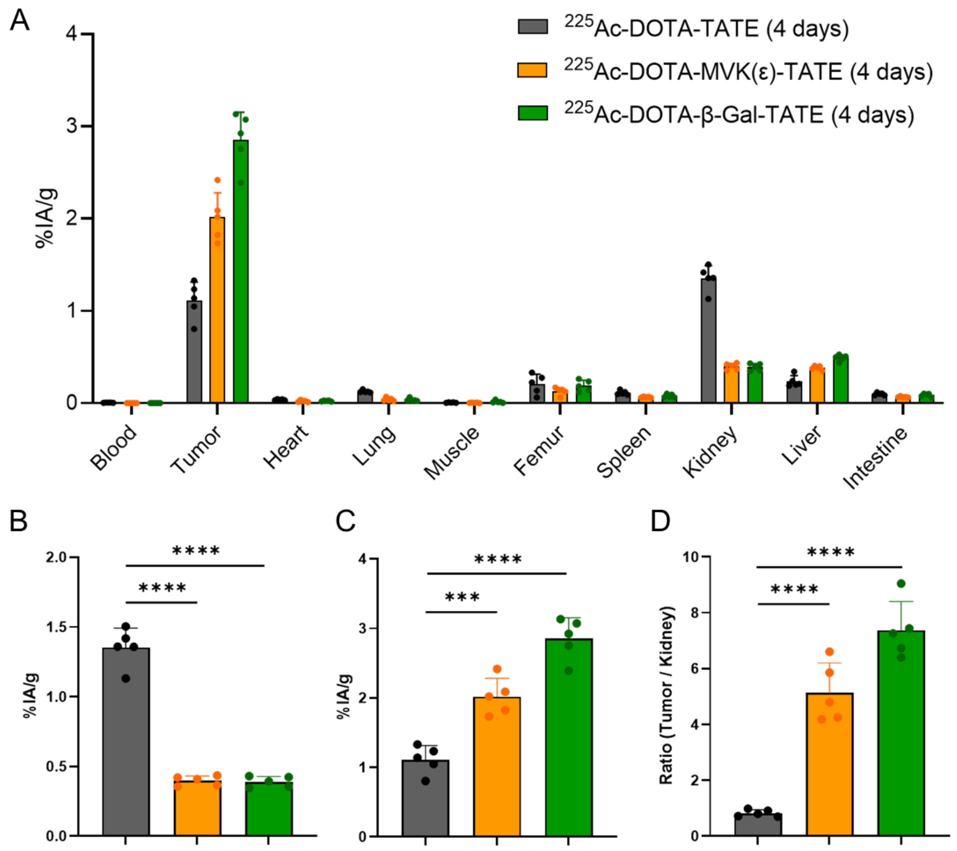


**Figure S20. Biodistribution of $^{225}$Ac-labeled DOTA-TATE variants in AR42J tumor-bearing mice at 4 days post-injection.** (**A**) Whole-organ biodistribution profiles. (**B**) Kidney uptake across the three groups. (**C**) Tumor uptake across the three groups. (**D**) Tumor-to-kidney uptake ratios for each group. Data are presented as mean ± SD (n = 4 per cohort). Statistical significance was assessed using unpaired two-tailed t-tests: ***$p < 0.001$, ****$p < 0.0001$.

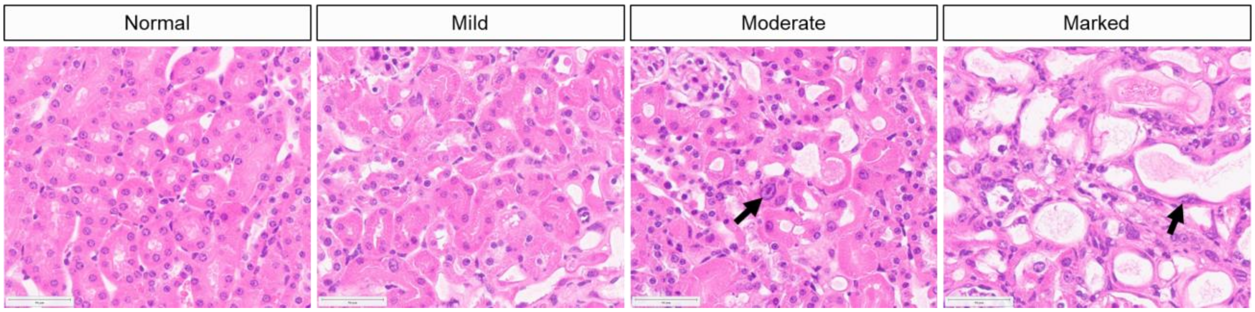


**Figure S21**. **Renal tubular changes from double-dose $^{225}$Ac-labeled DOTA-TATE variant treated mice.** Changes include renal tubular degeneration (cell swelling, cytoplasmic vacuolation) and renal tubular necrosis (nuclear pyknosis and karyorrhexis, cellular sloughing); markedly affected sections often had significant loss of cells and dilated tubular lumen. Karyomegaly (arrows) was observed in the renal cortical tubules, characterized by marked enlargement of nuclei with open chromatin and distinct nucleoli.

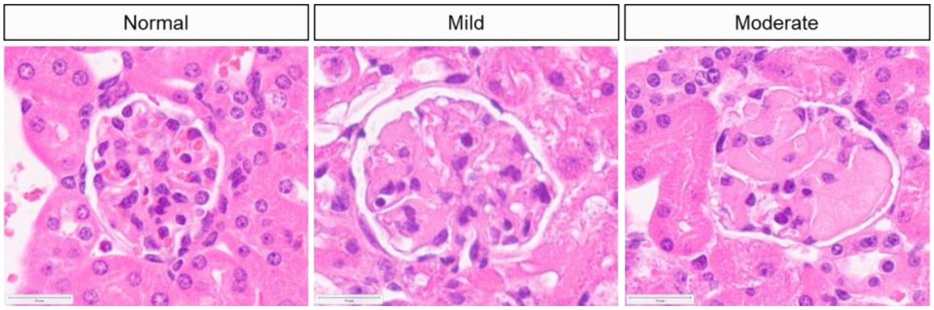


**Figure S22**. **Glomerular changes from double-dose $^{225}$Ac-labeled DOTA-TATE variant treated mice.** Glomerulopathy was observed characterized by increased mesangial matrix often manifesting as segmental eosinophilic deposits. Kidneys were graded based on the % of glomeruli affected and the extent of glomerular effacement. Changes were observed in all groups, including 1 control animal; however, these changes were increased in severity for $^{225}$Ac-DOTA-TATE group.

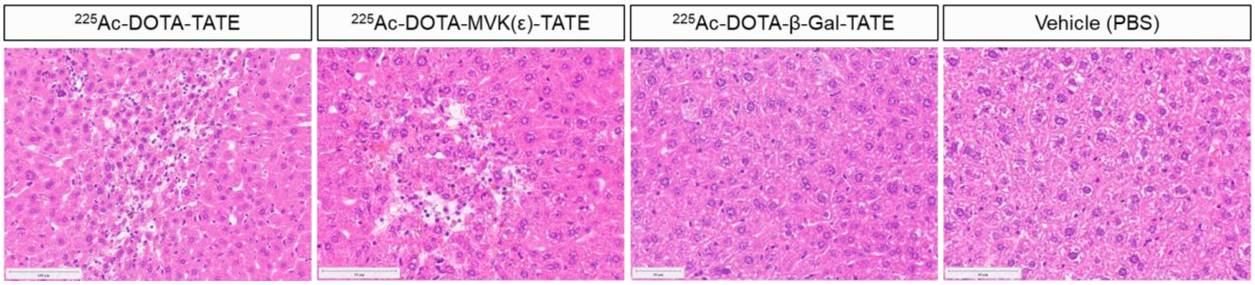


**Figure S23**. **Representative liver histopathology from double-dose $^{225}$Ac-labeled DOTA-TATE variant treated mice.** In individual mice (1 of 6 in $^{225}$Ac-DOTA-TATE and $^{225}$Ac-DOTA-MVK(ε)-TATE group), mild, focal hepatocyte injury was observed. The significance of this finding is unclear. Extramedullary hematopoiesis (not shown) was observed in liver sections from all groups.

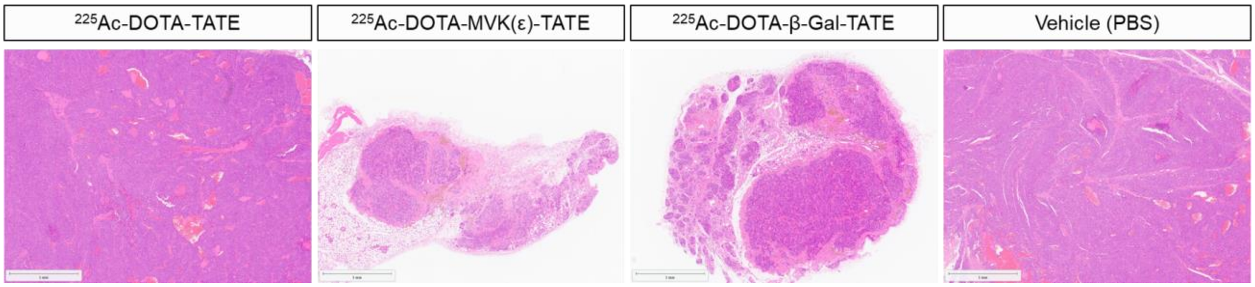


**Figure S24**. **Representative tumor sections histopathology from double-dose $^{225}$Ac-labeled DOTA-TATE variant treated mice.** Tumors in $^{225}$Ac-DOTA-MVK(ε)-TATE and $^{225}$Ac-DOTA-β-Gal-TATE group were significantly smaller.

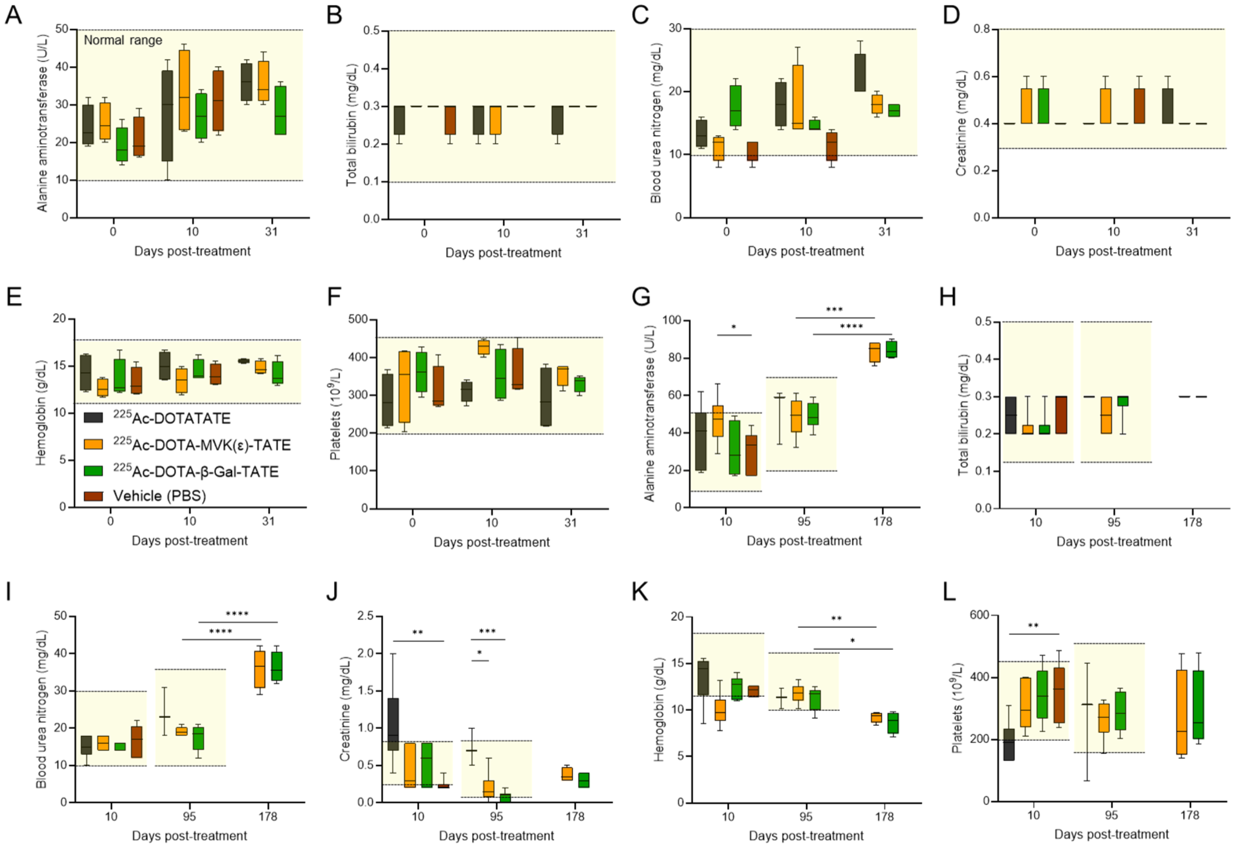


**Figure S25. Hematological and biochemical parameters following single dose and two cycle regimen with $^{225}$Ac-labeled DOTA-TATE variant treatments in AR42J tumor-bearing mice.** Blood samples were collected retro-orbitally at 10 days, 3 months (95 days), and 6 months (178 days) post-treatment and analyzed for basic metabolic panel (BMP) and complete blood count (CBC) parameters. (**A**–**F**) show results from the single-dose regimen (148 kBq), and (**G**–**L**) show results from the double-dose regimen (2 × 148 kBq, 8-day interval). Parameters measured included (**A**, **G**) alanine aminotransferase (ALT), (**B**, **H**) total bilirubin (TBIL), (**C**, **I**) blood urea nitrogen (BUN), (**D**, **J**) creatinine (CRE), (**E**, **K**) hemoglobin (HGB), (**F**, **L**) platelets (PLT). Shaded areas indicate reference ranges. Data are presented as mean ± SD (n = 4 or 6 per cohort). Statistical significance was defined as *$p < 0.05$, **$p < 0.01$, ***$p < 0.001$, and ****$p < 0.0001$.
